# LipoTag: A minimal motif for live and functional imaging of plant cell membranes

**DOI:** 10.64898/2026.04.08.717154

**Authors:** Maarten Besten, Tanguy Heesemans, Rik Froeling, Marie Zilliox, Youri Peeters, Robin Romein, Hui Tian, Jasper Lamers, Daan Vorselen, Bénédicte Charrier, Jan Willem Borst, Joris Sprakel

## Abstract

The plant plasma membrane is a highly dynamic structure that is crucial for cell compartmentalization, the maintenance of (bio)chemical gradients, signaling and cell growth and responses to stress. In plants, plasma membranes are tightly connected to the cell walls that encase them. These cell walls can act as diffusion barriers and prevent the use of a wide range of synthetic fluorescent probes that have been developed to study animal cell membranes, which lack a cell wall, with live functional imaging. Here, we introduce LipoTag, a minimal chemical motif that, upon chemical conjugation, transforms hydrophobic fluorophores into water-soluble, membrane-targeted probes that can permeate plant cell walls to reach their intended location. LipoTag uses a localized positive charge in combination with a short aliphatic spacer to direct cargo to the plasma membrane. We used LipoTag to develop a suite of membrane-specific fluorescent probes that work in walled organisms beyond the plant kingdom. In addition, we used LipoTag to develop functional reporters for the quantitative imaging of membrane density, lipid order and membrane oxidation in living plant tissues. LipoTag forms a modular platform for exploring the plant plasma membrane with a suite of contemporary imaging modalities.

## Introduction

All cells are surrounded and compartmentalized by a lipid bilayer that forms the cell membrane. These complex and dynamic structures mainly consist of saturated and unsaturated phospholipids, glycolipids and sterols [1, 2]. The physical properties and biological function of a membrane are governed by their lipid and sterol composition as well as the presence of membrane-anchored or membrane-bound proteins, which in turn govern the spatial organization of the membrane and its mechanical and biochemical features [3–5]. Membranes act as selective barriers between the inside of the cell and its surroundings, to enable generating chemical gradients across the membrane, allow selective fluxes of bio(chemical) substances across the membrane and as a crucial signaling hub through the presence of a wide array of cell surface receptors and accessory proteins; through these functions, membranes play a crucial role in virtually all aspects of cellular physiology and its responses to stresses [6–10].

Membranes in organisms that are surrounded by a cell wall, such as those in plants, are subjected to rather extreme mechanical conditions. Plant membranes have to withstand the extreme and fluctuating internal turgor pressures of the cell, while being pressed onto the plant cell wall that provides a mechanical reactionary force to create a state of mechanical homeostasis [11–13]. Moreover, plant cell membranes are home to a vast number of receptor-like kinases that perceive signals from the cell exterior and relay signals to the inside of the cell to activate downstream signaling processes. Cell surface signaling often involves highly dynamic and adaptive responses of the plasma membrane and its proteome for example the transient formation of membrane nano- and microdomains [14–16]. Given their critical role in cell homeostasis and mechanostasis, and in cell signaling, plant membranes have been the subject of extensive study [17–20]. Surprisingly however, the toolbox for studying plant membranes with fluorescence microscopy using synthetic fluorescent probes is very limited [21]. This is primarily caused by the fact that any probe that must make its way into the hydrophobic plasma membrane must first diffuse through, and not become bound, by the complex plant cell wall [22, 23].

Despite a lack of tools, fluorescence microscopy is an invaluable tool for studying cell membranes, and has been successful in making substantial advances in membrane biology in other kingdoms. Fluorescence microscopy of membranes requires membrane-specific fluorescent probes, which can be genetically encoded (fluorescent protein), based on antibody-antigen recognition, use metabolic labeling or use fully synthetic small molecule probes [24–28]. In mammalian cell biology, the toolbox of available antibodies and synthetic probes is extensive, and developing stable fluorescent marker lines in cell cultures requires typically only a few months depending on the cell type [29, 30]. By contrast, for plants, tools are much more limited. There are only a handful of commercially available synthetic fluorescent probes, such as the FM family, that properly function to reveal membrane localization but have broad excitation and emission spectra, and no known antibodies for live plant cell membrane imaging [31, 32]. Furthermore, generating stable fluorescent marker lines is often very time consuming for most plant species, while a large portion of plants are not genetically accessible.

Current efforts to develop bespoke synthetic membrane probes for plants are limited and often do not include imaging in tissues a couple of cell layers thick [33–35]. Alternatives such as metabolic labeling have been explored and show promising results in tracking lipids and membranes, although this method requires fixation and thus is not compatible with live imaging [36]. Because plants are surrounded by a cell wall, which is best described as a hydrophilic polysaccharide hydrogel matrix, the requirements for a synthetic fluorescent probe that performs well in plants will differ significantly from a probe that works in mammalian cells [37]. In 2D mammalian cell cultures, synthetic probes do not have to cross any physical barriers, which allows them to be highly hydrophobic and almost instantaneously insert into membranes. However, in plants, membrane probes must cross multiple barriers, such as the hydrophobic cuticle and the hydrophilic cell wall, with an acidic pH before inserting into the membrane. This requires synthetic plant membrane probes to be more hydrophilic and pH stable compared to their mammalian counterparts.

Here we describe the development and application of a toolbox specifically designed for, but not limited to, live and functional fluorescence imaging of plant cell membranes. This system is based on a minimal chemical motif, which we named LipoTag, that, when conjugated to a hydrophobic synthetic fluorophore, inserts spontaneously and rapidly into plant cell membranes. LipoTag exhibits minimal toxicity at useful concentrations and shows rapid permeation through plant cell walls to ensure deep tissue penetration. LipoTag was tested in a wide range of organisms and shown to function well in a range of different plant species as well as fungi, bacteria and mammalian cells. We used LipoTag to develop a range of cell membrane probes based on both commercially available and custom-made probes across the visible spectrum, making them highly suitable for multiplexing experiments. Combining LipoTag with functional probes, that reveal physico-chemical features in their environment, we constructed a library of fluorophores that allow for imaging cell membrane fluidity, lipid order and oxidation.

## Results

### LipoTag design

The design of LipoTag was guided by taking inspiration from known probes that work well in plants, as well as informed by the constraints imposed by plant anatomy. As plants cells are encased by a hydrophilic cell wall and plant organs are separated from the outside by a hydrophobic wax-like cuticle, a suitable probe must be able to cross both a very hydrophobic layer as well as permeate through a very hydrophilic structure, ideally without requiring chemical additives, such as DMSO, to accomplish this. This thus requires a delicate balance between hydrophilicity and hydrophobicity in the probe design [22]. Commercially available probes that perform well in plants such as dyes from the FM family (FM1-43, FM2-10, FM4-64, FM5-95) and Di-4-ANEPPDHQ as well as the custom made N+-BDP all have 1 shared molecular characteristic: the presence of two quaternary nitrogen atoms resulting in a net charge of +2 [31, 32, 35, 38, 39]. These quaternary nitrogen atoms, which are in close proximity to each-other, and provide a very hydrophilic center, are combined with a molecule that is otherwise highly hydrophobic; resulting in an amphiphilic molecule that can interact with membranes both by hydrophobic interactions as well as electrostatic bonding with negatively charged phospholipids. Moreover, in our previous work on cell wall targeting, we noted that cell wall transport of a fluorescent probe is highly sensitive to the sign of the charge, where negative charges appear to inhibit cell wall diffusion [22]. Even though many of the synthetic membrane probes that have been developed for mammalian cells also have a positive charge and an amphiphilic character, the majority these synthetic probes do not perform well in plants. For example, two widely used fluorophores in mammalian cell membrane imaging, DiA and DiL, each possess a single positive charge, which is part of a conjugated system, with the rest of the molecule being highly hydrophobic and comparatively large. While these probes effectively stain mammalian cell membranes, we found these to perform poorly in plants, resulting in little-to-no staining (Fig. S1). By contrast, FM1-43 and FM-64 possess 2 positive charges and have a significantly smaller hydrophobic region. These probes stain cell membranes in plants very well, for example, permeating deeply into the roots of *Arabidopsis thaliana* seedlings (Fig. S1-3). From this, we hypothesized that membrane probes for plants, need an amphiphilic balance that is strongly shifted to the hydrophilic side, as compared to those for animal cells, due to the presence of the hydrophilic cell wall as an additional diffusion barrier.

Hence, we set out to develop a modular motif with a hydrophilic head group and a modest hydrophobic spacer, that can be conjugated to a moderately small hydrophobic fluorophore to meet these design requirements. Starting from commercially available 1-bromo-6-chlorohexane we synthesized, in 4 steps without the need of chromatography purification, the structure we name LipoTag: a linear molecule bearing 2 positive charges at one end and an azide group at the other end (Fig. 1a top). This terminal azide group enables the conjugation to fluorophores equipped with an alkyne through click-chemistry (Fig. 1a bottom). Click chemistry is the umbrella term for a set of chemical reactions that occur with high specificity, under mild conditions and which produces no or minimal side products [40].

**Figure 1:**
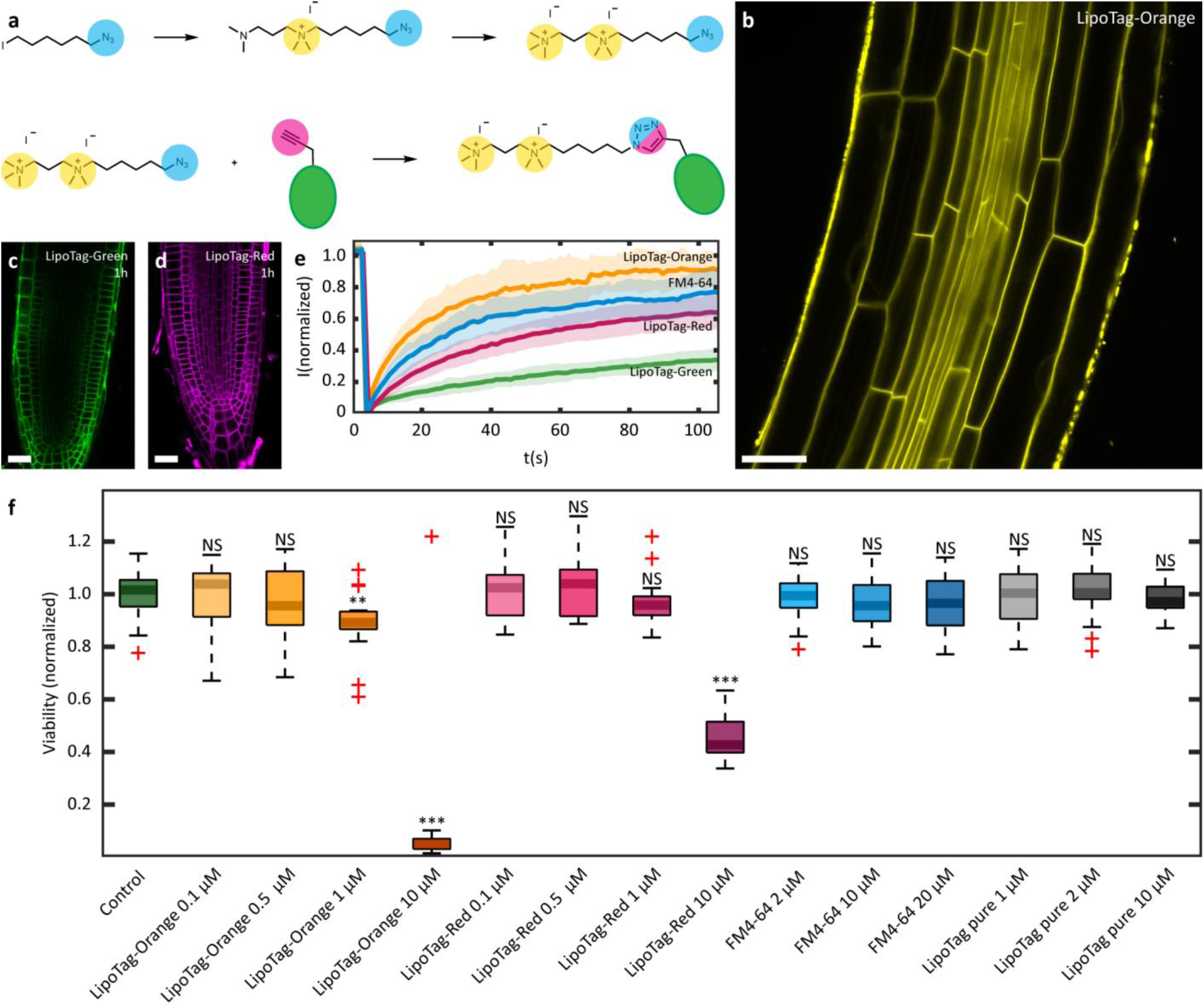
Lipotag synthesis, membrane dynamics and toxicity. **a**, Synthesis steps to make LipoTag (top) and LipoTag modified dyes (bottom). **b**, Arabidopsis root stained with LipoTag BDP-TR. **c-d**, Arabidopsis roots stained with LipoTag BDP (**c**) and LipoTag BDP-630 (**d**) for 1h. **e**, FRAP curve for LipoTag BDP (green), LipoTag BDP-TR (yellow), LipoTag BDP-630 (magenta) and FM4-64 (cyan). The shaded area represents the standard deviation. **f**, Protoplast assay toxicity test of LipoTag modified dyes (LipoTag-BDP-TR and LipoTag-BDP-630), FM4-64 and non-modified LipoTag at various concentrations. In box plots the center line is the median, box bounds represent the 25th and 75th percentiles. Whiskers span from the smallest to the largest data points not considered outliers. Red + symbols indicate outliers. \*\**P* < 0.01; \*\*\**P* < 0.001; NS, *P* > 0.05; two-sided Wilcoxon rank sum test. Scale bars represent 25 µm

Next we conjugated the azide-functional LipoTag to the alkyn tag of a commercially available orange-emitting BODIPY fluorophore yielding the probe: LipoTag-Orange. Staining of the *Arabidopsis thaliana* root with LipoTag-Orange for 30 minutes shows clear staining of the cell membrane up to the vasculature tissue (Fig. 1b). By contrast, incubating Arabidopsis roots with the orange BODIPY probe without the LipoTag motif did not result in any detectable staining of the membrane (Fig. S1, confirming that LipoTag is operational in the membrane targeting. We followed an identical strategy to create a green (LipoTag-Green) and far-red (LipoTag-Red) fluorescent membrane probe. We found the optimal staining concentration to be ∼0.5 μM for all 3 probes (Fig. S4). However, when determining the time required for staining Arabidopsis seedling roots, we noticed that LipoTag-Green penetrates the multicellular tissues noticeably slower in roots compared to LipoTag-Orange and LipoTag-Red (Fig. 1c-d, Fig. S5-7). This is evidenced by a lack of fluorescence in the root tip in plants stained with LipoTag-Green compared to tissue stained with LipoTag-Orange and LipoTag-Red with the same concentration and incubation time (Fig. S4).

To further investigate this, we performed Fluorescence Recovery After Photobleaching (FRAP) experiments on all 3 LipoTag probes (LipoTag-Green, LipoTag-Orange and LipoTag-Red) and using the current state-of-the-art plant membrane probe FM4-64 as a control. Interestingly, while both LipoTag-Orange and LipoTag-Red exhibit similar recovery times compared to FM4-64, LipoTag-Green fluorescence recovers significantly slower, indicative of hindered diffusion (Fig. 1e). This result aligns with the prolonged required staining time as noted above. The molecular weights of all four probes do not differ substantially; instead, we noted a key structural distinction. LipoTag-Green is the only probe in this set where the LipoTag motif and fluorescent core are oriented perpendicularly with respect to each-other, forming a T-shaped molecule. By contrast, the other three probes all have a linear geometry (Fig. S8) [31]. This suggests that when developing membrane probes the number of charges and the size of the hydrophobic region determine whether or not plant membranes can be stained but their relative orientation determines their mobility within a membrane and ultimately plant tissue.

The membrane specificity of all 3 LipoTag dyes was confirmed by performing a plasmolysis experiment on LipoTag dye stained seedlings. Upon plasmolysis, the fluorescence signal remained primarily localized to the retracted plasma membrane. However, as some signal appeared to originate from the cell wall, we looked in more detail and found that the probe also retains in both Hechtian strands and the Hechtian reticulum. Counterstaining with cell wall specific CarboTag dyes during plasmolysis confirmed there is no staining of the cell wall by LipoTag dyes (Fig. S9). LipoTag thus results in specific plasma membrane localization in Arabidopsis seedlings.

Given that the amphiphilic structure of membrane-targeting dyes may disrupt membrane packing and integrity, we determined their toxicity. This was done in a protoplast viability assay, since protoplasts, plant cells stripped from their cell wall, are highly sensitive to membrane stability. The two tested LipoTag dye conjugates LipoTag-Orange and LipoTag-Red did not affect cell viability at concentrations of 0.1 and 0.5 μM while a concentration of 1 μM only induced a minor decrease in viability for LipoTag-Orange and did not result in a significant decrease in viability for LipoTag-Red (Fig. 1f). Only after increasing the concentration to 10 μM a sharp decrease in viability is observed. FM4-64, which also has an amphiphilic structure, did not exhibit any toxic effects between 2 and 20 μM. Interestingly, the LipoTag motif alone did not show a decrease in viability between 1 and 10 μM, indicating that the toxic effects observed at higher LipoTag-Orange and LipoTag-Red concentrations originate from the probe-LipoTag combination, which we also noted in our earlier work on cell wall probes [22]. It should be noted that toxic effects in protoplasts might be exaggerated by their fragility since their cell wall has been stripped away. Additionally, lower cell viability might not be the only effect of a potentially disrupting compound, FM4-64 for example exhibits no toxic effects in protoplasts but has been reported to alter the localization of PIN proteins. ([31])

#### Applicability

In the FM dye family the amphiphilic structure is partially encoded in their fluorescent core. As this family of dyes is characterized by broad excitation and emission spectra, and their targeting and fluorescent properties are intertwined there is no room for engineering their fluorescent properties. This results in limited spectral flexibility when choosing a synthetic membrane probe, complicating multiplexing experiments with other synthetic fluorophores or genetically encoded reporters. LipoTag can transform hydrophobic, alkyne-modified fluorophores into plant cell membrane stains. The increased use of fluorescence techniques in research has resulted in an enormous diversity in fluorophores with various modifications, including alkyne-modified fluorophores that can be commercially procured. To demonstrate the flexibility of LipoTag we synthesized LipoTag probes ranging from green to deep red. These fluorophore-LipoTag fusions all stain cell membranes in Arabidopsis root cells with no observed binding to the adjacent cell wall (Fig. 2a-b,Fig. S9). This raises the question on the generality of our approach. We also procured alkyne-modified Cy5 and synthesized a hydrophobic, alkyne-modified oxazine dye. Upon conjugation with LipoTag the resulting probes did exhibit excellent penetration in roots and displayed high levels of cell internalization (Fig. S10). We suspect this rapid internalization is due to the presence of a delocalized positive charge in both dyes, causing membrane destabilization upon insertion, which could lead to increased endocytosis depending on the membrane potential [41]. Hence, while LipoTag works as an efficient targeting motif for some fluorophores, each specific LipoTag-fluorophore pair needs to be subjected to compatibility as the fluorophore has a strong impact on the probe behavior.

**Figure 2:**
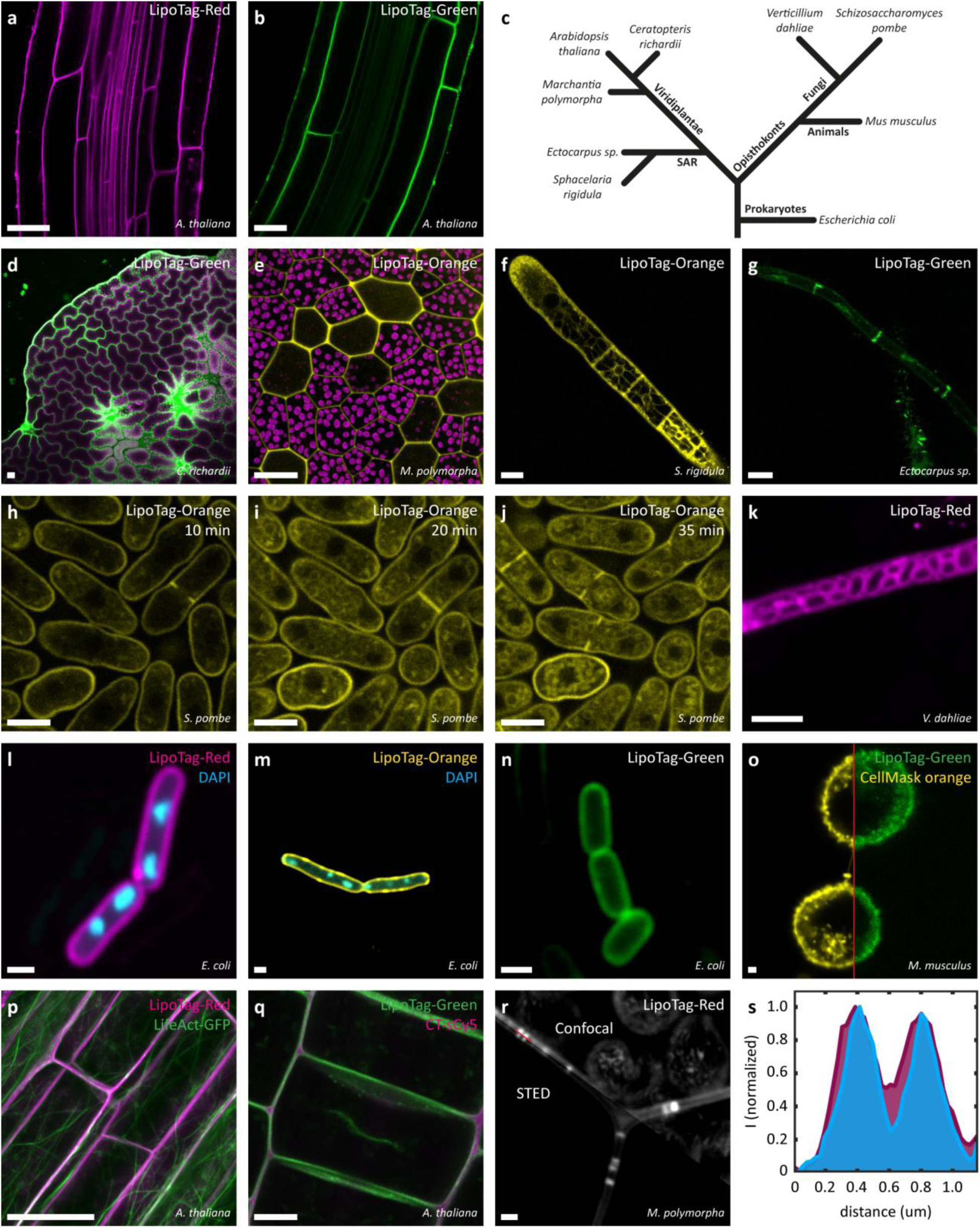
Performance of LipoTag in various organisms, multicolor imaging and super-resolution imaging. **a,b,** Arabidopsis roots stained with LipoTag-Red (**a**) and LipoTag-Green (**b**). **c,** phylogenetic tree of organisms used in this research. **d,** Gametophyte of the fern *C. richardii* stained with LipoTag-Green, chloroplasts autofluorescence in magenta. **e,** Gemma of the liverwort *M. polymorpha* stained with LipoTag-Orange, chloroplasts autofluorescence in magenta. **f,g,** staining of the brown algae *S. rigidula* (**f**) and *Ectocarpus sp.* (**g**) with LipoTag-Orange and LipoTag-Green respectively. **h-j,** growing and dividing *S. pombe* stained with LipoTag-Orange after 10 minutes (**h**), 20 minutes (**i**) and 35 minutes (**j**). **k,** *V. dahliae* stained with LipoTag-Red. **l-n,** *E. coli* stained with LipoTag-Red (**l**), LipoTag-Orange (**m**) and LipoTag-Green (**n**). dsDNA stained with DAPI is shown in cyan (**l,m**). **o,** Murine macrophage-like J774 cells co-stained with CellMask Orange (yellow) and LipoTag-Green (green). **p,** LipoTag-Red (magenta) with actin marker LifeAct-GFP (green) in Arabidopsis root tissue. **q,** Arabidopsis root co-stained with LipoTag-Green (green) and CarboTag-sCy5 (magenta) to targeting the cell membrane and cell wall respectively. **r,s,** STED vs confocal imaging of plasmodesmata in Marchantia gemmae with LipoTag-Red (**r**) and a line plot of the normalized STED (cyan) and confocal (magenta) signal (**s**). Scale bars represent 25 µm (**a, b, d, e, p**), 5 µm (**f-k, o, q**) or 1 µm (**m-o, r**).

LipoTag is suitable for a range of species in the plant lineage. We successfully stained gemmae of the liverwort *Marchantia polymorpha* and gametophytes of the fern *Ceratopteris richardii* with LipoTag dyes (Fig. 2d-e). Because these organisms possess thick cell walls and cuticles compared to Arabidopsis, reliable staining was achieved after 2 hours of incubation, compared to the roughly 15-30 minutes required for Arabidopsis roots (Fig. S5-7) [42].

We then explored the suitability of LipoTag for membrane imaging in organisms outside of the plant kingdom (Fig. 2c). We began with several species of brown algae, which are phylogenetically distinct from plants and part of the Stramenopiles phylum. In *Sphacelaria rigidula,* LipoTag-Orange managed to stain the cell membrane and membranes of organelles (Fig. 2f).

However, not all LipoTag probes stained equally well, LipoTag-Green for example only managed to stain the membranes between cells of *Ectocarpus sp.* (Fig. 2g, Fig. S11-14). This may be attributed by brown algae possessing a dense alginate-rich cell wall that might be problematic to navigate for LipoTag dyes [43]. Another explanation is the high osmotic strength of Provasoli Enriched Seawater (PES) medium required for culturing brown algae that could lower the critical micelle formation, creating less mobile dye clusters in solution as the fluorophores resemble surfactants. Compared to novel bespoke brown algae probes such as styryl benzoindoleninium sulfonate (SBIS) LipoTag probes perform considerably less well, underscoring the possible role of differences in media, cell walls and possibly plasma membranes between brown algae and land plants [44].

We extended our imaging to micro-organisms. We imaged the filamentous fungus *Verticillium dahliae*, the yeast *Schizosaccharomyces pombe* and bacterium *Escherichia coli* with multiple LipoTag dyes. In both fungi, LipoTag dyes successfully stained the membrane but suffered from high endocytosis, which is consistent with the high membrane turnover in these actively dividing and growing cells. (Fig. 2h-k). In *S. pombe,* LipoTag-Orange appears to accumulate in small vesicles that are transported to the cell division machinery during cytokinesis; time-lapse imaging of dividing cells with this probe highlights the cytokinetic ring (Fig. 2h-j). Additionally, *E. coli* membranes are successfully stained with all 3 LipoTag dyes. LipoTag-Orange and LipoTag-Red were used for multiplexing in combination with the dsDNA stain DAPI (Fig. 2l-n).

Finally, we explored LipoTag’s compatibility with cells of the animal kingdom by imaging murine macrophages. As the membrane of macrophages generally have a high endocytosis rate as part of their role in the immune system, a high degree of internalization is observed [45]. Nevertheless, staining of the membrane with LipoTag-Green is observed which was confirmed by CellMask Orange counterstain,a commonly used membrane probe for mammalian cells (Fig. 2o). Incubation with LipoTag-Orange and LipoTag-Red did not result in satisfactory staining of the membrane but rather exhibited rapid internalization (Fig. S15). The slow diffusion of LipoTag-Green could be an advantage or even prerequisite in staining single cells, such as those in suspension cultures. These observations suggest that the reduced mobility of LipoTag-Green, identified in our FRAP experiments, may be an advantage or even a prerequisite for stable plasma membrane staining of cells that lack a cell wall.

Access to a range of cell membrane stains with different spectral properties facilitates multiplexed imaging, for example by combining with existing lines that express genetically-encoded fluorescent markers. We demonstrate this by combining LipoTag-Red with the actin marker LifeAct-GFP in *Arabidopsis thalian*a that allows for simultaneous imaging of the membrane and actin cytoskeleton (Fig. 2p). Multiplexing can also be achieved by combining multiple small molecule probes, such as LipoTag with our previously developed cell wall toolbox CarboTag. Although the physical distance between the cell membrane and cell wall is below the optical resolution limit of confocal microscopes at the right Z-plane the membrane and wall can be distinguished from each other. Moreover, also here, LipoTag acts as a vesicular marker seen by the accumulation of lipid material at a new cell plate in a dividing cell (Fig. 2q).

During our investigations of gemmae, the asexual propagules of the liverwort *Marchantia polymorpha*, we observed that LipoTag-Red revealed strongly fluorescent connections bridging adjacent cells through the thick cell walls. Counterstaining with aniline blue confirmed these structures as plasmodesmata (Fig. S16). These structures have been imaged and studied before in Marchantia, but to our knowledge, the membrane spanning across 2 cells has not previously been labeled with synthetic probes [46]. Since the diameter of these structures is well below the diffraction-limited resolution of regular confocal microscopes, we employed Stimulated Emission Depletion (STED) microscopy to increase the resolution [47]. LipoTag-Red was used because of its far-red emission, high photostability and quantum yield, which make it highly suitable for this technique [48]. With this probe, we were able to clearly stain plasmodesmata in Marchantia gemmae and improve its resolution by the use of STED microscopy (Fig. 2r). This improvement is evident in the increased sharpness of the intensity profile along the membrane as compared to that obtained from the conventional confocal image of the same sample (Fig. 2s).

### Imaging membrane fluidity: LipoTag-BDP

The cell membrane of plants is a complex and dynamic structure that is constantly adapting to meet changing demands, whether in developmental processes or upon encountering external stresses. Moreover, maintaining membrane integrity is fundamental to plant survival, as it must accommodate the rapid expansion, division, and structural remodeling required for growth. Consequently, characterizing the membrane’s physicochemical properties under live conditions can provide critical insights into the membranes function in these processes. To this end, in the following sections, we develop a set of LipoTag-based functional probes that can be used for the spatiotemporal mapping of key membrane features: fluidity, lipid order and oxidation.

We previously reported a membrane mechanoprobe based on a phenyl-BODIPY (BDP) molecular rotor probe coupled to an aliphatic carbon chain with 2 quaternary nitrogen atoms, similar to our LipoTag motif. This probe, N+-BDP was utilized to detect changes in membrane tension in osmotically stressed cells, across different organisms, tissue types and in mutants [17, 35, 49]. The BDP rotor undergoes an intermolecular rotation upon photoexcitation to a nonradiative, dark state. To make this rotation the fluid surrounding the molecule must flow, thereby coupling the molecular rotation to the viscous and hydrodynamic features of its surroundings [50]. The result is that this probe has a low fluorescence lifetime when BDP can rotate freely, such as in open spaces and low viscosity media, and in high lifetimes when the probe rotation is hindered either by the viscosity of the surrounding medium or by the presence of narrow porous matrices, such as cell walls. The coupling of fluorescent lifetime to its surroundings allows for using BDP rotors as probes to map the fluidity and tension of lipid membranes with fluorescence lifetime imaging (FLIM).

We conjugated a custom-made BDP rotor to LipoTag; the resulting probe, LipoTag-BDP, (Fig. 3a) exhibits green fluorescence (Fig. 3b) and clear cell membrane localization. Indeed, this molecule shows an increase in fluorescence lifetime with an increase viscosity of the medium in which it is dissolved (Fig. S17). To evaluate its applicability in measuring cell membrane tension we observed stomata guard cells upon an abscisic acid (ABA) treatment. ABA induces the closing of stomata by causing a loss of turgor, which lowers the outward pressure exerted on the membrane and causes surrounding cells to push inward, resulting in an altered membrane aperture compared to open stomata (Fig. 3c-d). We find that the membranes of guard cells of closed stomata show a significantly decreased fluorescence lifetime compared to opened stomata (Fig. 3c-e). A lower lifetime indicates a less sterically hindered intramolecular rotation, signifying a reduction in lipid packing density and an increase in membrane tension.

**Figure 3:**
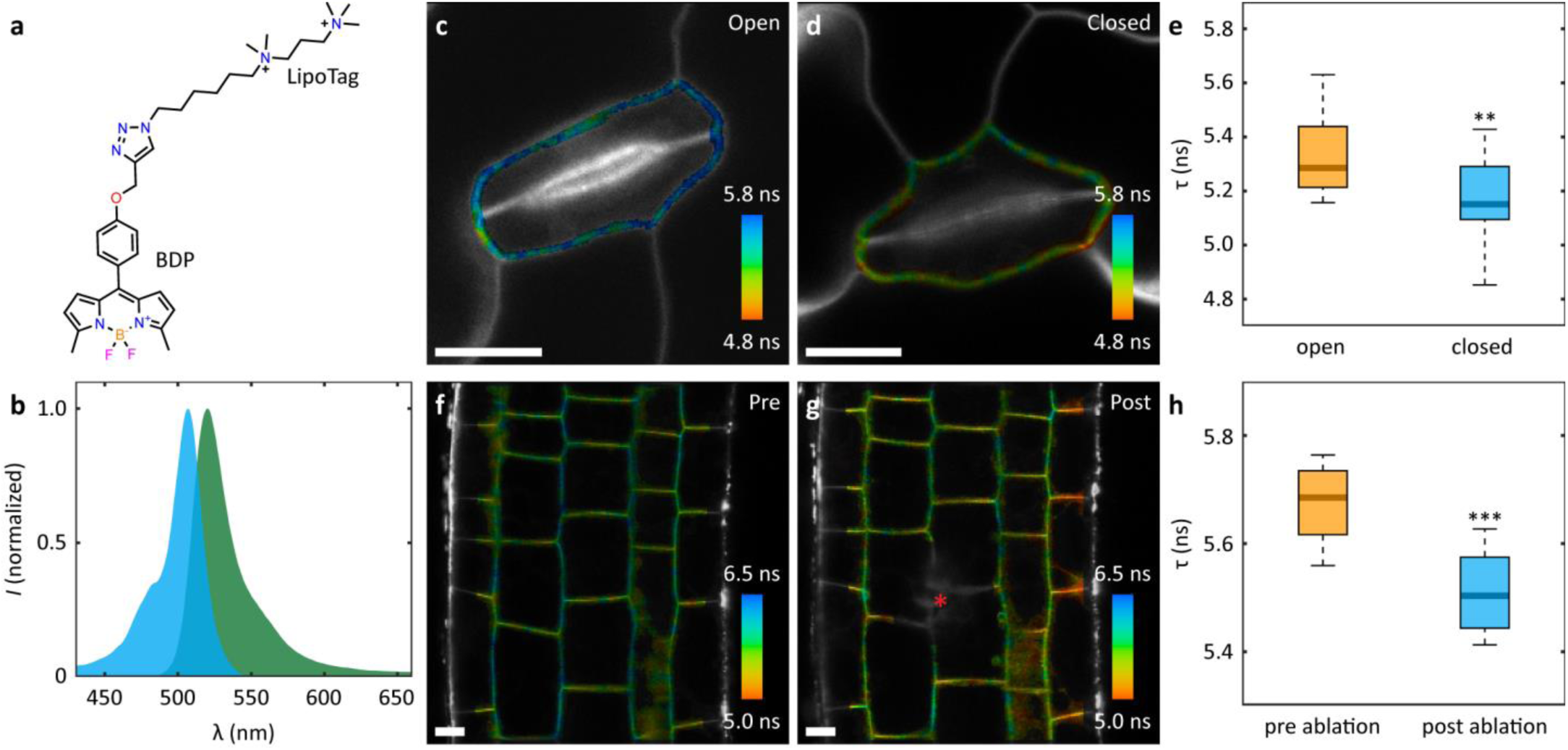
LipoTag-BDP based imaging of cell membrane density. **a,** Chemical structure of LipoTag-BDP density probe. **b,** Normalized excitation (cyan) and emission (green) spectra of LipoTag-BDP. **c,d,** FLIM images of opened (**c**) and closed (**d**) stomata in Arabidopsis leaves. **e,** Comparison of fluorescence lifetimes between opened and closed guard cell outer membranes. **f,g,** FLIM images of Arabidopsis root before (**f**) and after (**g**) ablation, ablation site is indicated by red asterisk. **h,** Comparison of fluorescence lifetimes of membranes before and after ablation. In box plots the center line is the median, box bounds represent the 25th and 75th percentiles. Whiskers span from the smallest to the largest data points not considered outliers. \*\**P* < 0.01; \*\*\**P* < 0.001; two-sided Wilcoxon rank sum test. Scale bars represent 10 µm. Color scale in FLIM images corresponds to the fluorescence lifetime.

The mechanics of a plant tissue are governed by a tissue-wide balance between tensile and compressive forces created by turgescent cells pushing and pulling on their neighbors. Deflating a single cell, by laser ablation, disturbs this balance and creates a mechanical imbalance in the tissue leading to rapid changes in tension across load bearing structures including the cell membranes. The LipoTag-BDP membrane tension probe can reveal these changes. Laser ablation of a single cell, leads to a fast decrease in fluorescence lifetime in the surrounding tissue, indicating a rapid change in tension patterns (Fig. 3f-h). The physical damage of ablation generates auto-fluorescent compounds with short lifetimes; hence, the membranes directly contacting the ablation site were omitted from the analysis to prevent contamination of the FLIM signal. The abrupt, localized loss of turgor at the ablation site causes neighboring epidermal cells to deform towards the ablation site which is accompanied with the inner intact cell layers pushing outward [51]. This process is comparable to the closing of stomata where changes in turgor change the cell morphology and increase the surface area of the membrane, resulting in a decrease in lifetime, which signal an increase in membrane tension.

### Imaging lipid order – LipoTag-NR

Biological membranes are heterogenous structures with a mosaic-like structure of highly ordered regions containing protein complexes and lower order structures; often envisioned as a liquid sea of disordered lipids containing floating solid domains with a higher degree of order [52–54]. In plants, these highly ordered regions, also called lipid rafts or micro- and nanodomains, are likely involved in the polar localization of various transmembrane and membrane-associated proteins and play a key role in various signaling processes [15, 16, 55–60]. In turn, membrane order is sensitive to the local sterol content of the membrane as sterols can alter the chemical polarity of the membrane interior [61, 62].

Currently, the two main strategies to visualize membrane organization and chemical polarity involve tracking fluorescently labelled membrane proteins such as flotillins or remorins or the use of solvatochromic probes like di-4-ANNEPS and Laurdan [38, 63–67]. While the localization of nanodomain-associated proteins can serve as a proxy for the localization of membrane domains, it does not provide direct biophysical data regarding the bilayer itself. By contrast, solvatochromic probes provide real-time information on membrane order through environment-dependent shifts in their excitation or emission spectra. The two probes that have been employed in plants, Laurdan and di-4-ANNEPS, both have a blue shifted emission spectrum when incorporated in an order lipid environment, making them useful as ratiometric probes. Like most other probes that are well established in mammalian cells their penetration in live plants is limited.

To overcome this limitation, we first tested the ratiometric Nile Red based membrane order probe NR12S [68]. It exhibits a characteristic blue-shifted emission in ordered lipid domains and has been successfully employed to map membrane organization in mammalian cells and the walled oomycete *Phytophthora infestans* [68, 69]. Unfortunately using NR12S in Arabidopsis roots this did not result in satisfactory staining of the membrane which can likely be attributed to its high hydrophobicity caused by its long alkyl chain. The narrower excitation and emission spectra of Nile Red and its chemical flexibility compared to Laurdan and di-4-ANNEPS led us to develop an alkyne modified Nile Red derivative that can be conjugated to LipoTag, resulting in the red-fluorescent ratiometric membrane order probe LipoTag-NR (Fig. 4a-b).

**Figure 4:**
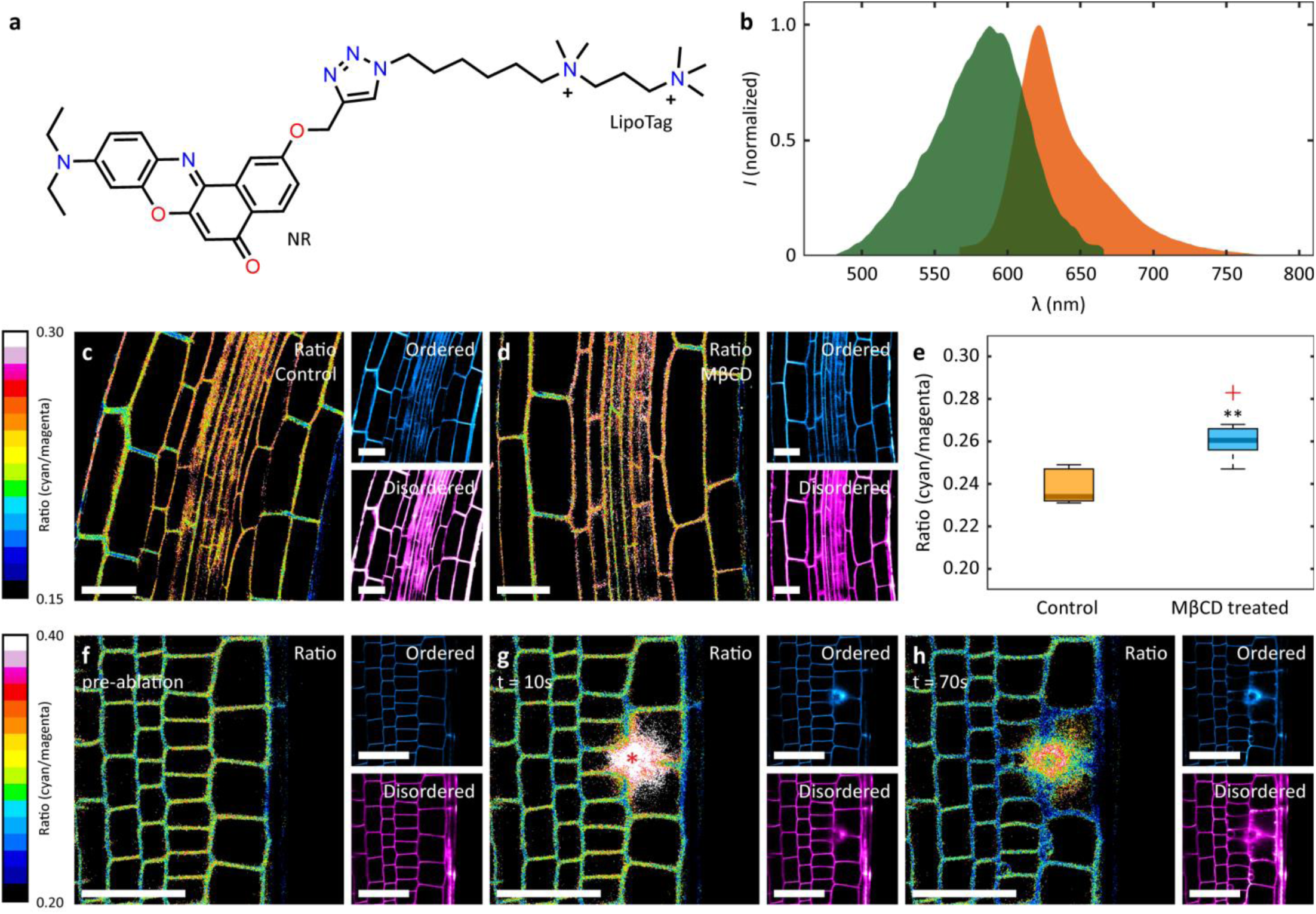
LipoTag-NR based imaging of cell membrane order. **a,** Chemical structure of LipoTag-NR membrane order probe. **b,** Normalized excitation (green) and emission (orange) spectra of LipoTag-NR. **c,d,** Ratiometric images of Arabidopsis roots treated without (**c**) and with MβCD (**d**). The color corresponds to the cyan fluorescence signal (ordered) divided by the magenta fluorescence signal (disordered). Ordered and disordered signals are shown separately in cyan and magenta, respectively. **e,** Comparison of the cyan/magenta ratio between control and MβCD treated roots. **f-h,** Timelapse of LipoTag-NR stained Arabidopsis roots before (**f**), 10 seconds (**g**) and 70 seconds (**h**) after ablation. Ablation site is indicated by a red asterisk. The color corresponds to the cyan fluorescence signal (ordered) divided by the magenta fluorescence signal (disordered). Ordered and disordered signals are shown separately in cyan and magenta, respectively. In box plots the center line is the median, box bounds represent the 25th and 75th percentiles. Whiskers span from the smallest to the largest data points not considered outliers. Red + symbols indicate outliers. **P < 0.01; two-sided Wilcoxon rank sum test. Scale bars represent 25 µm

To evaluate the responsiveness of this probe towards changes in membrane order we performed a sterol depletion experiment in which we treated roots of Arabidopsis seedlings with methyl-beta-cyclodextrin (MβCD) (Fig. 4c-e). This treatment is used in mammalian cell research to remove cholesterol, which in turn decreases membrane order [68, 70, 71]. Intriguingly, in plants, treatment with MβCD results in the opposite: LipoTag-NR reveals an increase in membrane order in the root elongation zone (Fig. 4e). While we do not have a definitive answer to this surprising result, and it does reveal the responsiveness of our probe, it is worth noting that plant membranes, unlike mammalian cells, contain 3 different sterols with antagonistic effects: campesterol, sitosterol and stigmasterol [72]. Whereas campesterol and sitosterol have a lipid ordering effect, similar to cholesterol, stigmasterol promotes membrane disorder. Plants adjust the ratio between these 3 phytosterols plants to prevent sharp cell membrane phase transitions and maintain membrane integrity over a range of temperatures and during growth and division [73, 74]. As the elongation zone undergoes continuous expansion, a high degree of membrane fluidity may be required to accommodate the rapid increase in surface area and prevent membrane rupture. This may in turn require that that more stigmasterol is present. Once removed by the MβCD this could lead to the observed increase in membrane order. Although stigmasterol levels have not been specifically determined in the elongation zone of Arabidopsis roots, it is known that inhibition of stigmasterol biosynthesis leads to reduced root growth [75].

Here, we also used laser ablation to induce damage and thereby generate a mechanical imbalance in a plant tissue; more specifically in the epidermal-cortex cell layer in Arabidopsis root tips. This is comparable to the ablation done on roots stained with LipoTag-BDP but since the readout of the LipoTag-NR probe is ratiometric, and hence does not require FLIM, we were able to track membrane deformation with a 10 second interval. Upon ablation an immediate decrease in lipid order of cells neighboring the damage site is observed (Fig. 4f-h). Interestingly, this decrease appears to be more pronounced in periclinal membranes. This indicates that the orientation of the membrane relative to the ablation site determines how strong the lipid order decreases, either because of the way the mechanical imbalance affects differently oriented membranes or because in their native configuration these membranes were already polarized. These results demonstrate how LipoTag-NR can be used to study the responses of membranes towards an altered chemical composition or changes in mechanical homeostasis.

### Imaging lipid oxidation: LipoTag-Ox

Plant membranes continuously change in composition to accommodate developmental processes or to adapt to changes in their environment [76]. Changes in lipid composition via biosynthetic pathways however are relatively slow. By contrast, chemical oxidation of unsaturated cell membrane lipids by reactive oxygen species (ROS) is a rapid mechanism by which organisms can alter membrane properties and activate membrane proteins [77]. Despite its high reactivity, plants maintain ROS levels that is essential for cell growth and division [78]. Elevated ROS levels or transient bursts of ROS are associated with cellular stress and are known to mediate stress signaling [79]. Here, we developed a ROS sensitive lipid peroxidation probe (LipoTag-Ox), a red-fluorescent probe that is based on the BDP581/591 fluorophore. This contains a large conjugated system with two double bonds being susceptible towards oxidation (Fig 5 a-b). Upon oxidation, the conjugated system shortens and the emission shifts from red to green. This allows its use in the ratiometric imaging of lipid oxidation.

**Figure 5:**
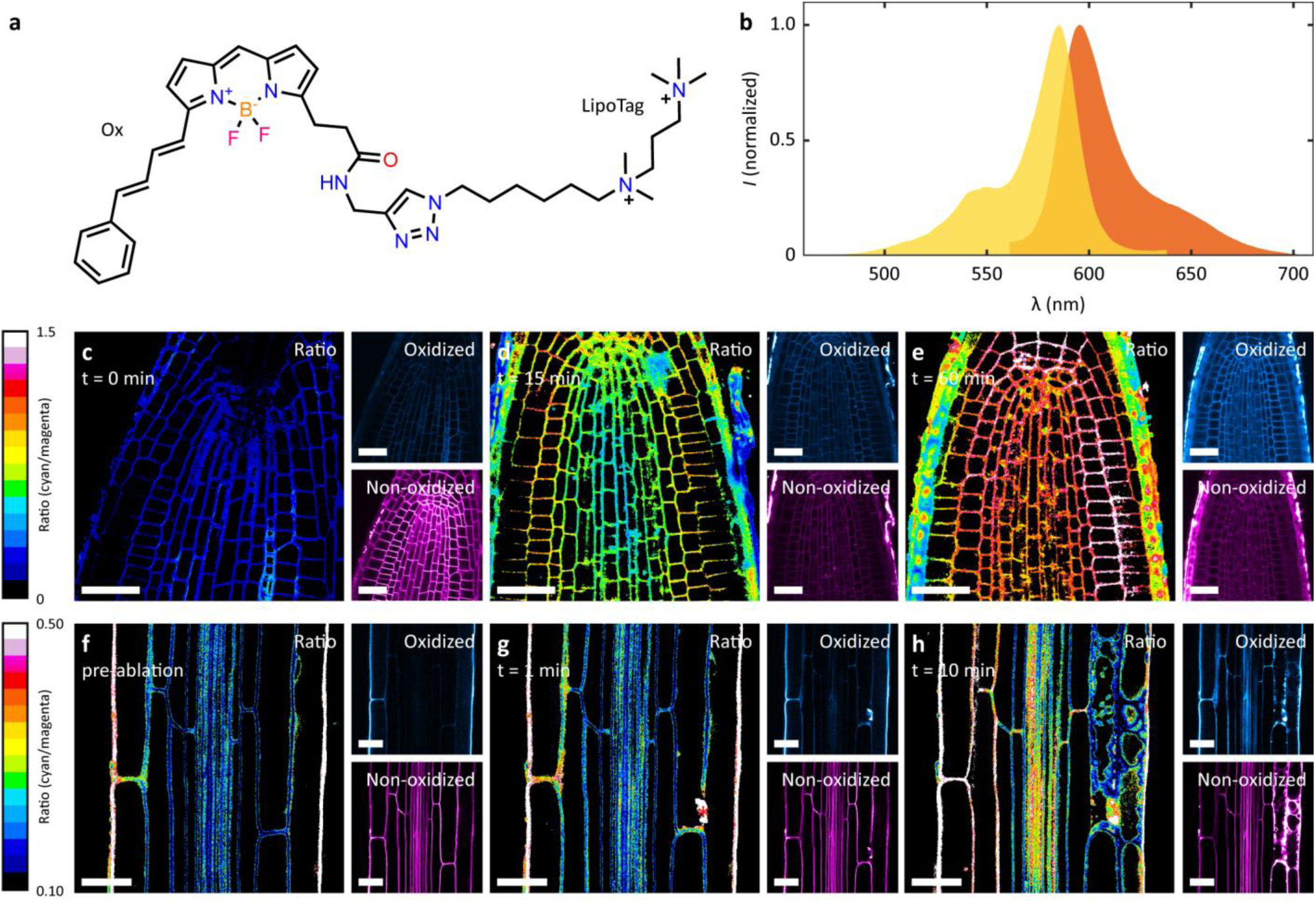
LipoTag-Ox based imaging of cell membrane oxidation. **a,** Chemical structure of LipoTag-Ox membrane oxidation probe. **b,** Normalized excitation (yellow) and emission (orange) spectra of LipoTag-Ox. **c-e,** Timelapse of LipoTag-Ox stained roots treated with exogenous ROS at t = 0 minutes (**c**), t = 15 minutes (**d**) and t = 60 min (**e**). The color corresponds to the cyan fluorescence signal (oxidized) divided by the magenta fluorescence signal (non-oxidized). Oxidized and non-oxidized signals are shown separately in cyan and magenta, respectively. **f-h,** Timelapse of LipoTag-Ox stained Arabidopsis roots before (**f**), 1 minute (**g**) and 10 minutes (**h**) after ablation. Ablation site is indicated by a red asterisk. The color corresponds to the cyan fluorescence signal (oxidized) divided by the magenta fluorescence signal (non-oxidized). Oxidized and non-oxidized signals are shown separately in cyan and magenta, respectively. Scale bars represent 25 µm

Also this probe, conjugated to the LipoTag motif, permeates and labels plant membranes well and quickly. To test the ability of this probe to detect ROS, we treated Arabidopsis roots with exogenous radical oxygen generated by a chemical, so-called Fenton, reaction between hemin and cumene peroxide [80]. This resulted in the rapid oxidation LipoTag-Ox within an hour, which we tracked over time (Fig. 5c-e). Mock treatment did not show significant oxidation, confirming that normal root metabolism or solvent induced stress are not responsible for the probe’s oxidation Fig. S19).

Measuring membrane oxidation caused by endogenously produced ROS is a more relevant situation with useful applications in plant research. To test this, we performed laser ablations in the epidermal-cortex layer to elicit a damage-associated ROS burst [81]. As these responses are reported to be tissue-specific, we performed these measurements in the differentiation zone of Arabidopsis roots, which is reported to be sensitive towards ablations [82]. Following ablation, we observed immediate oxidation of membrane material near the ablation site. More notably, the ablation appears to induce a systemic response since oxidation of cell membranes on the other side of the vasculature tissue is also observed within 10 minutes (Fig. 5f-h). This oxidation does not appear to occur homogeneously but rather starts at the epidermis-cortex layer and moves inward. The mechanism of ROS generation and membrane oxidation cannot be deduced using this probe but previous research suggests the involvement of Ca^2+^ waves that trigger the activation of NADPH oxidases [83].

## Discussion

In this paper we demonstrated LipoTag as a minimal and modular chemical motif for live and functional imaging of plant cell membranes. LipoTag can transform hydrophobic fluorophores into water-soluble probes that localize in membranes of walled organisms. We noted that the kinetics of probe internalization by endocytosis are sensitive to the physicochemical properties of the fluorophore, specifically, the use of lipophilic cations, such as Cy5 and oxazine, appears to accelerate the rate of internalization compared to neutral fluorophores. Compared to the current standard in plant membrane imaging, our LipoTag toolbox offers the advantage of spectral flexibility, rapid tissue permeability and not having to rely on potentially toxic permeabilization agents. LipoTag probes work in a range of tested plant species and can also stain cell membranes in walled organisms beyond the plant kingdom, which was previously challenging. Moreover, it allowed us to construct a set of probes for functional imaging of cell membrane properties that were demonstrated to allow spatial mapping of membrane adaptations without the need for invasive techniques or genetic engineering of the organism.

### Limitations of LipoTag

Although the LipoTag platform provides spectral flexibility, we noted limitations in what fluorophores can be used as its cargo, in particular related to their net charge. For now, generating LipoTag probes was successfully achieved for BODIPY dyes and Nile Red derivatives; and we foresee it could also work for other uncharged aromatic systems such as coumarins, pyrenes and perylenes. However, probes from the cyanine and oxazine family were found not to be suitable as their additional charge gave rise to rapid internalization by endocytosis and subsequent localization in endomembranes. The required incubation time for good staining seems to depend on the relative orientation of the fluorophore and LipoTag moiety, where a perpendicular orientation (LipoTag-Green) results in significantly slower movement through membranes. Also, the cuticle can form a barrier layer to prolong the required staining time, as found here for the specific cases of ferns and liverworts, whose cuticles resulted in minimal staining times of 2h for a good signal. Also, the medium can be disruptive as we found that brown algae in sea water show limited staining, which we attribute to the effects of the high ionic strength on the physical stability of the probes.

Care should also be exercised in the interpretation of the functional imaging results. The FLIM-based LipoTag-BDP for cell membrane density was designed to be responsive to its local environment. Besides being rigidochromic, these probes are typically solvatochromic as well which means they are sensitive towards their chemical environment as well. We observed that the fluorescence lifetime of LipoTag-rotor in the outer membrane of root epidermal cells is consistently below the lowest lifetime for the probe in calibration media. This indicates probe quenching, likely caused by aromatic components of the cuticle surrounding the epidermis which complicates the study of the outer epidermal membranes with FLIM. Finally we note that, while the functional probes presented in this work can in principle be calibrated *in vivo*, e.g. in reconstituted membranes such as GUVs [35], we ask for caution in transferring such calibrations to *in vivo* measurements [34]. Translating calibration data from simple *in vitro* experiments to the reality of *in vivo* membranes is dangerous as the physical and chemical microenvironment in plants is too complex to be captured in minimalistic model systems. As such, we recommend that our functional probes are used for the spatio-temporal mapping of relative changes in plant membranes, e.g. as the result of exogeneous signals or by comparing wild-type plants to specific mutants, but not to attempt to quantify these changes in the context of absolute physico-chemical metrics.

### Outlook

Based on our experiences in developing these tools, we foresee that LipoTag can be extended to beyond what we report here. The modularity of LipoTag allows for simple expansion of the toolbox of functional probes with additional functionalities such as membrane potential and composition, if the design constraints explained above are kept in mind. Moreover, we anticipate that LipoTag can be used to create active functionality, for example by attaching photoactivatable fluorophores or motors to enable single-molecule based super resolution imaging of plant membranes or induce membrane rupture upon UV-stimulation, respectively [84, 85]. LipoTag, through its modularity and easy-of-use, opens the way for imaging membranes of walled organisms with unprecedented spectral flexibility without the need for sophisticated chemical synthesis.

## Materials and Methods

A detailed overview of synthetic procedures, NMR characterization of synthesized compounds and statistical details for all experiments can be found in the supporting information file.

### LipoTag fluorophore general conjugation protocol

10 mg of alkyne modified fluorophore was dissolved in 4.5 ml DMF. CuSO4 and THPTA were added from a 10mM and 5 mM stock solution in water so their final concentration is 200 and 500 uM respectively. 2 equivalents of LipoTag were added (compared to fluorophore) from a 100 mM stock solution in water. The resulting solution was stirred and bubbled with N2 before the addition of sodium ascorbate from a 10 mM stock solution in water so its final concentration is 400 uM. The resulting mixture is left stirring overnight before being concentrated under air flow. The resulting product was run over a basic alumina plug, first washing with 10:90 MeOH/DCM followed by elution with 100% MeOH.

### Imaging

Imaging experiments were performed on a Leica TCS SP8 inverted confocal microscope coupled to a pulsed white light laser or a Leica Stellaris STED microscope coupled to a pulsed white light laser and a 405 nm diode laser. Experiments on brown algae that were performed on a Zeiss LSM880 confocal microscope. Functional imaging with LipoTag-NR and LipoTag-Ox, FRAP and ablations were performed on a Leica SP8 multiphoton system with a pulsed Coherent Chameleon Ti:sapphire laser. Fluorescence was captured through a ×20 ((NA) 0.75) dry objective, a 40x (NA 1.1) water objective, a ×63 (NA 1.2) water objective, a ×86 (NA 1.2) water objective or a ×100 (NA 1.4) oil objective depending on the sample type.

LipoTag-Green, LipoTag-BDP (membrane tension probe) and FM1-43 were excited with a 488 nm laser line, fluorescence was captured between 500 and 600 nm using HyD detectors. FM4-64 was excited with a 488 nm laser line and its fluorescence captured between 650 and 750 nm. LipoTag-Orange was excited with a 594 nm laser line, fluorescence was captured between 605 and 655 nm using a HyD detector. LipoTag-Red was excited with a 625 nm laser line, fluorescence was captured between 635 and 700 nm using a HyD detector. LipoTag-Oxa was excited with a 666 nm laser line, fluorescence was captured between 680 and 780 nm using a HyD detector. LipoTag-Cy5 was excited with a 646 nm laser line, fluorescence was captured between 656 and 689 nm using a HyD detector. LipoTag-NR was excited with a 830 nm laser line on a Leica SP8 multiphoton system, fluorescence was captured between 500 and 585 nm for the ‘ordered’ signal and between 585 and 700 nm for the ‘disordered’ signal using HyD detectors. LipoTag-Ox was excited with a 980 nm laser line on a Leica SP8 multiphoton system, fluorescence was captured between 500 and 560 nm for the ‘oxidized’ signal and between 590 and 620 nm for the ‘non-oxidized signal using HyD detectors.

LifeAct-GFP, was excited with a 488 nm laser line, fluorescence was captured between 500 and 600 nm using a HyD detector. DAPI was excited with a 405 nm laser line, fluorescence was captured between 420 and 553 nm using a HyD detector. CellMask Orange was excited with a 561 nm laser line, fluorescence was captured between 570 and 620 nm using a HyD detector. Aniline Blue was excited with a 405 nm laser line, fluorescence was captured between 420 and 540 nm using a HyD detector. Chlorophyl autofluorescence wat captured with varying excitation wavelengths depending on the experiment, fluorescence was captured in experiment depending windows between 700 and 770 nm.

For multicolor experiments excitation and emission wavelengths of LipoTag dyes were sometimes changed to be spectrally compatible with other fluorophores.

### STED

Super-resolution STED imaging was performed on a Leica Stellaris STED microscope coupled to a pulsed white light laser and pulsed 775 nm depletion laser. LipoTag-Red was excited and depleted with a 638 nm laser line and 775 depletion laser at 20% respectively. Fluorescence was collected between 640 and 670 nm through a ×100 (NA 1.4) oil objective. The TauGate function was applied of 1-11 ns to limit autofluorescence. Line plots of plasmodesmata were made in Fiji, exported to and normalized in excel before being plotted in MATLAB2023b.

### FRAP

FRAP was performed on six day old Arabidopsis seedlings using a Leica SP8 multiphoton system equipped with diode lasers and a PMT detector. A ROI of constant size was drawn on the epidermis-cortex barrier and ablated using a 488 nm diode laser (LipoTag-Green), a 552 nm diode laser (LipoTag-Orange and FM4-64) or a 638 nm diode laser (LipoTag-Red). After bleaching images were taken with 1.3 s interval, before bleaching 3 reference images were taken. Images were first analyzed in Fiji to obtain intensity values for the bleached ROI, a reference ROI and a background ROI for each frame. These values were loaded into excel, the background value subtracted from the bleach ROI and reference ROI per frame. The bleach ROI was normalized to 0, then normalized to 1 by using the average intensity of the first 3 reference images before being divided by the normalized reference ROI, yielding normalized, background and photobleaching corrected intensities. After combining the normalized values a standard deviation was determined and data plotted in MATLAB2023b.

### Ablation

Ablation was performed on a Leica SP8 multiphoton system with a pulsed Coherent Chameleon Ti:sapphire infrared laser. Ablation was performed by selecting 3-4 point for ablating with 100-200 ms exposure times at approximately 50% laser power, 980 nm for experiments with LipoTag-Ox and 830 nm for experiments with LipoTag-NR and LipoTag-BDP. Ablated samples with LipoTag-BDP were quickly transferred to a microscope with FLIM capabilities. For experiments with LipoTag-NR and LipoTag-Ox acquisition was starting immediately after ablating with a 10 second and 60 second interval respectively.

### FLIM

FLIM experiments using LipoTag-BDP were performed on a Leica TCS SP8 inverted confocal microscope coupled to a Becker-Hickl SPC830 time-correlated single photon counting module. An excitation line of 488 nm with repetition rate of 20 MHz was used and fluorescence was captured between 500 and 600 nm using a hybrid detector. Acquisition times were between 60 and 120 s were used depending on signal strength. Resulting FLIM 256×256 images were processed in SPCImage v.8.5 software to obtain two-component exponential decay curves for every pixel. . FLIM images are presented in a false-color scale that represents the mean fluorescence lifetime per pixel in nanoseconds.

### Ratiometric analysis

Ratiometric analysis for LipoTag-NR and LipoTag-Ox was performed the same way. 2 channel images and stacks are first converted to a 32 bit format, the pixel values multiplied by 5 and a Gaussian blur with a sigma of 0.5 applied. A threshold was set manually for the first channel and applied to both channels. The pixel values of the first channel where then divided by the pixel values of the second channel, yielding a ratiometric image in which the values are represented by a 16-color LUT.

### Toxicity assay

Seeds (a volume of about 800 µl) were sterilized in 10 mL 70% ethanol for 20 min, followed by 10 mL 0.8% bleach for 40 min. Seeds were washed three times with MQ and plated in three dense rows on a 1/2X MS-agar plate containing ampicillin. Seeds were stratified for 2 nights at 4 °C and then transferred to a growth cabinet. After 10 days, the green parts are harvested by cutting them using a scalpel. Protoplasts were isolated using a standard protocol. 250 µL of cell suspension was added to each well of a 24-well plate. LipoTag dyes, FM4-64 and non-modified LipoTag were added at various final concentrations (0.1 µM, 0.5 µM 1 µM and 10 µM for LipoTag-Orange and LipoTag-Red; 2 µM, 10 µM and 20 µM for FM4-64; 1 µM, 2 µM and 10 µM for non-modified LipoTag) and incubated for 1h.To each well, 1 µl FDA (5 mg/mL in acetone) was added. Wells were incubated for a minimum of 10 min before 5 images per well were taken on an epifluorescence microscope to detect both brightfield and FDA signal. Viability was calculated by calculating the FDA cell count as a fraction of the total cell count, expressed as a percentage.

### Organism growth and staining conditions

*Arabidopsis thaliana* Col-0 wild-type and marker line seeds were surface sterilized by washing them in a 50% ethanol solution, followed by a wash in 70% ethanol and a 96% ethanol solution (5 min per solution). The ethanol solution was replaced by MQ water and seeds were left to swell for 5 minutes. Sterilized seeds were put on half strength (0.5×) MS plates and placed at 4 °C overnight before placing them vertically under long-day growth conditions (16 h light, 8 h dark) for 6-7 days. Seedlings were transferred to 0.5x MS supplemented with 0.5 µM LipoTag dye and incubated for at least 30 minutes unless stated otherwise. Seedlings were transferred to clean 0.5x MS for 1 minute and subsequently imaged.

*M. polymorpha* was grown on B5 medium under continuous light (40 μmol m−2 s−1) at 25 °C. Gemma were isolated using a 200-μl pipet tip before being transferred to 0.5xMS supplemented with 0.5 µM LipoTag dye. Staining with Aniline Blue was performed by directly placing gemma in a 0.1% Aniline Blue solution on a microscopy slide, covering it with a cover slip and immediately image.

*C. richardii* spores (Hn-n strain) were sterilized and grown as previously describe in a Hettich MPC600 plant growth incubator set at 28 °C, with 16 h of 100 μmol m−2 s−1. Plants were grown on 0.5× MS medium supplemented with 1% sucrose. Gametophytes were grown from spores and synchronized by imbibing the spores in the dark in water for >4 days. Gametophytes were transferred to 0.5x MS supplemented with 0.5 µM LipoTag dye and incubated for 2 hours under gentle shaking before being transferred to clean 0.5x MS and subsequently imaged.

All brown algae were grown in Provasoli enriched sea water (PES) under low white light conditions with a 12 h/12 h light/dark cycle (Le Bail and Charrier, 2013). Temperatures and light intensity varied depending on the algal species. Male gametophytes of *Sphacelaria rigidula* were cultivated at 16 °C under 30 μmol m^−2^ s^−1^ of light intensity. Filaments of *Ectocarpus sp*. sporophytes were grown at 13°C under 16 μmol m^−2^ s^−1^ (Le Bail and Charrier, 2013). Fragments of female and male *Saccharina latissima* gametophytes were mixed and inoculated at 13°C in 16 μmol m^−2^ s^−1^ (Theodorou et al., 2021). Once zygotes of *S. latissimi* were formed, the material was transferred to 50 μmol m^−2^ s^−1^ of light intensity. Embryos of *Fucus serratus* were produced from fertile female and male thalli collected in Perharidy, Roscoff, France (GPS coordinates Latitude : 48.726288 | Longitude : −3.99972) and cultivated at 16 μmol m^−2^ s^−1^ as described in Siméon and Hervé (2017). Brown algae were stained by adding 0.5 µM LipoTag dye to the medium and incubating for a minimum of 30 minutes before washing with fresh PES (3x) and imaging. Verticillium dahliae wild-type strain JR2 was cultured on potato dextrose agar (PDA) plates at room temperature for 5 days. Conidiospores were harvested and suspended in sterile distilled water to a final concentration of 5 × 10⁶ conidia mL⁻¹. The conidial suspension was treated with LipoTag dyes at final concentrations of 0.5 µM. Subsequently, 20 µL of each treated suspension was placed onto a microscope slide and covered with a 24*24 mm coverslip.

*Verticillium dahliae* wild-type strain JR2 was cultured on potato dextrose agar (PDA) plates at room temperature for 5 days. Conidiospores were harvested and suspended in sterile distilled water to a final concentration of 5 × 10⁶ conidia mL⁻¹. The conidial suspension was treated with LipoTag dyes at final concentrations of 0.5 µM. Subsequently, 20 µL of each treated suspension was placed onto a microscope slide and covered with a 24*24 mm coverslip.

*S. pombe* (DB146) pre-culture was inoculated by taking a small sample from plate to 5mL YE5S medium in 50mL falcon tubes using a loop and cultured over night at 28°C,160 rpm. In an 1.5 mL Eppendorf, LipoTag probe was added with final concentration of 1μM. 100μL of stained cells were directly transferred to a chambered microscopy slide and imaged as fast as possible.

*E. coli* (DH5α) was grown in 5 ml LB at 37 °C overnight under shaking (150 rpm). The resulting was then inoculated the next morning in fresh LB (37 °C, 150 rpm) until it reached OD600 ∼0.5-0.6. 1 ml of the culture was transferred to a 1.5 ml Eppendorf and centrifuged for 6 minutes at 6000 rcf. The cells were resuspended in PBS supplemented with 5 µg/ml DAPI (in the case of LipoTag-Orange and LipoTag-Red) and 0.5 µM LipoTag dye. 1 µl of stained suspension was transferred to an agar pad and imaged.

Murine macrophage-like J774A.1 cells (ATCC, TIB-67) were maintained in complete DMEM (DMEM (Gibco, 32430-100) supplemented with 10% FBS and 1% Antibiotic-Antimycotic (Gibco, 15240–062)). Cells were maintained at 37 °C in a humidified atmosphere with 5% CO₂ and subcultured by scraping. For macrophage membrane labeling with LipoTag dyes, cells were seeded in µ-Slide 18 Well chambers (Ibidi, 81817) at a density of 3 * 10^4^ cells per well in 100 µl complete DMEM and allowed to adhere overnight. The following day, the media was removed and replaced with 100 µl complete DMEM + CellMask Orange Plasma Membrane Stain (Invitrogen, C10045, 1:1000) to obtain a plasma membrane reference stain. Cells were then incubated for 10 minutes at 37 °C followed by 3 washes with complete DMEM. Medium was then replaced by 100 µl L-15 medium (Gibco, 21083027) + 10% FBS and cells were transferred to the microscope. The Lipotag probe was diluted in L-15 medium and added to the cells at a final concentration of 0.5 µM and immediately imaged.

### Chemical treatments

**ABA.** Arabidopsis seedlings were stained with 0.5 uM LipoTag-rotor and simultaneously treated with 20 μM ABA from a 10 mM stock for 1 h to induced stomata closure.

**MβCD.** Seedlings were incubated in 0.5xMS supplemented with 5 mM MβCD (from 5 mM stock in water) for 15 minutes, transfered to clean 0.5x MS supplemented with 0.5 µM LipoTag-NR for 15 minutes before being washed and imaged.

**Hemin cumene peroxide.** Arabidopsis seedlings were stained for 30 minutes in 0.5x MS supplemented with 0.5 µM LipoTag-Ox. Seedlings were transferred to fresh 0.5xMS supplemented with 2 mM cumene peroxide (from 500mM stock in EtOH) an 5 µM hemin (from 1 mM stock in DMS). They were incubated for a specific time, transferred to clean 0.5xMS and imaged.

## Acknowledgements

This work and J.S., M.B. and J.L. are funded by the European Research Council project Catch, project number 101000981. The work of B.C. and M.Z. are funded by the European Research Council project ALTER e-GROW, project number 101055148. Views and opinions expresses are, however, those of the author(s) only and do not necessarily reflect those of the European Union or the European Research Council Executive Agency. The work of T.H. is supported by the Dutch Research Council (NWO) through the gravitation program GreenTE with project number 024.006.001. The work of R.F. is financially supported by the TopSector TKI Horticulture and Starting Materials, the Netherlands, through the project TU202312. The work of R.R and H.T are funded by the Dutch Research Council (NWO) project Battle at the Wall with file number VI.C.242.004 and grant ID https://doi.org/10.61686/ZGIFD55241. This publication is part of the project NL-BioImaging-AM (with project number 184.036.012) of the National Roadmap research program, which is (partly) financed by the Dutch Research Council (NWO). We gratefully acknowledge invaluable discussions, suggestions, and the supply of plant and bacterial material by A. Daamen, M. de Roij, A Bellandi, Y. Hata,A. Sanz Andres, C. Borassi, D. Weijers, CH Ang and S Koh Wee Han

## Author contributions

M.B. and J.S. established the LipoTag concept. M.B., J.S. and J.W.B. designed the experiments and contributed to data interpretation. M.B. designed and performed the synthesis of the LipoTag motif. M.B. synthesized all LipoTag-based probes and performed all plant imaging experiments. J.L. determined the evolutionary relationships and common ancestry of organisms used in this research and developed a phylogenetic tree. Imaging experiments on brown algae were performed by B.C. and M.Z. Imaging experiments on fungi were performed by H.T. Imaging experiments on yeast were performed by R.R. Imaging experiments on mammalian macrophages were performed by Y.P. and D.V. Toxicity assays were performed and analyzed by R.F. Initial tests on staining kinetics and FRAP of LipoTag-Orange and FM4-64 were performed by T.H. J.S. and J.W.B. supervised the project. M.B., J.S. and J.W.B. wrote the manuscript. All authors reviewed the paper.

