## Supplementary information for "LipoTag: A minimal motif for live and functional imaging of plant cell membranes"

**Table of contents**


| | |
|---|---|
| **Synthetic procedures LipoTag precursors, fluorophore precursors and LipoTag modified fluorophores** | **2-8** |
| **Figure S1, staining of Arabidopsis with commonly used mammalian membrane probes and LipoTag dye precursors** | **9** |
| **Figure S2, concentration range FM4-64 in different Arabidopsis root zones** | **10** |
| **Figure S3, penetration depth of FM4-64 in different tissues over time** | **11** |
| **Figure S4, effect of staining concentration of non-functional LipoTag probes on tissue penetration after 30 minutes** | **12** |
| **Figure S5, staining kinetics of LipoTag-Green in the root tip and elongation zone** | **13** |
| **Figure S6, staining kinetics of LipoTag-Orange in the root tip and elongation zone** | **14** |
| **Figure S7, staining kinetics of LipoTag-Red in the root tip and elongation zone** | **15** |
| **Figure S8, chemical structures of non-functional LipoTag probes** | **16** |
| **Figure S9, plasmolysis of LipoTag stained Arabidopsis roots** | **17** |
| **Figure S10, staining of Arabidopsis roots with LipoTag-Cy5 and LipoTag-Oxa.** | **18** |
| **Figure S11, staining of F. serratus with LipoTag probes** | **19** |
| **Figure S12, staining of Ectocarpus with LipoTag probes** | **20** |
| **Figure S13, staining of *S. rigidula* with LipoTag probes** | **21** |
| **Figure S14, staining of *S. latissima* with LipoTag probes** | **22** |
| **Figure S15, staining of Murine macrophage cells with LipoTag dyes** | **23** |
| **Figure S16, staining of plasmodesmata in Marchantia gemmae** | **24** |
| **Figure S17, calibration of LipoTag-BDP** | **25** |
| **Figure S18, staining of Arabidopsis root with NR12S** | **26** |
| **Figure S19, mock hemin treatment with LipoTag-Ox** | **27** |
| **Figure S20, chemical structures of functional LipoTag probes** | **28** |
| **Figure S21, normalized fluorescence spectra of all LipoTag probes** | **29** |
| **Table S1, number of samples** | **30-31** |
| **Figure S22-45, NMR-spectra** | **32-55** |

**LipoTag fluorophore conjugation, LipoTag and LipoTag precursors synthesis**

**LipoTag fluorophore general conjugation protocol.**

10 mg of alkyne modified fluorophore was dissolved in 4.5 ml DMF. $CuSO_4$ and THPTA were added from a 10mM and 5 mM stock solution in water so their final concentration is 200 and 500 µM respectively. 2 equivalents of LipoTag were added (compared to fluorophore) from a 100 mM stock solution in water. The resulting solution was stirred and bubbled with N2 before the addition of sodium ascorbate from a 10 mM stock solution in water so its final concentration is 400 µM. The resulting mixture is left stirring overnight before being concentrated under air flow. The resulting product was run over a basic alumina plug, first washing with 10:90 MeOH/DCM followed by elution with 100% MeOH.

**1-azido-6-chlorohexane (1)**

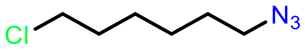


1-bromo-6-chlorohexane (6.9 ml, 46.2 mmol) was dissolved in 50 ml DMF. $NaN_3$ (3g, 46.2 mmol) was added and the mixture left to stir overnight at room temperature. The mixture was diluted with 60 ml of $H_2O$ and the product extracted with 3x 75 $Et_2O$. The combined organic layers were washed with 3x100 ml $H_2O$. The organic layer was dried with $MgSO_4$ and concentrated under reduced pressure, yielding 6.99 g of **1** as an oil. Yield: 94%.$^{1}$**H NMR** (400 MHz, $CDCl_3$) δ 3.53 (t, *J* = 6.6 Hz, 2H), 3.27 (t, *J* = 6.9 Hz, 2H), 1.83 – 1.73 (m, 2H), 1.65 – 1.56 (m, 2H), 1.51 – 1.35 (m, 4H).$^{13}$**C NMR** (101 MHz, CDCl3) δ 51.33, 44.90, 32.41, 28.73, 26.43, 26.04.

**1-azido-6-iodohexane (2)**

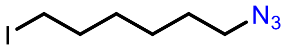


**1** (6.9 g, 42.7 mmol) and NaI (12.77 g, 85.2 mmol) were added to 200 ml acetone. The mixture was bubbled with $N_2$ for 10 minutes and then left stirring for 24h under reflux. After cooling the mixture was diluted with 100 ml $H_2O$ and extracted wit 3x 100 ml EtOAc. The combined organic layers were dried with $MgSO_4$ and concentrated under vacuum, yielding 5.88g of **2**. Yield: 54%. $^{1}$**H NMR** (400 MHz, $CDCl_3$) δ 3.26 (t, *J* = 6.9 Hz, 2H), 3.18 (t, *J* = 6.9 Hz, 2H), 1.88 – 1.72 (m, 2H), 1.66 – 1.53 (m, 2H), 1.51 – 1.32 (m, 4H). $^{13}$**C NMR** (101 MHz, $CDCl_3$) δ 51.31, 33.24, 30.00, 28.67, 25.67, 6.77.

**6-azido-N-(3-(dimethylamino)propyl)-N,N-dimethylhexan-1-aminium (3)**

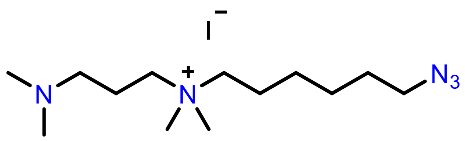

**2** (800 mg, 3.2 mmol) and N,N,N′,N′-tetramethyl-1,3-propanediamine (15 ml, 90 mmol) were dissolved in 40 ml of dry THF and left stirring overnight at room temperature. The mixture was concentrated under vacuum at 50 °C and washed three times with $Et_2O$ and dried under vacuum yielding 1.12 g of **4**. Yield: 91%. **$^1$H NMR** (400 MHz, MeOD) δ 3.43 – 3.32 (m, 6H), 3.11 (s, 6H), 2.41 (t, J = 7.1 Hz, 2H), 2.28 (s, 6H), 1.99 – 1.88 (m, 2H), 1.85 – 1.74 (m, 2H), 1.69 – 1.58 (m, 2H), 1.57 – 1.38 (m, 4H). **$^{13}$C NMR** (101 MHz, D2O) δ 64.70, 64.14, 62.09, 54.76, 50.99, 50.81, 43.72, 27.74, 27.70, 25.50, 25.45, 25.03, 21.91, 21.80, 19.77.

**LipoTag (4)**

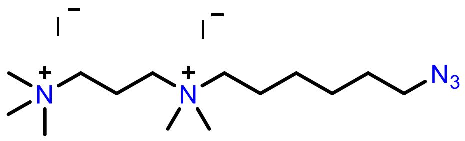

**3** (400 mg, 1.04 mmol) was added to 5 ml of DMF followed by 2 ml of MeI (31 mmol). The mixture was left stirring overnight at room temperature and then concentrated under vacuum at 50 °C and washed with Et2O yielding 508 mg of **4** as a viscous, brown oil. Yield: 93%. **$^1$H NMR** (400 MHz, $D_2O$) δ 3.52 – 3.34 (m, 9H), 3.24 (s, 9H), 3.18 (s, 6H), 2.37 (s, 2H), 1.84 (s, 2H), 1.66 (s, 2H), 1.47 (s, 4H). **$^{13}$C NMR** (101 MHz, $D_2O$) δ 64.89, 62.48, 60.06, 53.36, 51.00, 34.57, 27.74, 25.49, 25.02, 21.90, 17.19.

**4-(prop-2-yn-1-yloxy)benzaldehyde (5)**

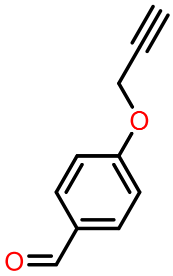


4-hydroxy-benzaldehyde (3g, 24.6 mmol) was dissolved in 250 ml of acetone. $K_2CO_3$ (13.58g, 98.3 mmol) and propargyl bromide (10.5 ml from a 80% solution with toluene, 98.3 mmol) were added and the mixture was bubbled with $N_2$ for 5 minutes. The mixture was heated to reflux for 4 hours, cooled to room temperature and diluted with 250 ml $H_2O$ (all residual $K_2SO_3$ needs to be dissolved). The aqueous solution was extracted with 3x 200 ml DCM, the combined organic layers were dried with MgSO4 and concentrated under vacuum, yielding 3.6g of **5** as a brown solid. Yield: 91%.$^{1}$**H NMR** (400 MHz, CDCl3) δ 9.91 (s, 1H), 7.86 (d, J = 8.8 Hz, 2H), 7.09 (d, J = 8.8 Hz, 2H), 4.78 (d, J = 2.4 Hz, 2H), 2.57 (t, J = 2.4 Hz, 1H). $^{13}$**C NMR** (101 MHz, CDCl3) δ 190.77, 162.38, 131.90, 130.62, 115.19, 77.55, 76.37, 55.96.

**BDP-green alkyne (6)**

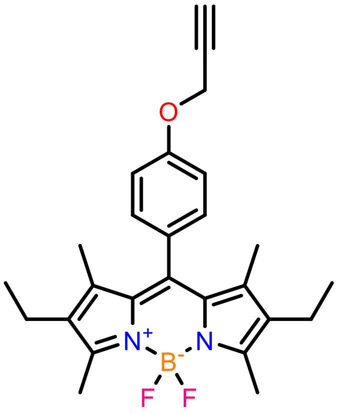


**5** (500 mg, 3.12 mmol) and 2,4-dimethyl-3-ethyl-pyrrole (769 mg, 6.24 mmol) dissolved in 125 ml of dry DCM. The mixture was bubbled with $N_2$ for 30 minutes after which TFA (125 µl, 1.5 mmol) was added and the mixture was left to stir for 2 hours. 2,3-Dichloro-5,6-dicyano-1,4-benzoquinone (DDQ, 708 mg, 3.12 mmol) was added an the mixture left to stir for 30 minutes. DIPEA (3.75 ml, excess) and $BF_3$-$OEt_2$ (3.8 ml, excess) were added subsequently and the mixture was left stirring overnight at room temperature. After purification on silica (2:3 EtOAc:PE) the 437 mg of **6** was isolated as a red, crystalline solid. Yield: 32%. $^{1}$**H NMR** (400 MHz, CDCl3) δ 7.19 (d, J = 8.6 Hz, 2H), 7.08 (d, J = 8.7 Hz, 2H), 4.77 (d, J = 2.4 Hz, 2H), 2.56 (t, J = 2.4 Hz, 1H), 2.53 (d, J = 1.3 Hz, 6H), 2.30 (q, J = 7.5 Hz, 4H), 1.33 (s, 6H), 0.98 (t, J = 7.5 Hz, 6H). $^{13}$**C NMR** (101 MHz, CDCl3) δ 158.11, 153.76, 140.07, 138.54, 132.85, 131.25, 129.65, 129.01, 115.69, 78.26, 75.98, 56.19, 17.21, 14.76, 12.63, 11.96.

**BDP-rotor alkyne (7)**

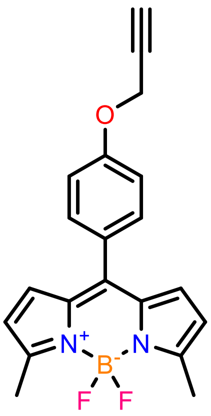


**5** (500 mg, 3.12 mmol) and 2-methyl-1H-pyrrole (506 mg, 6.24 mmol) dissolved in 125 ml of dry DCM. The mixture was bubbled with $N_2$ for 30 minutes after which TFA (125 µl, 1.5 mmol) was added and the mixture was left to stir for 2 hours. 2,3-Dichloro-5,6-dicyano-1,4-benzoquinone (DDQ, 708 mg, 3.12 mmol) was added and the mixture left to stir for 30 minutes. DIPEA (3.75 ml, excess) and $BF_3$-$OEt_2$ (3.8 ml, excess) were added subsequently and the mixture was left stirring overnight at room temperature. After purification on silica (2:3 EtOAc:PE) the 86 mg of **7** was isolated as a red, crystalline solid. Yield: 8%. **$^1$H NMR** (400 MHz, CDCl3) δ 7.47 (d, J = 8.7 Hz, 2H), 7.08 (d, J = 8.7 Hz, 2H), 6.74 (d, J = 4.1 Hz, 2H), 6.27 (d, J = 4.2 Hz, 2H), 4.78 (d, J = 2.4 Hz, 2H), 2.65 (s, 6H), 2.58 (t, J = 2.4 Hz, 1H). **$^{13}$C NMR** (101 MHz, CDCl3) δ 159.14, 157.20, 142.30, 134.51, 131.96, 130.27, 127.43, 119.26, 114.62, 78.06, 76.08, 55.92, 14.88.

**5-(diethylamino)-2-nitrosophenol (8)**

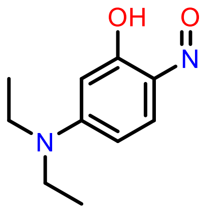


3-diethylaminophenol (8.7 g, 52.7 mmol) was dissolved in 30 ml concentrated HCl + 10 ml $H_2O$. The solution was cooled to 0 °C after which $NaNO_2$ (4.36 g, 63 mmol) dissolved in 30 ml $H_2O$ was added dropwise. The mixture was left stirring for 1h, filtered and the residue washed with a saturated sodium acetate solution. The residue was recrystallized from acetone, yielding 1.07g of **8**. Yield: 16%. **$^1$H NMR** (400 MHz, DMSO) δ 7.31 (d, J = 9.9 Hz, 1H), 6.88 (dd, J = 10.0, 2.6 Hz, 1H), 5.74 (d, J = 2.6 Hz, 1H), 3.59 (q, J = 7.1 Hz, 4H), 1.18 (t, J = 7.1 Hz, 6H). **$^{13}$C NMR** (101 MHz, DMSO) δ 169.20, 157.56, 149.71, 134.96, 115.72, 95.60, 46.07, 13.64

**NR-alkyne (9)**

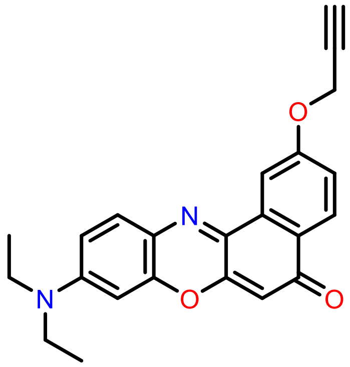

**8** (292 mg, 1.27 mmol) and 1,6-dihydroxynaphthalene (204 mg, 1.27 mmol) were dissolved in 10 ml DMF and refluxed at 155 °C for 4 hour. The mixture was cooled to 110 °C after which $K_2CO_3$ (675 mg, 5.08 mmol) was added followed by propargyl bromide (0.55 ml from a 80% solution in toluene, 5.08 mmol) and left stirring for 1 hour at 110 °C. The mixture was cooled to room temperature and diluted with $Et_2O$ (25 ml) and brine (25 ml). The aqueous layer was washed with 3x25 ml $Et_2O$ and the organic layers were combined, concentrated and purified on silica (2:1 hexane:EtOAc, 5% TEA) yielding 60 mg of **9**. Yield: 13%. **$^1$H NMR** (400 MHz, CDCl3) δ 8.25 (d, J = 8.8 Hz, 1H), 8.14 (d, J = 2.6 Hz, 1H), 7.56 (d, J = 9.1 Hz, 1H), 7.23 (d, J = 2.6 Hz, 1H), 6.66 (dd, J = 9.0, 2.7 Hz, 1H), 6.47 (d, J = 2.7 Hz, 1H), 6.33 (s, 1H), 4.90 (d, J = 2.4 Hz, 2H), 3.47 (qd, J = 7.1, 4.3 Hz, 10H), 2.58 (t, J = 2.4 Hz, 1H), 1.26 (td, J = 7.1, 5.7 Hz, 18H).

**1-(prop-2-yn-1-yl)-1,2,3,4-tetrahydroquinolin-7-ol (10)**

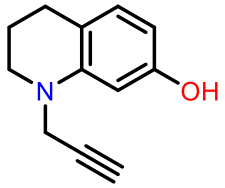

7-hydroxy-1,2,3,4-tetrahydroquinoline (1g, 6.67 mmol) and $K_2CO_3$ (1.1 g, 8 mmol) were dissolved in 8.5 ml of dry DMF. Propargyl bromide (1.5 ml from a 80% solution in toluene, 10 mmol) was added and the mixture left to stir for 1.5 hours at 65 °C. The mixture was cooled to room temperature, EtOAc (100 ml) and $H_2O$ (20 ml) were added. The aqueous layer was washed with 3x20 ml EtOAc. The organic layers were combined, concentrated and purified on silica (1:4 EtOOAc:hexane) yielding 761 mg of **10**. Yield: 61%. **$^1$H NMR** (400 MHz, CDCl3) δ 6.82 (d, J = 8.0 Hz, 1H), 6.24 (d, J = 2.4 Hz, 1H), 6.15 (dd, J = 8.0, 2.4 Hz, 1H), 4.55 (s, 1H), 3.97 (d, J = 2.4 Hz, 2H), 3.34 – 3.18 (m, 2H), 2.69 (t, J = 6.5 Hz, 2H), 2.16 (t, J = 2.4 Hz, 1H), 2.02 – 1.93 (m, 2H). **$^{13}$C NMR** (101 MHz, CDCl3) δ 154.73, 145.58, 129.78, 116.47, 104.12, 99.30, 79.48, 71.75, 49.12, 40.76, 26.91, 22.54.

**1-ethyl-7-methoxy-2,2,4-trimethyl-1,2-dihydroquinoline (11)**

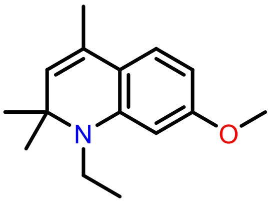


7-methoxy-2,2,4-trimethyl-1,2-dihydroquinoline (2 g, 9.8 mmol) and $K_2CO_3$ (2.12 g, 15.4 mmol) were added to 35 ml MeCN. Iodoethane (2 ml, 24.8 mmol) was added and the mixture was left stirring overnight at 90 °C. The mixture was filtered, concentrated and purified on silica (1:19 EtOAc:cyclohexane) yielding 850 mg of **11**. Yield: 37%. **$^1$H NMR** (400 MHz, $CDCl_3$) δ 6.96 (d, *J* = 8.3 Hz, 1H), 6.14 (dd, *J* = 8.3, 2.4 Hz, 1H), 6.08 (d, *J* = 2.4 Hz, 1H), 5.10 (q, *J* = 1.4 Hz, 1H), 3.79 (s, 3H), 3.30 (d, *J* = 7.1 Hz, 2H), 1.94 (d, *J* = 1.4 Hz, 3H), 1.30 (s, 6H), 1.20 (t, *J* = 7.0 Hz, 3H). **$^{13}$C NMR** (101 MHz, CDCl3) δ 160.53, 144.98, 127.35, 126.99, 124.44, 116.60, 98.76, 97.51, 56.89, 55.09, 38.19, 18.76, 14.25.

**(E)-1-ethyl-7-methoxy-2,2,4-trimethyl-6-((4-nitrophenyl)diazenyl)-1,2-dihydroquinoline (12)**

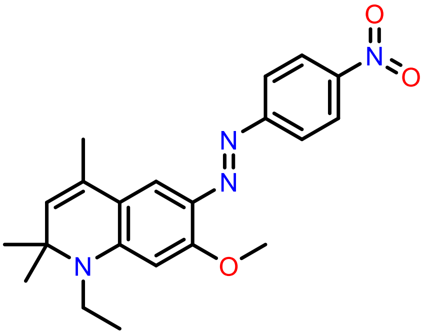


**11** (200 mg, 0.86 mg) was dissolved in 8 ml MeOH. A suspension of 4-nitrobenzenediazonium tetrafluoroborate (215 mg, 0.91 mmol) in 1 ml of a 10 % $H_2SO_4$ in $H_2O$ was added dropwise, the resulting solution was stirred for 1 hour and then cooled on ice before adding 0.16 ml of a 30% $NH_4OH$ solution. The precipitate was isolated, washed with 300 ml of $H_2O$, and then purified on silica (1:4 EtOAc:cyclohexane) yielding 286 mg of **12**. Yield: 87%. **$^1$H NMR** (400 MHz, CDCl3) δ 8.29 (d, J = 9.0 Hz, 2H), 7.87 (d, J = 9.0 Hz, 2H), 7.66 (s, 1H), 6.07 (s, 1H), 5.27 (d, J = 1.5 Hz, 1H), 4.04 (s, 3H), 3.51 (q, J = 7.1 Hz, 2H), 2.02 (d, J = 1.4 Hz, 3H), 1.41 (s, 6H), 1.33 (t, J = 7.0 Hz, 3H). **$^{13}$C NMR** (101 MHz, $CDCl_3$) δ 161.20, 157.97, 150.28, 146.76, 133.44, 128.54, 127.04, 124.83, 122.51, 116.48, 112.50, 93.24, 77.48, 77.16, 76.84, 58.78, 56.46, 39.18, 29.56, 27.07, 18.92, 13.98, 0.14.

**Oxazine alkyne (13)**

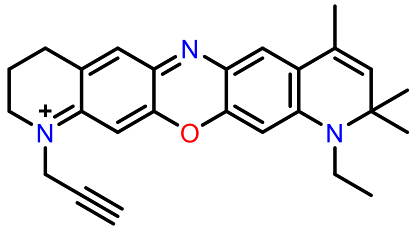


**10** (24.5 mg, 0.13 mmol) and **12** (50 mg, 0.13 mmol) were dissolved in an aqueous 1.5% HCl solution. The solution was heated to 80 °C for 4 hours, cooled and concentrated followed b purification on silica (1:19 MeOH:$CHCl_3$) yielding 42 mg of **13** which was not pure by NMR. **$^1$H NMR** (400 MHz, CDCl3) δ 7.47 (s, 1H), 7.36 (s, 1H), 7.28 (s, 1H), 6.47 (s, 1H), 5.53 (d, J = 1.5 Hz, 1H), 4.27 – 4.18 (m, 1H), 3.82 (q, J = 5.7 Hz, 2H), 3.74 (s, 2H), 3.73 – 3.66 (m, 3H), 3.25 (d, J = 7.5 Hz, 3H), 2.88 (t, J = 6.3 Hz, 3H), 2.29 (s, 2H), 2.14 – 2.07 (m, 3H), 2.06 (s, 3H), 2.04 – 1.95 (m, 4H), 1.52 (s, 6H).

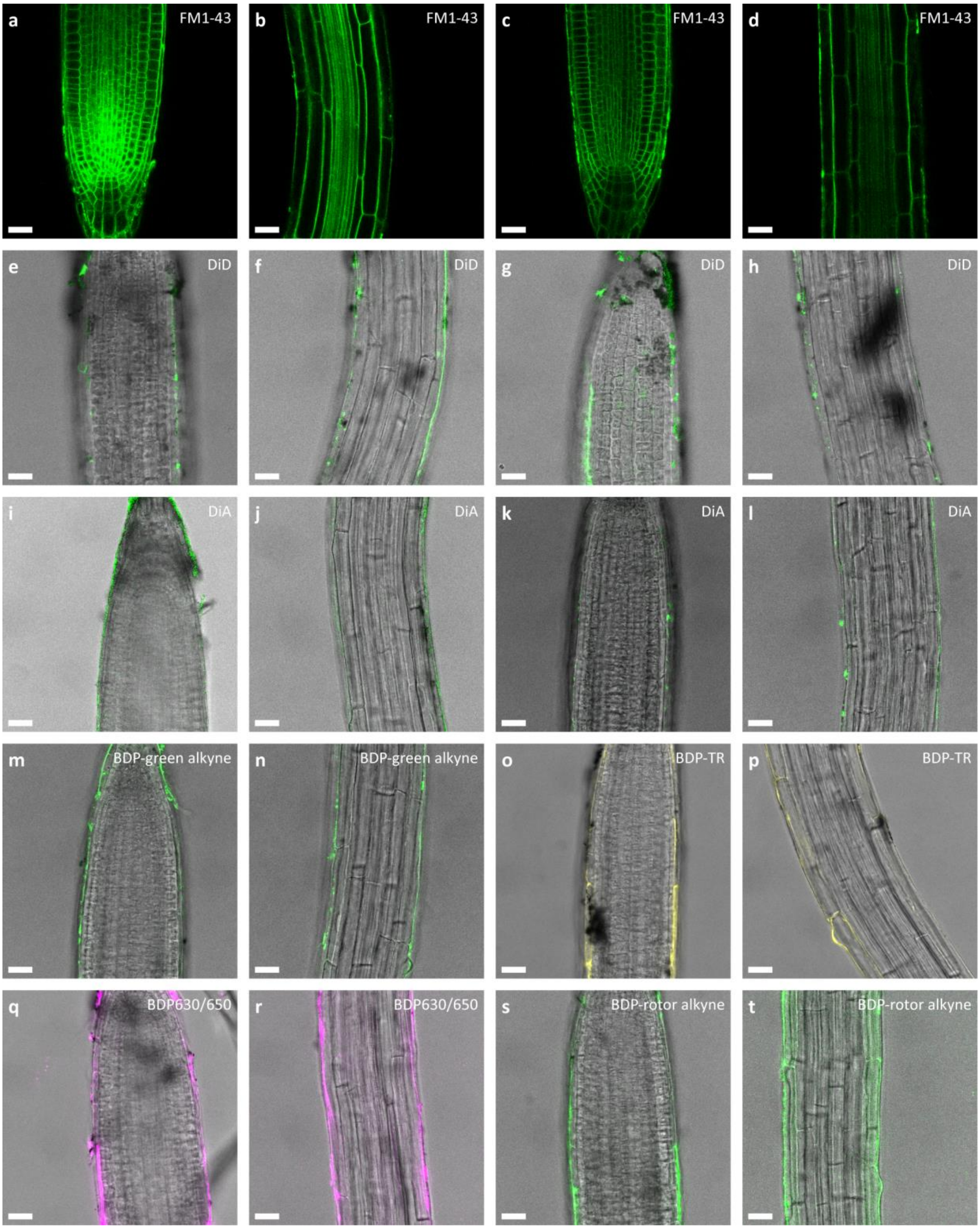


**Fig. S1.** Staining of Arabidopsis with commonly used mammalian membrane probes and LipoTag dye precursors.

**a-d,** Staining of different regions of Arabidopsis roots with 10 µM (**a**,**b**) and 1 µM (**c**,**d**) FM1-43. **e-h,** Staining of different regions of Arabidopsis roots with 10 µM (**e**,**f**) and 1 µM (**g**,**h**) DiD. **i-l,** Staining of different regions of Arabidopsis roots with 10 µM (**i**,**j**) and 1 µM (**k**,**l**) DiA . **m-n,** Staining of Arabidopsis root with 1 µM LipoTag-Green precursor BDP-alkyne green (**6**). **o-p,** Staining of Arabidopsis root with 1 µM LipoTag-Orange precursor BDP-TR alkyne. **q-r,** Staining of Arabidopsis root with 1 µM LipoTag-Red precursor BDP630/650 alkyne. **s-t,** Staining of Arabidopsis root with 1 µM LipoTag-BDP precursor BDP-alkyne green (**6**). **m-n,** Staining of Arabidopsis root with 1 µM LipoTag-Green precursor BDP-rotor alkyne (**7**).Scale bars represent 25 µm.

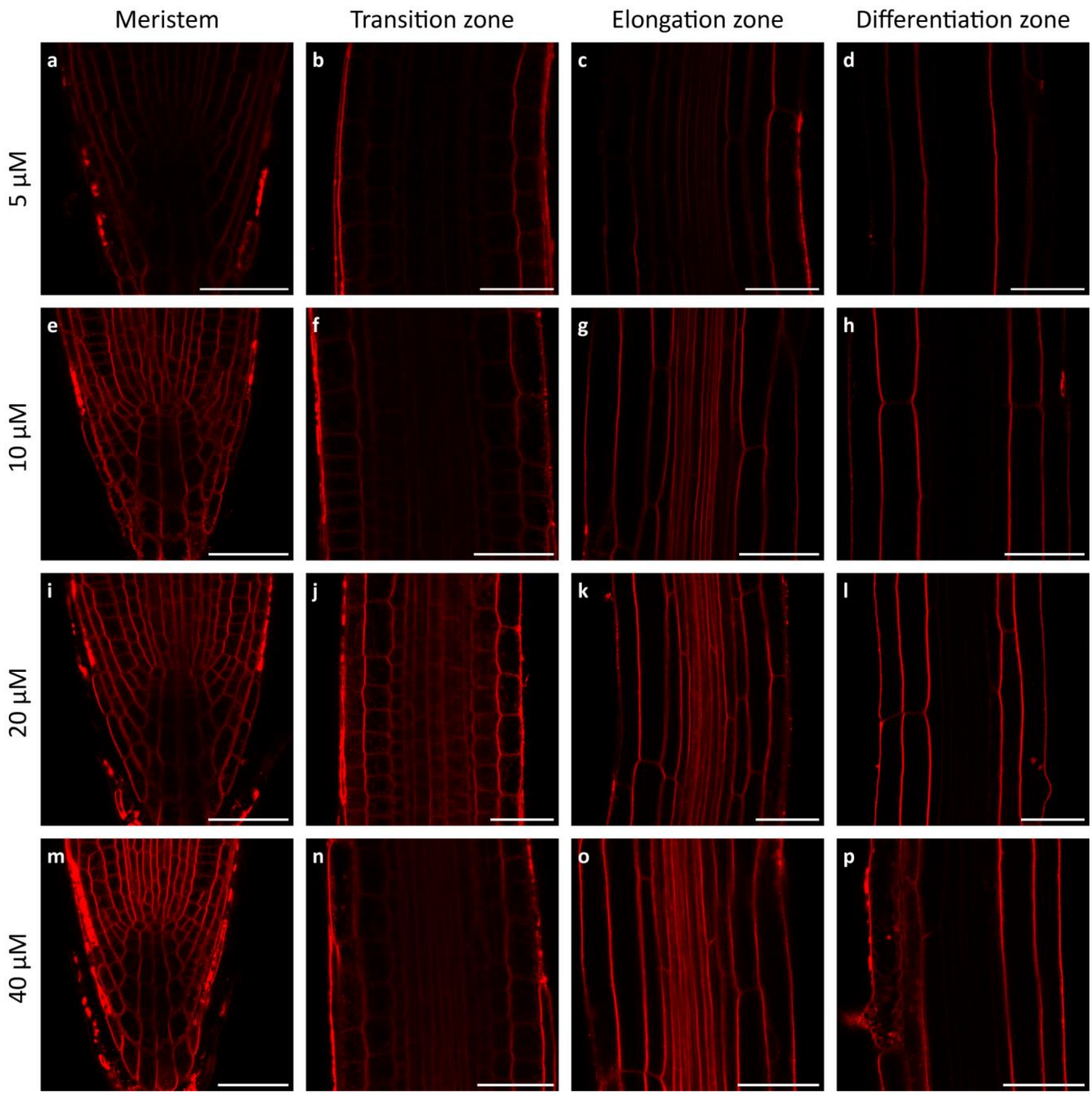


**Fig. S2.** Concentration range FM4-64 in different Arabidopsis root zones.

Concentration range with FM4-64 for different root tissues with 5 µM in 0.5x MS (**a-d**), 10 µM in 0.5x MS (**e-h**), 20 µM in 0.5x MS (**i-l**), and 40 µM in 0.5x MS (**m-p**) for 20 minutes. All measurements were performed and processed the same way. Scale bar represents 40 µm.

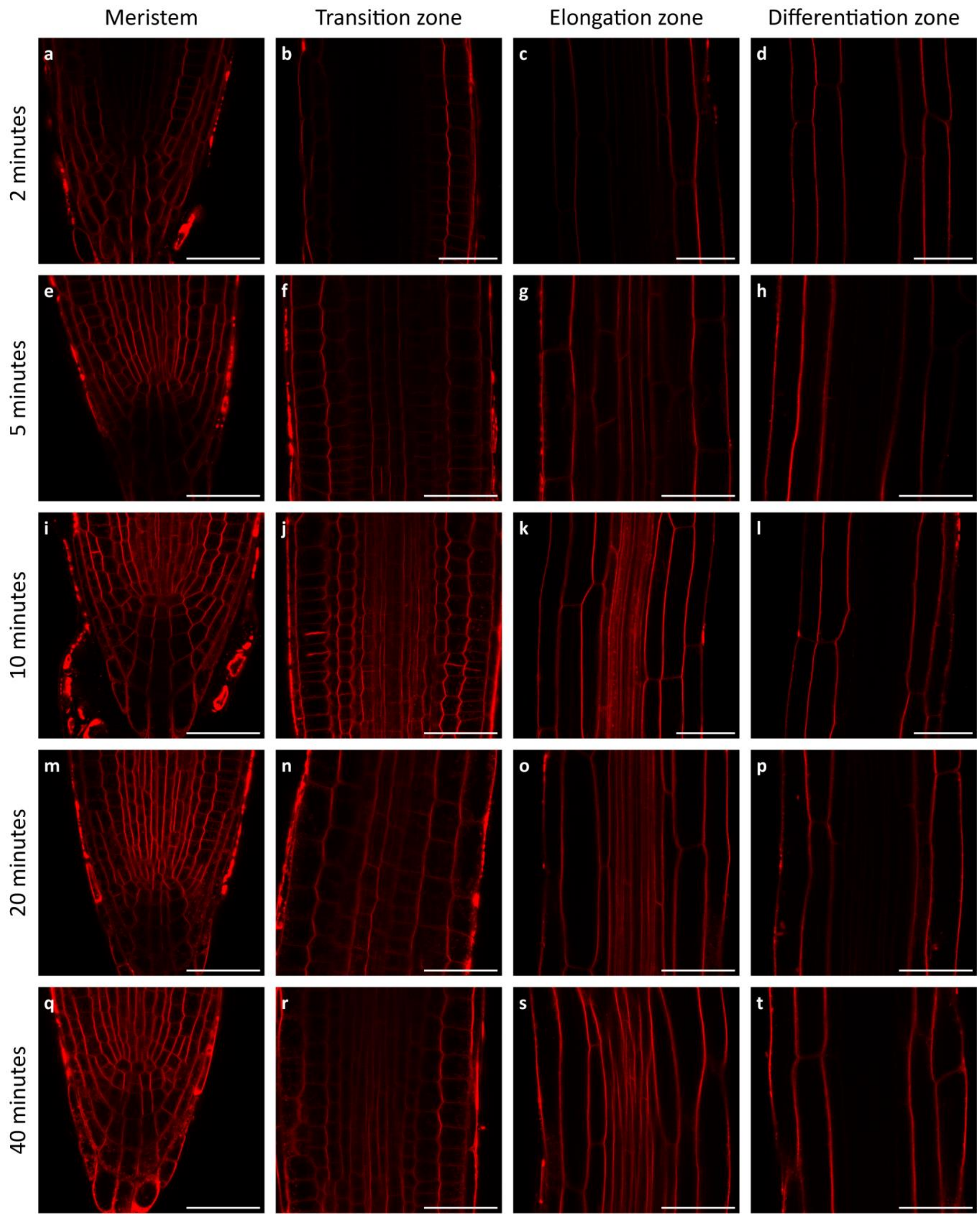


**Fig. S3.** Penetration depth of FM4-64 in different tissues over time.

Time series for FM4-64 at 20 µM in 0.5x MS for different root tissues at time 2 minutes (**a-d**), 5 minutes (**d-h**), 10 minutes (**i-l**), 20 minutes (**m-p**), 40 minutes (**q-t**). All measurements were performed and processed the same way. Scale bar represents 40 µm.

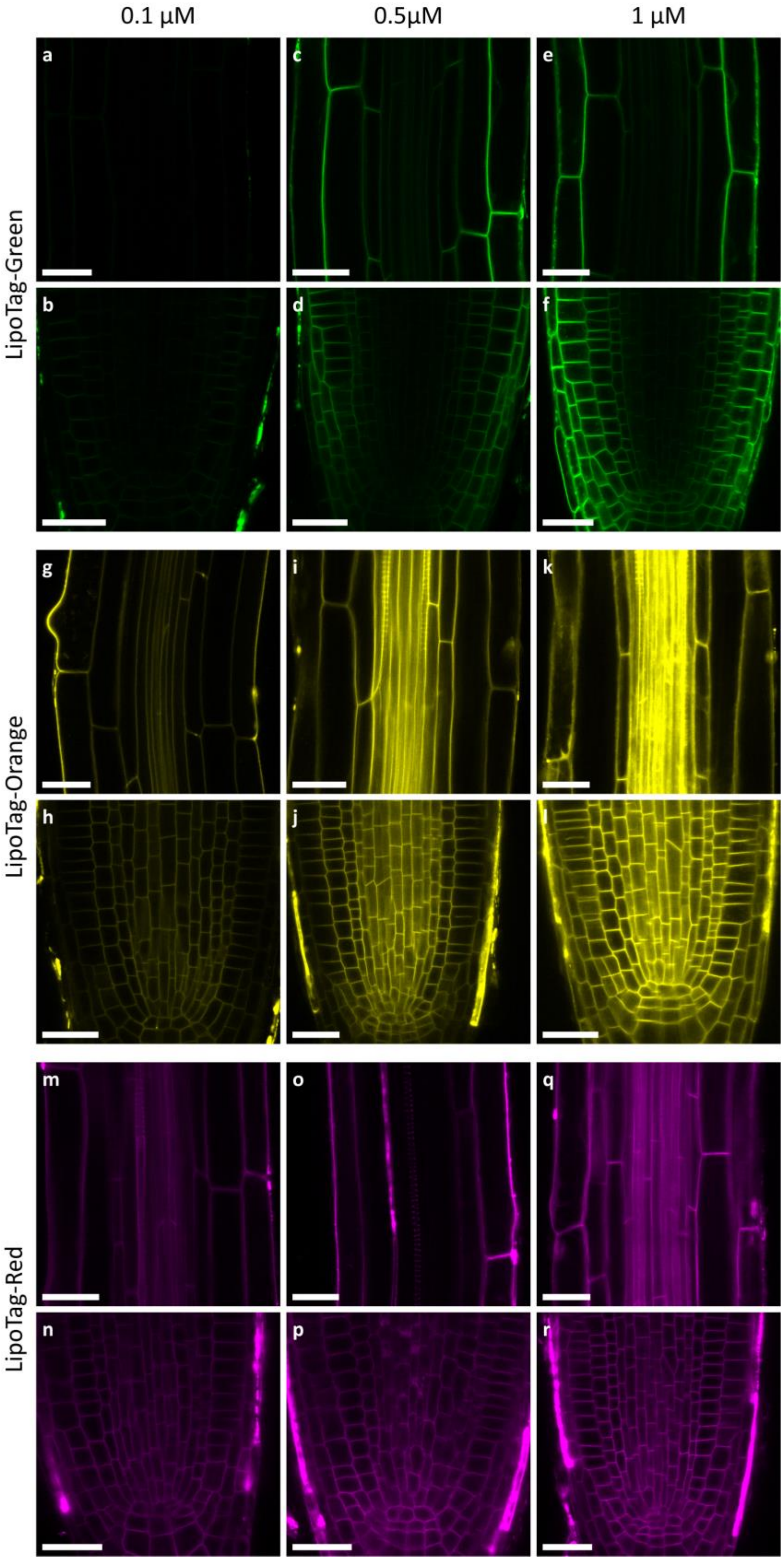


**Fig. S4.** Effect of staining concentration of non-functional LipoTag probes on tissue penetration after 30 minutes.

**a-f,** Staining of LipoTag-Green in the elongation zone (**a,c,e**) and the root tip (**b,d,f**) after 30 minutes of incubation. **g-l,** Staining of LipoTag-Orange in the elongation zone (**g,i,k**) and the root tip (**h,j,l**) after 30 minutes of incubation. **m-r,** Staining of LipoTag-Red in the elongation zone (**m,o,q**) and the root tip (**n,p,r**) after 30 minutes of incubation. All measurements were performed and processed the same way. Scale bar represents 25 µm.

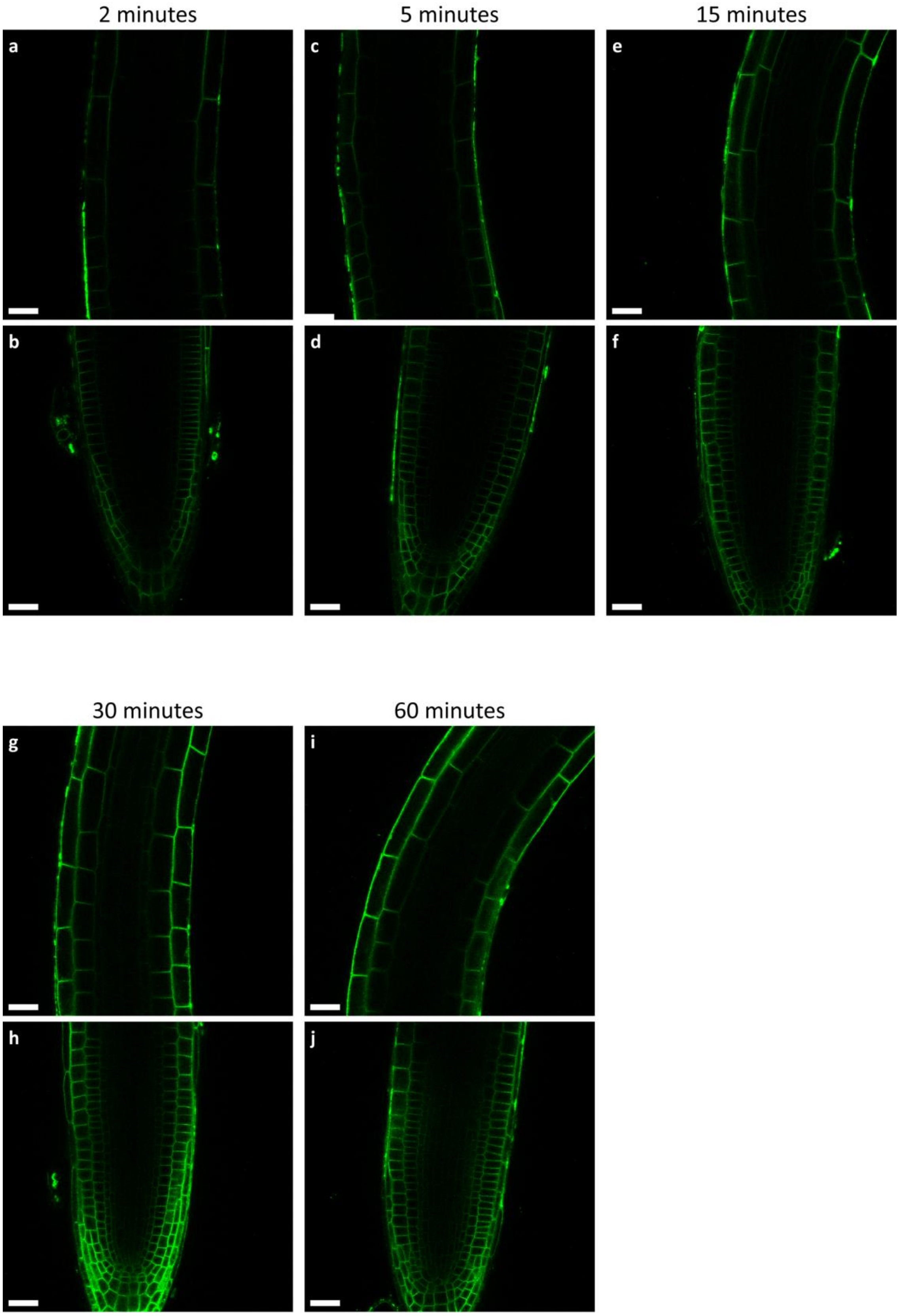


**Fig. S5.** Staining kinetics of LipoTag-Green in the root tip and elongation zone.

Images of Arabidopsis root stained with 0.5 μM LipoTag-Green after 2 minutes (**a,b**), 5 minutes (**c,d**), 15 minutes (**e,f**), 30 minutes (**g,h**) and 60 minutes (**i,j**). All measurements were performed and processed the same way. Scale bar represents 25 μm.

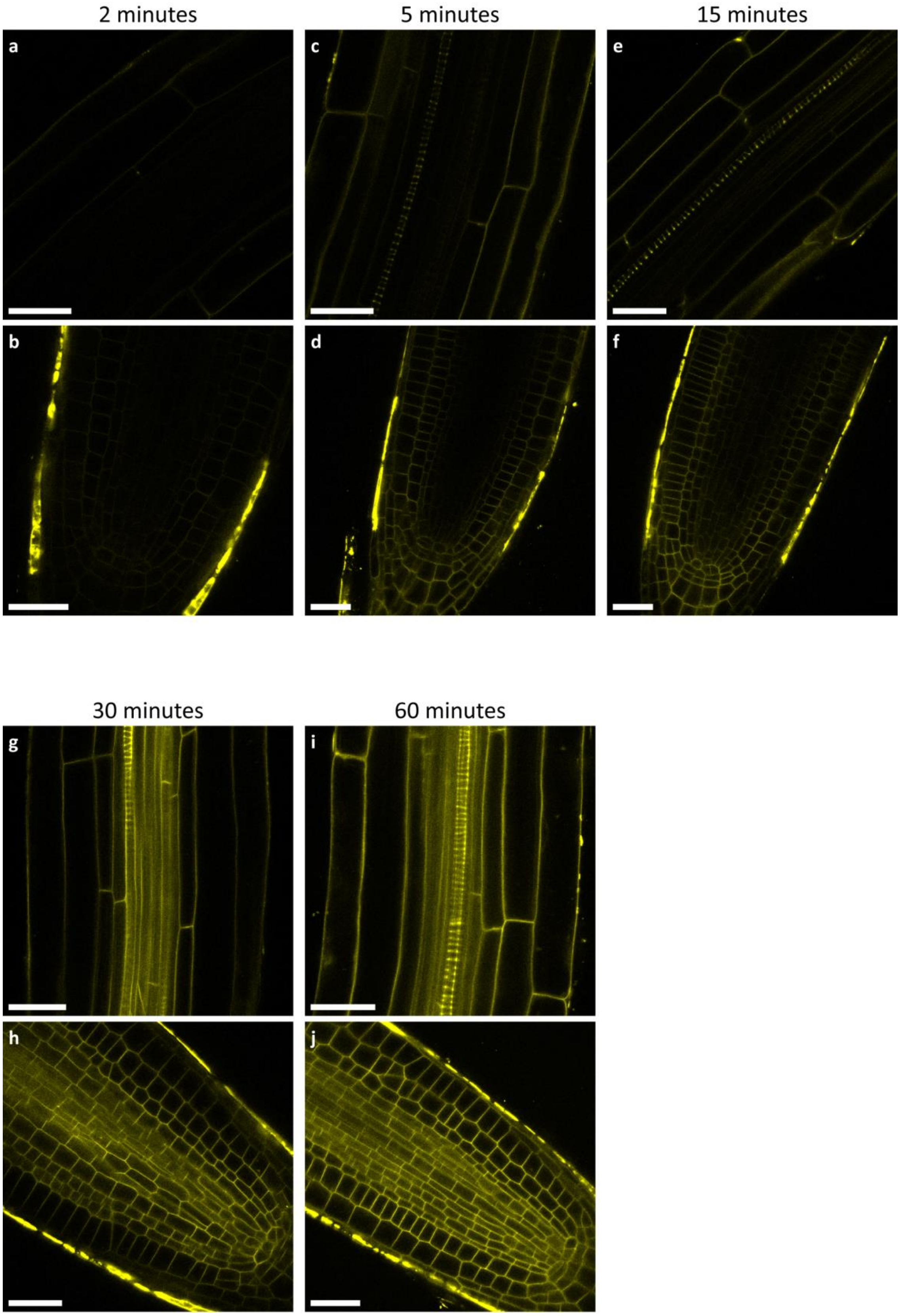


**Fig. S6.** Staining kinetics of LipoTag-Orange in the root tip and elongation zone.

Images of Arabidopsis root stained with 0.5 μM LipoTag-Orange after 2 minutes (**a,b**), 5 minutes (**c,d**), 15 minutes (**e,f**), 30 minutes (**g,h**) and 60 minutes (**i,j**). All measurements were performed and processed the same way. Scale bar represents 25 μm.

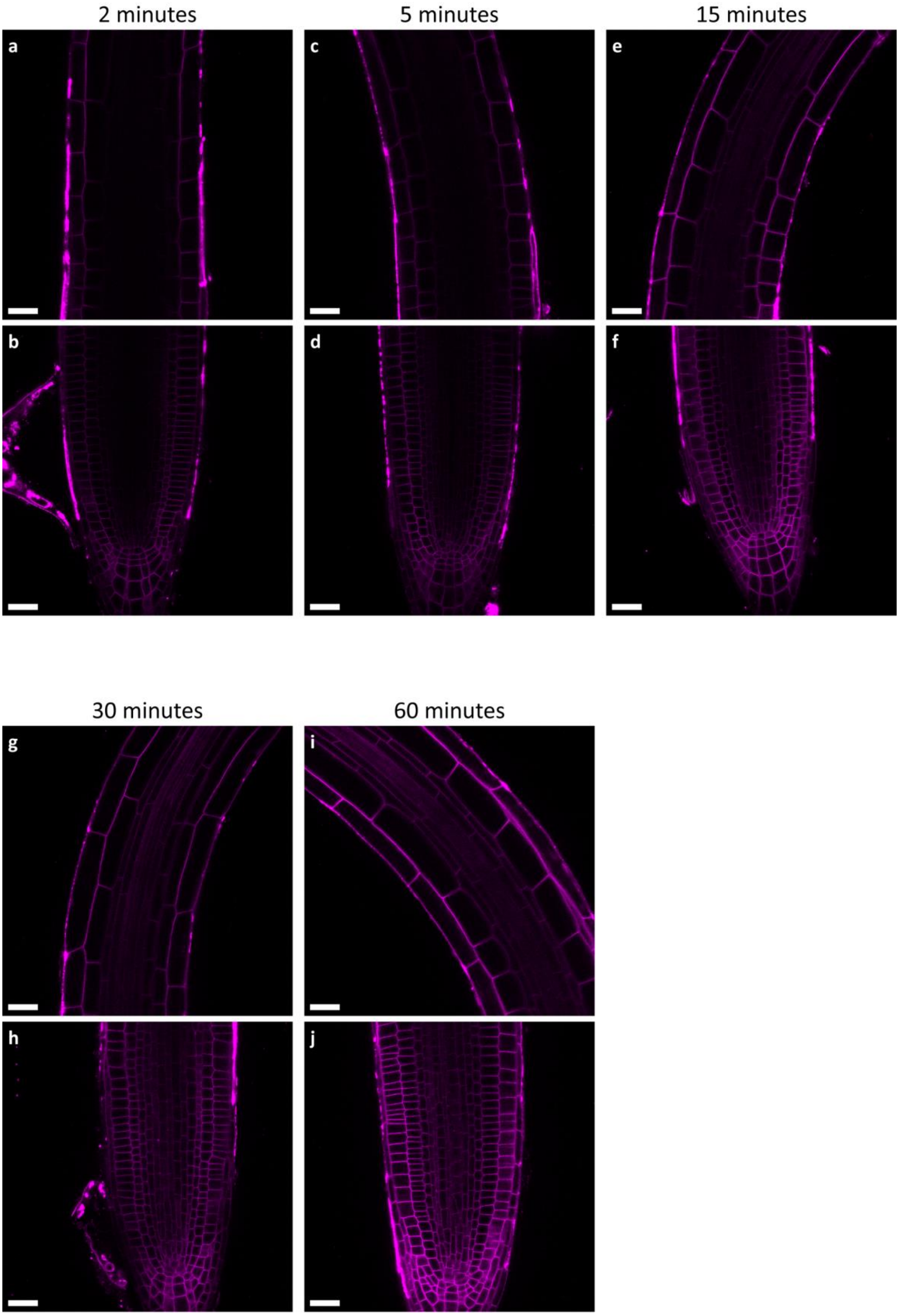


**Fig. S7.** Staining kinetics of LipoTag-Red in the root tip and elongation zone.

Images of Arabidopsis root stained with 0.5 µM LipoTag-Red after 2 minutes (**a,b**), 5 minutes (**c,d**), 15 minutes (**e,f**), 30 minutes (**g,h**) and 60 minutes (**i,j**). All measurements were performed and processed the same way. Scale bar represents 25 µm.

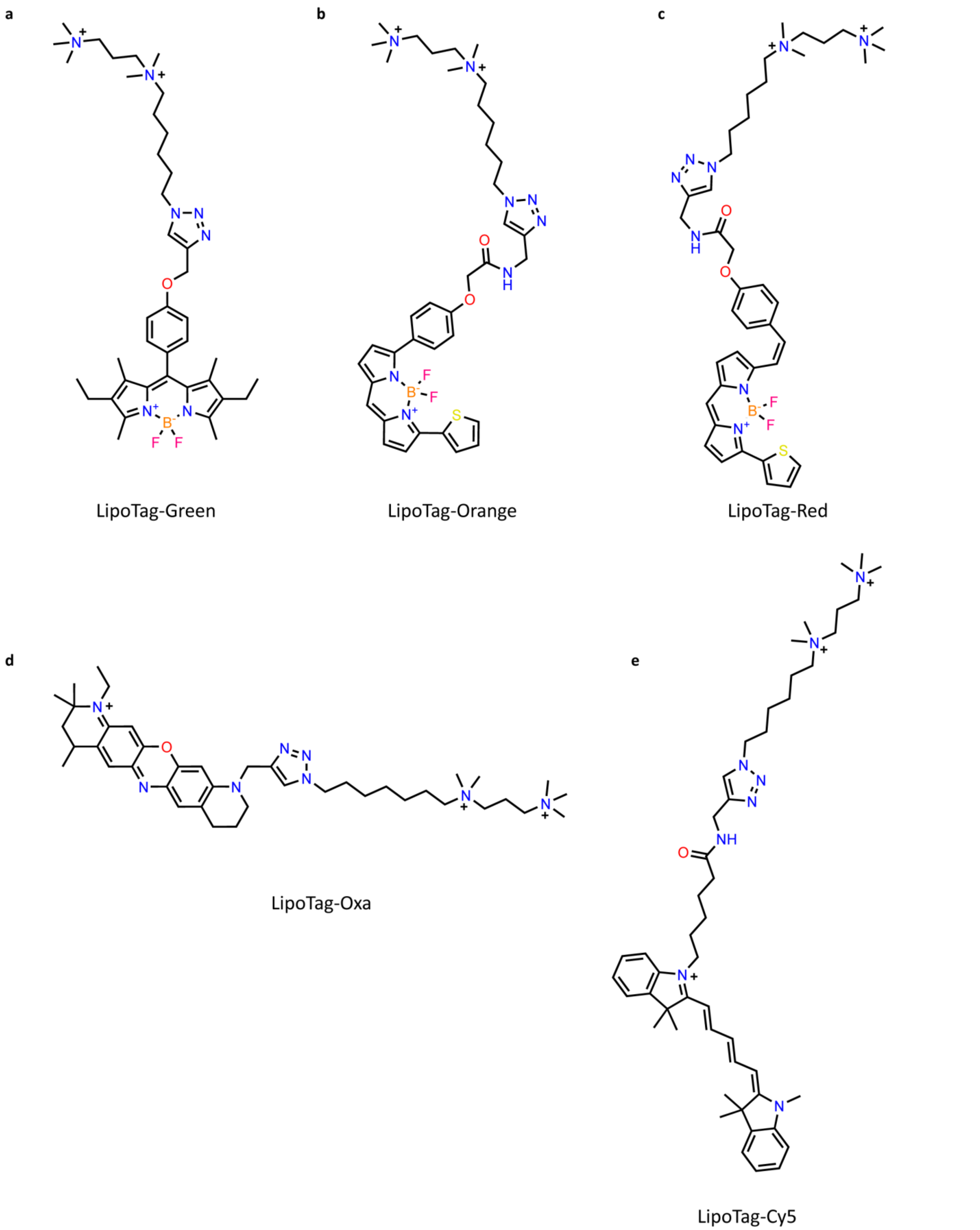


**Fig. S8.** Chemical structures of non-functional LipoTag probes.

Chemical structure of LipoTag-Green (**a**), LipoTag-Orange (**b**), LipoTag-Red (**c**), Lipotag-Oxa (**d**) and LipoTag-Cy5 (**e**)

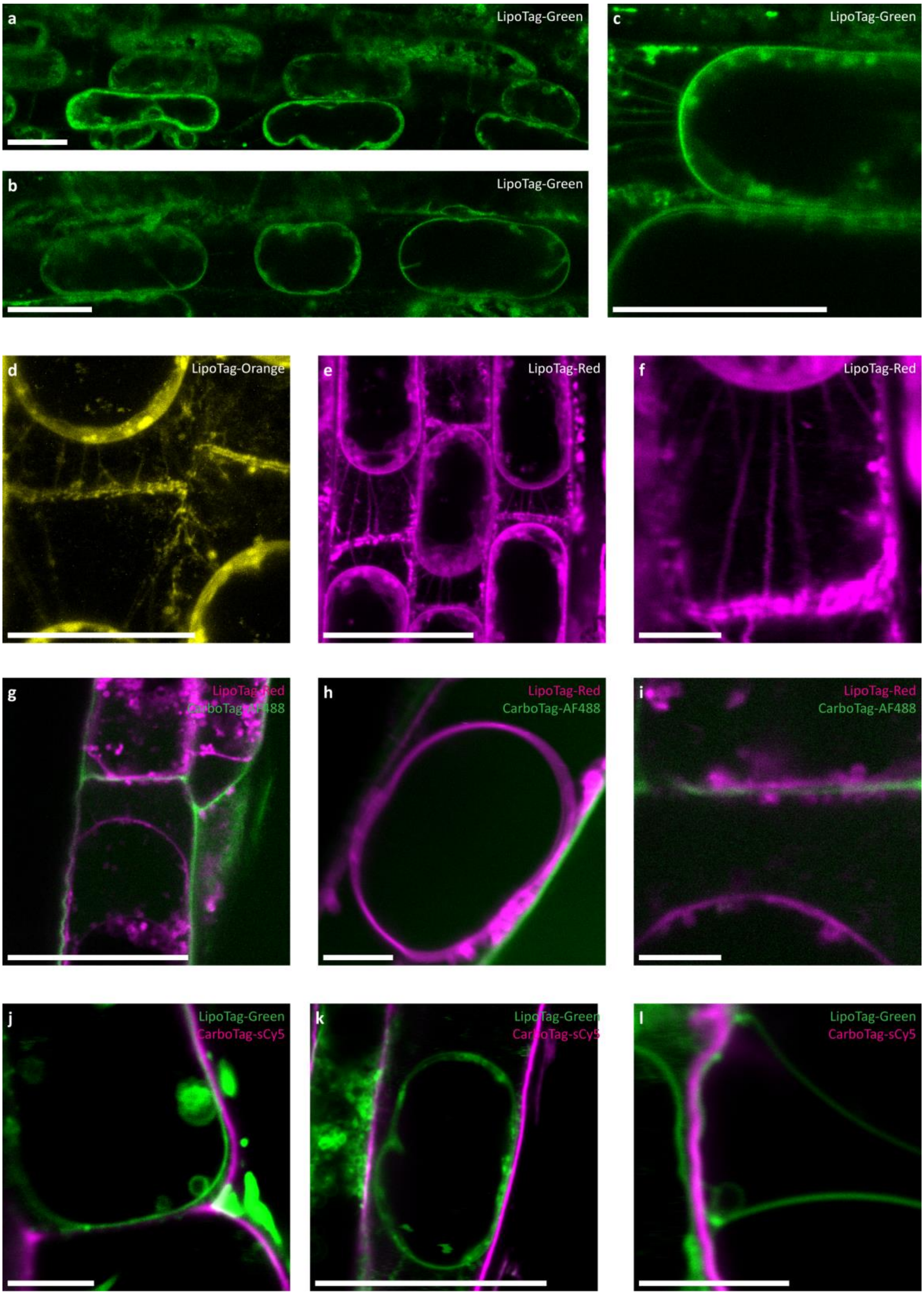


**Fig. S9.** Plasmolysis of LipoTag stained Arabidopsis roots.

Arabidopsis roots stained with LipoTag-Green (**a-c**), LipoTag-Orange (**b**) and LipoTag-Red (**e-f**) after gradual plasmolysis with 0.3 M mannitol followed by 0.5 M mannitol. Besides the retracted membrane Hechtian strands and the Hechtian reticulum are visible. To confirm membrane localization CarboTag-AF488 (green) and CarboTag-sCy5 (magenta) were used to co-stain the cell wall during plasmolysis with LipoTag-Red (magenta) (**g-i**) and LipoTag-Green (green) (**j-l)** respectively. Scale bars represent 5 μm for **f, h, i, j** and **l,** all others represent 25 μm.

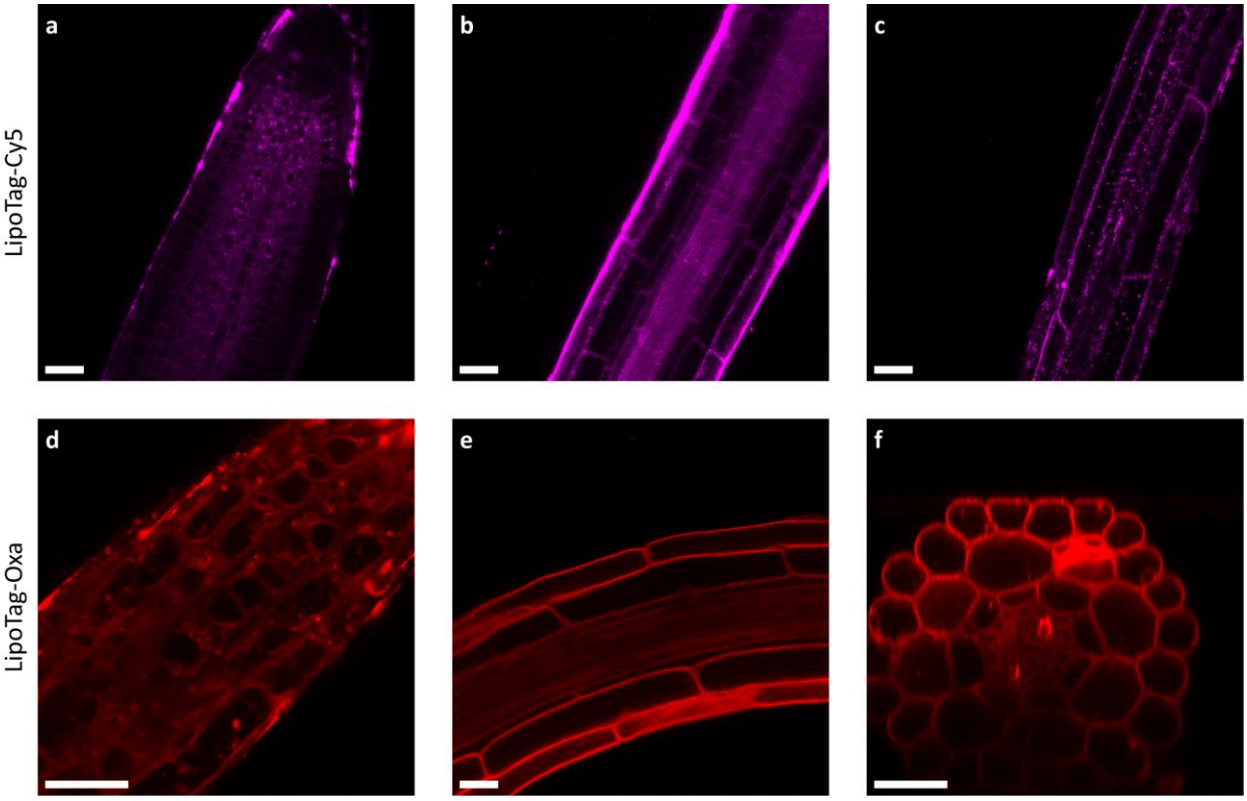


**Fig. S10.** Staining of Arabidopsis roots with LipoTag-Cy5 and LipoTag-Oxa.

Staining of Arabidopsis roots with 1 µM LipoTag-Cy5 (**a-c**) and 1 µM LipoTag-Oxa (**d**) and 0.1 µM LipoTag-Oxa (**e-f**). **f** is a xzy projection. Scale bars represent 25 µm.

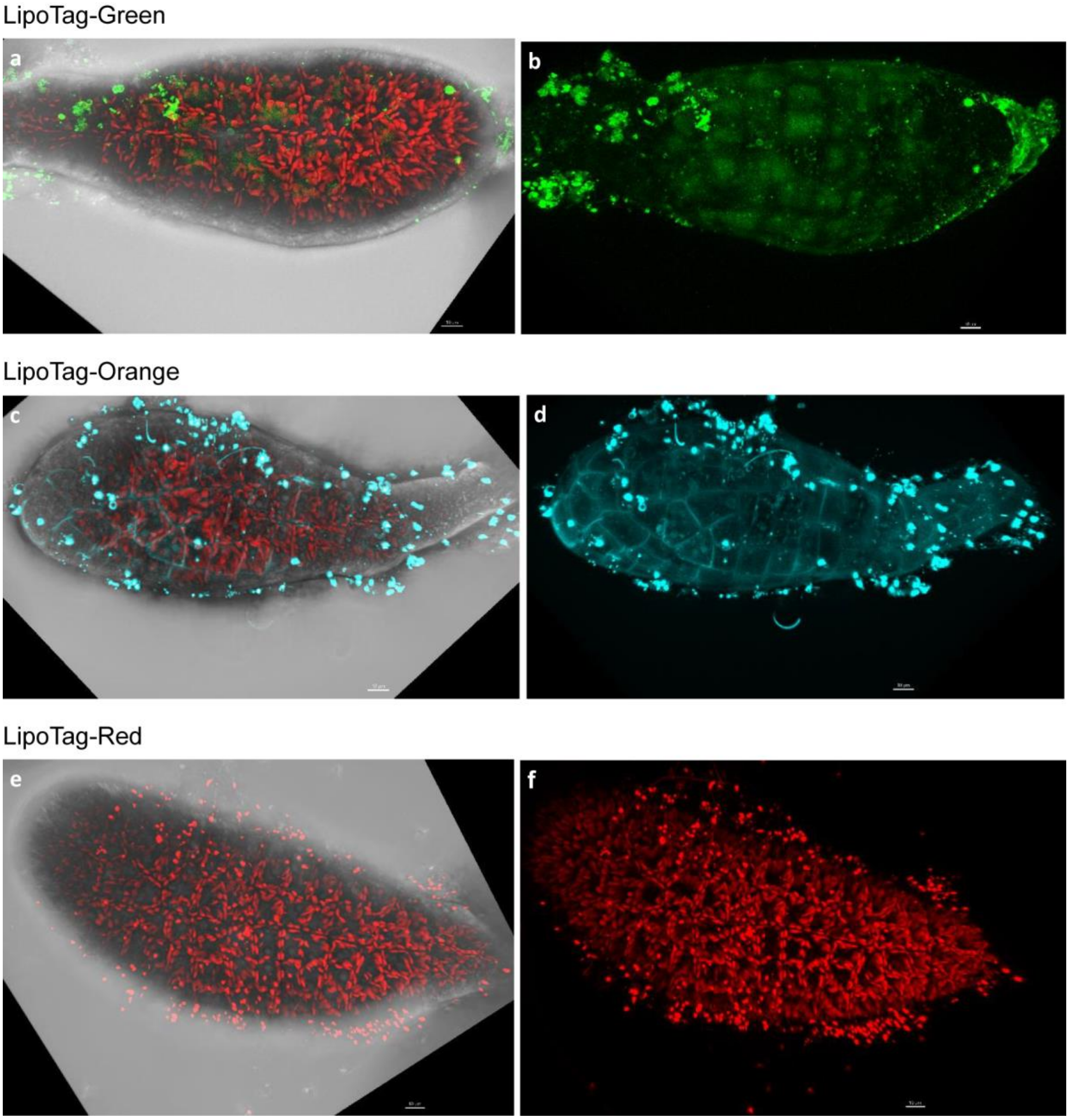


**Fig. S11.** Staining of *F. serratus* with LipoTag probes.

Images of *F. serratus* stained with LipoTag-Green (green) (**a-b**) with the brightfield overlay shown in **a**, LipoTag-Orange (cyan) (**c-d**) with the brightfield overlay shown in **c** and LipoTag-Red (red) (**e-f**) with the brightfield overlay shown in **e**. Chlorophyl autofluorescence is shown in red, scale bars represent 10 µm.

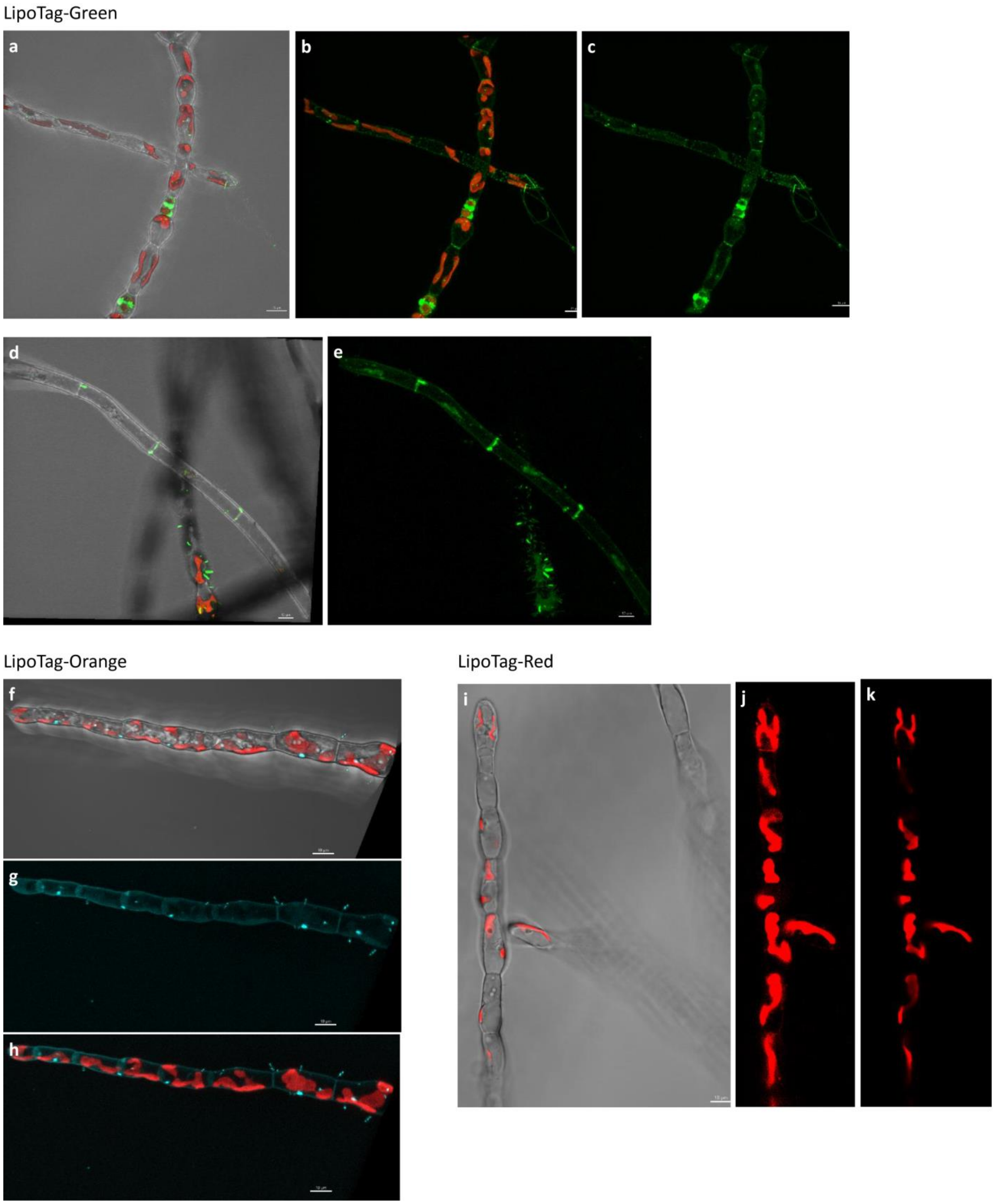


**Fig. S12.** Staining of *Ectocarpus sp.* with LipoTag probes.

Images of *Ectocarpus sp.* stained with LipoTag-Green (green) (**a-e**) with the brightfield overlay shown in **a** and d, LipoTag-Orange (cyan) (**f-h**) with the brightfield overlay shown in **f** and LipoTag-Red (red) (**i-k**) with the brightfield overlay shown in **i**. Chlorophyl autofluorescence is shown in red, scale bars represent 10 µm.

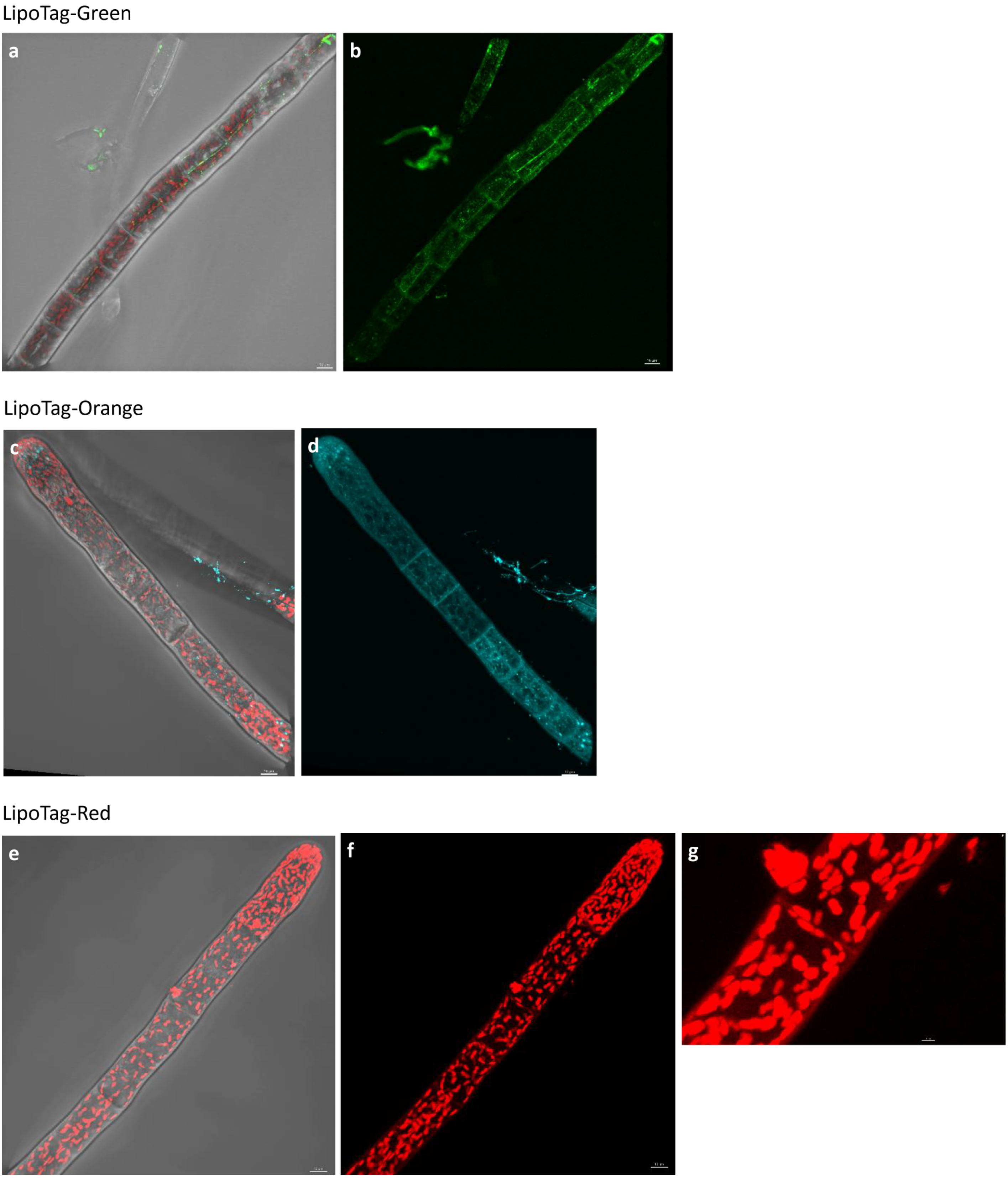


**Fig. S13.** Staining of *S. rigidula* with LipoTag probes.

Images of *S. rigidula* stained with LipoTag-Green (green) (**a-b**) with the brightfield overlay shown in **a**, LipoTag-Orange (cyan) (**c-d**) with the brightfield overlay shown in **c** and LipoTag-Red (red) (**e-g**) with the brightfield overlay shown in **e**. Chlorophyl autofluorescence is shown in red, scale bars represent 10 µm.

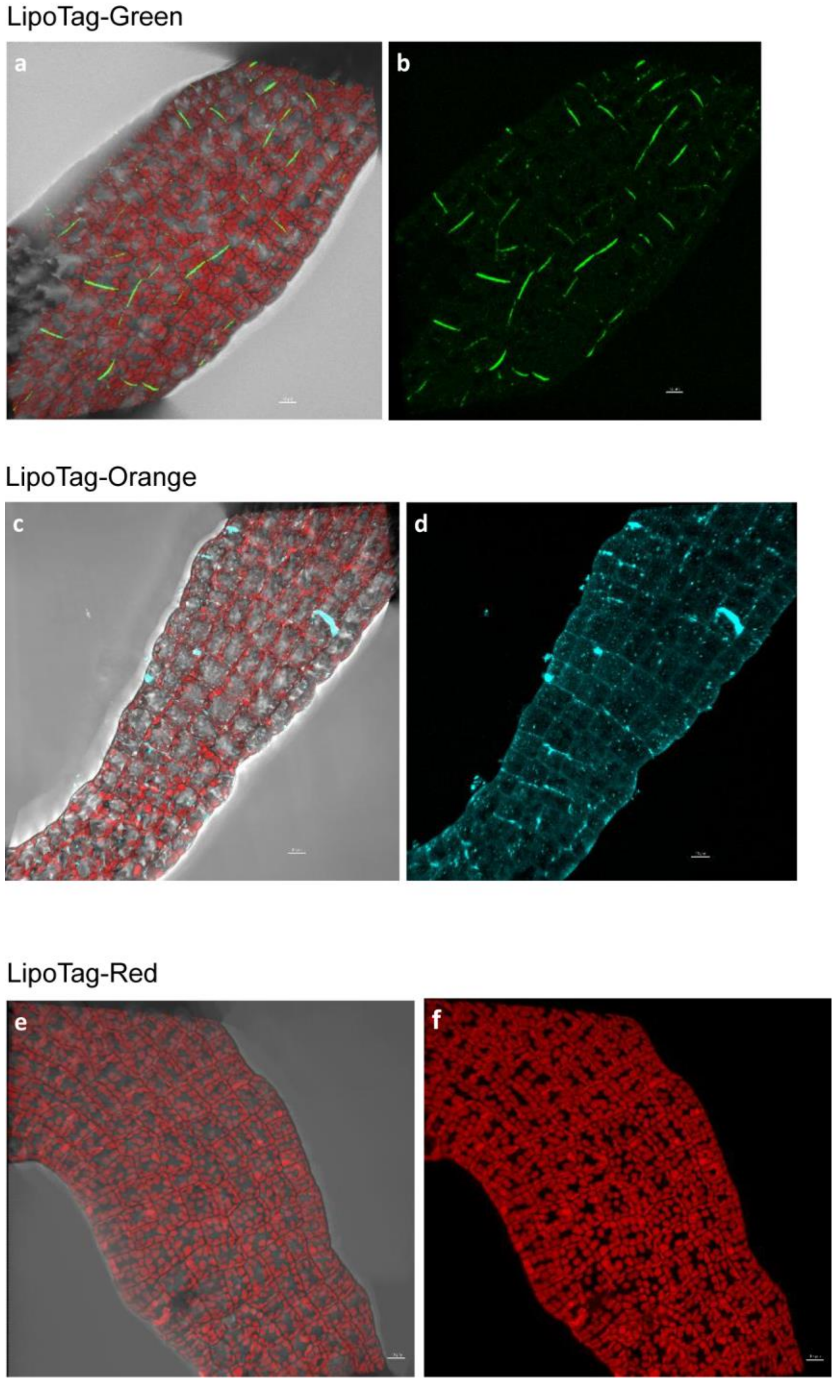


**Fig. S14.** Staining of *S. latissima* with LipoTag probes.

Images of *S. latissima* stained with LipoTag-Green (green) (**a-b**) with the brightfield overlay shown in **a**, LipoTag-Orange (cyan) (**c-d**) with the brightfield overlay shown in **c** and LipoTag-Red (red) (**e-f**) with the brightfield overlay shown in **e**. Chlorophyl autofluorescence is shown in red, scale bars represent 10 µm.

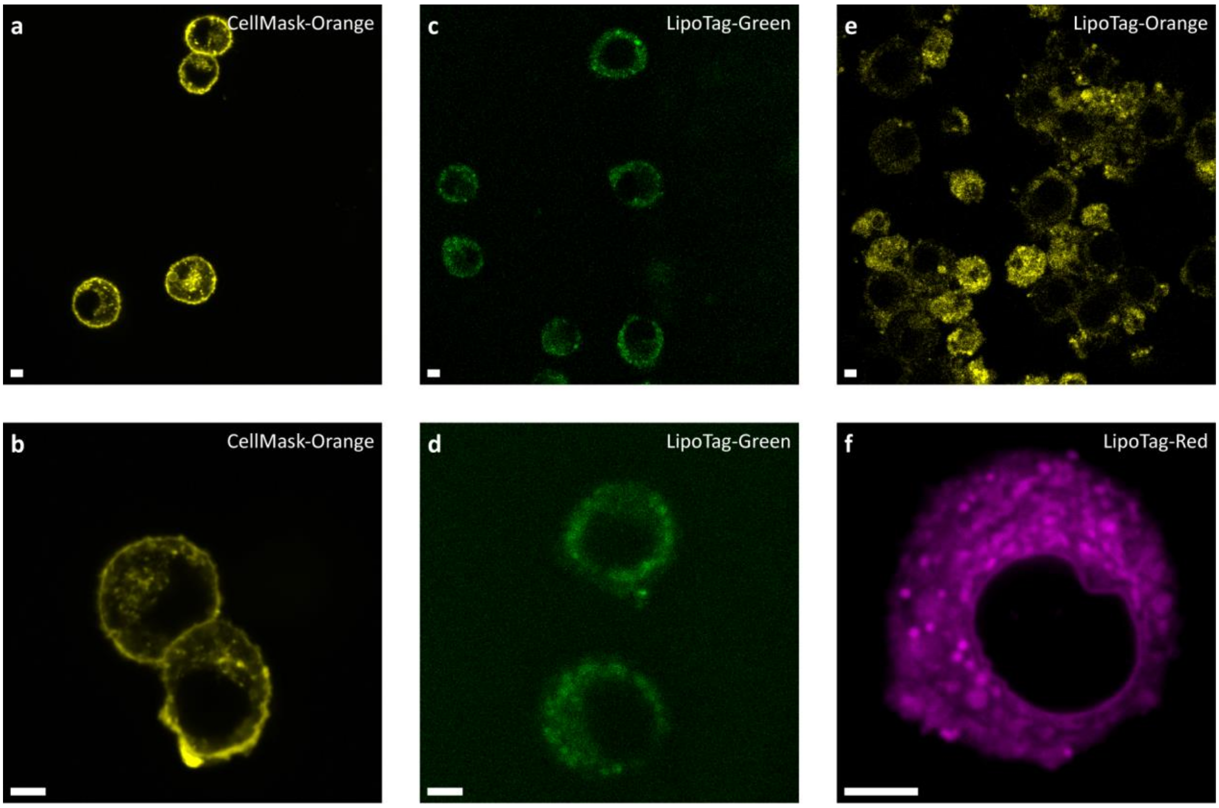


**Fig. S15.** Staining of Murine macrophage cells with LipoTag dyes.

Images of Murine cells stained with Cellmask-Orange (**a-b**), LipoTag-Green (**c-d**), LipoTag-Orange (**e**) and LipoTag-Red (**f**). Scale bars represent 5 µm.

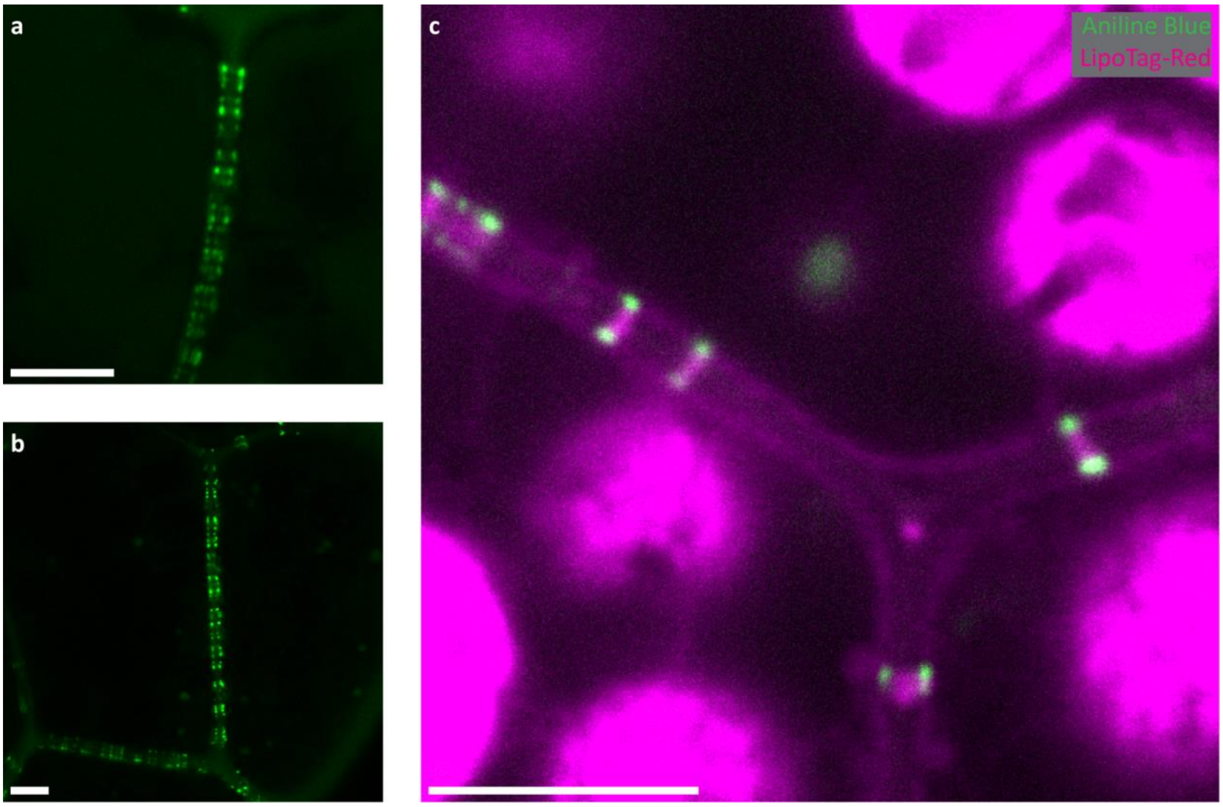


**Fig. S16.** Staining of plasmodesmata in Marchantia gemmae.

Staining of plasmodesmata in Marchantia gemmae with Aniline Blue (**a-b**) and gemmae co-stained with Aniline Blue (green) and LipoTag-Red (magenta) (**c**). Autofluorescence of chlorophyl is visible together with the outline of cell membranes and the plasmodesmata membrane. Scale bars represent 5 μm.

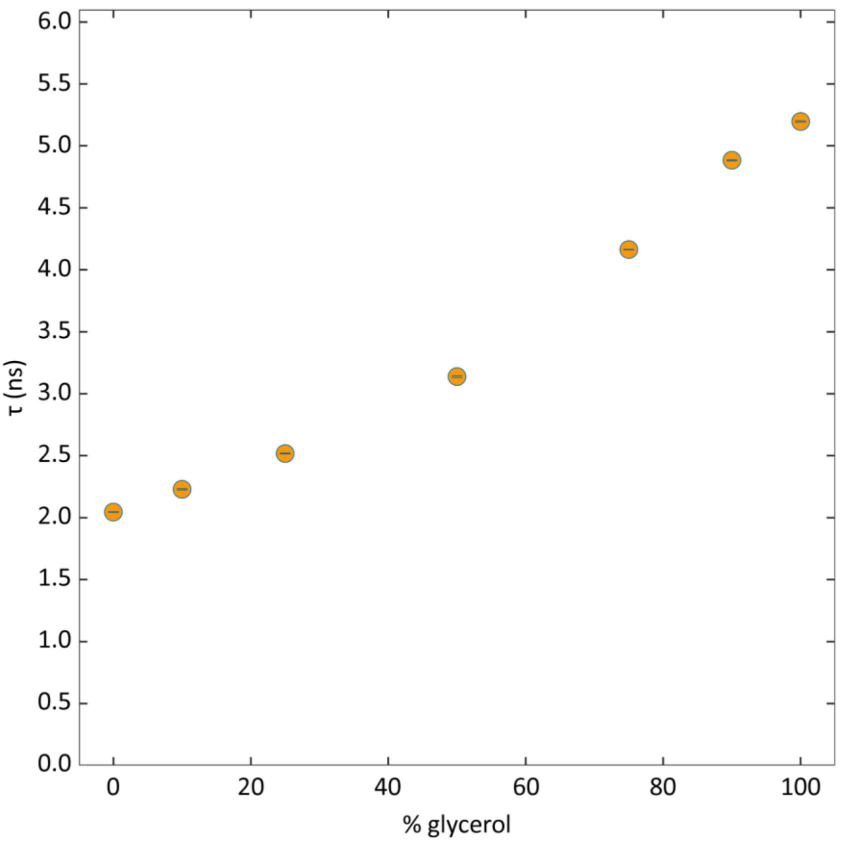


**Fig. S17.** Calibration of LipoTag-BDP.

Fluorescence lifetime of LipoTag-BDP in mixtures with varying glycerol concentration. All measurements were repeated 5 times, error bars fall within the point size.

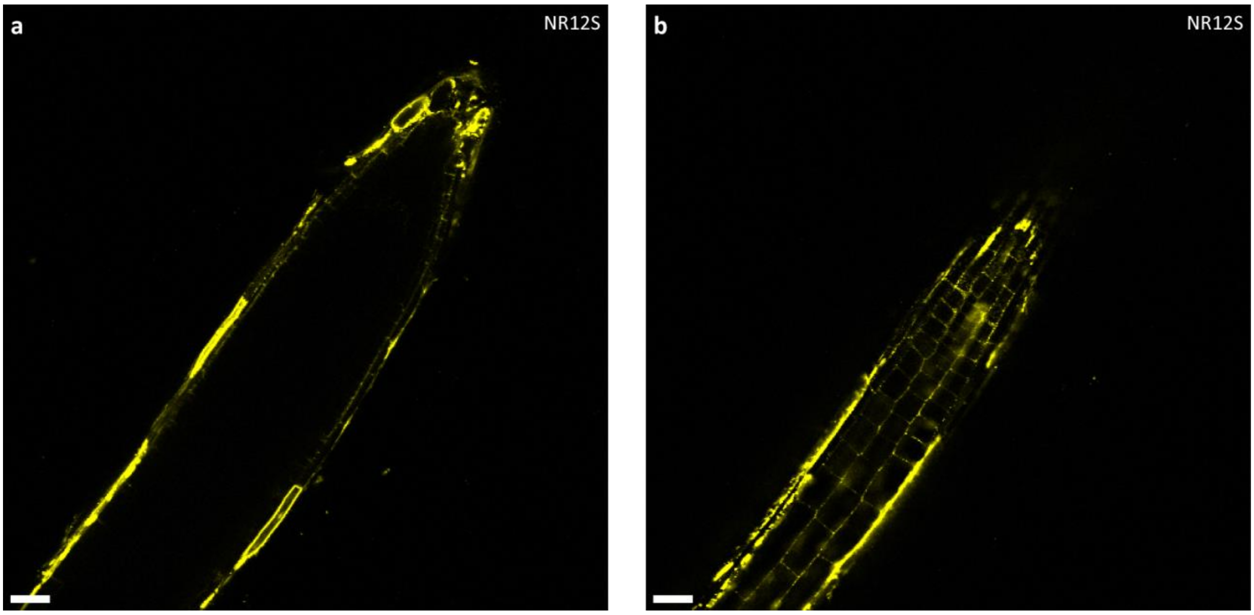


**Fig. S18.** Staining of Arabidopsis root with NR12S.

Staining of Arabidopsis root with 10 μM NR12S for 1 hour. Scale bars represent 25 μm.

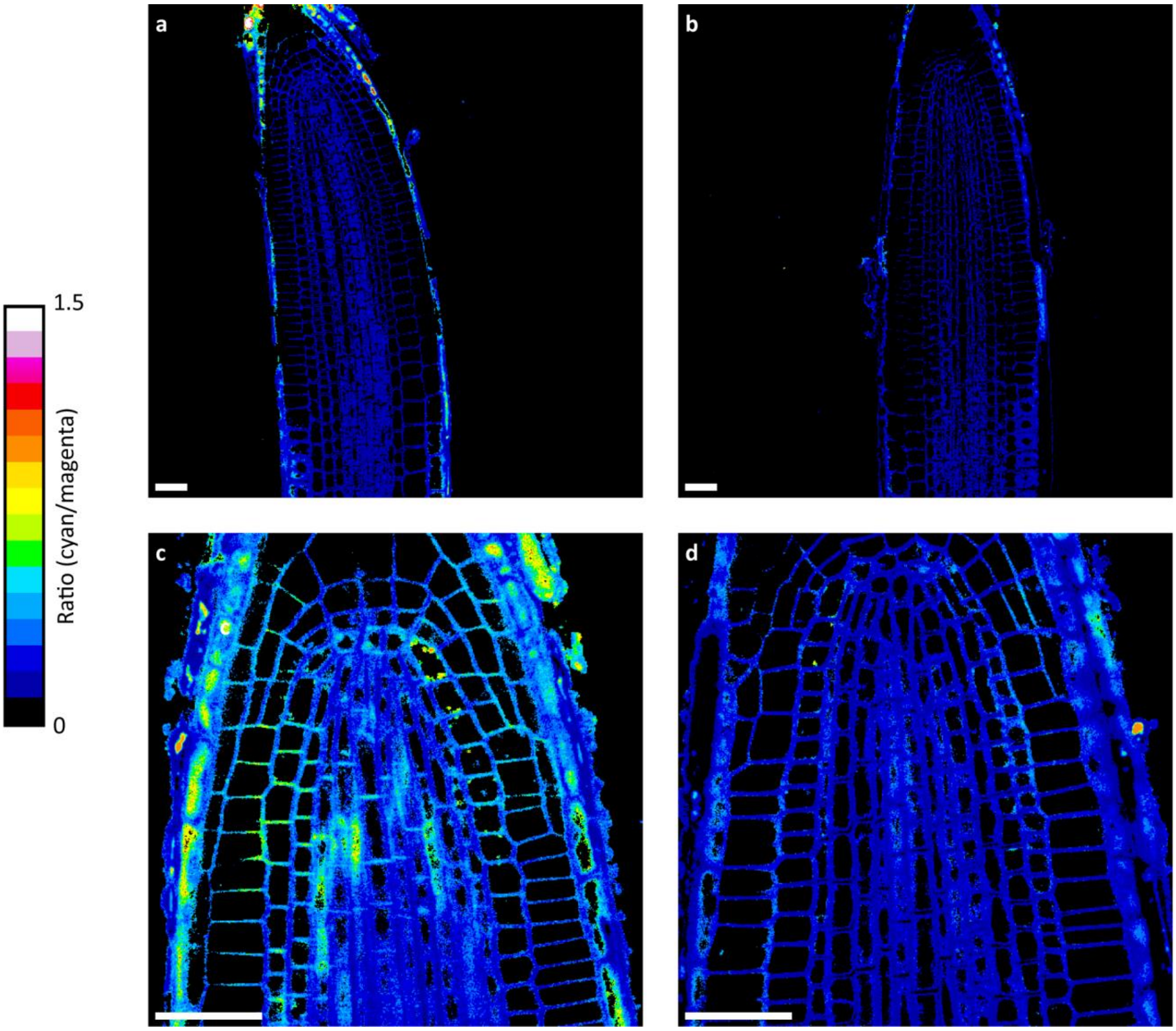


**Fig. S19.** Mock hemin treatment with LipoTag-Ox.

**a-d,** Ratiometric images of Arabidopsis root tips incubated for 1 hour with DMSO and ethanol (10 µl DMSO and 8 µl ethanol per 2 ml of 0.5x MS). Scale bars represent 25 µm.

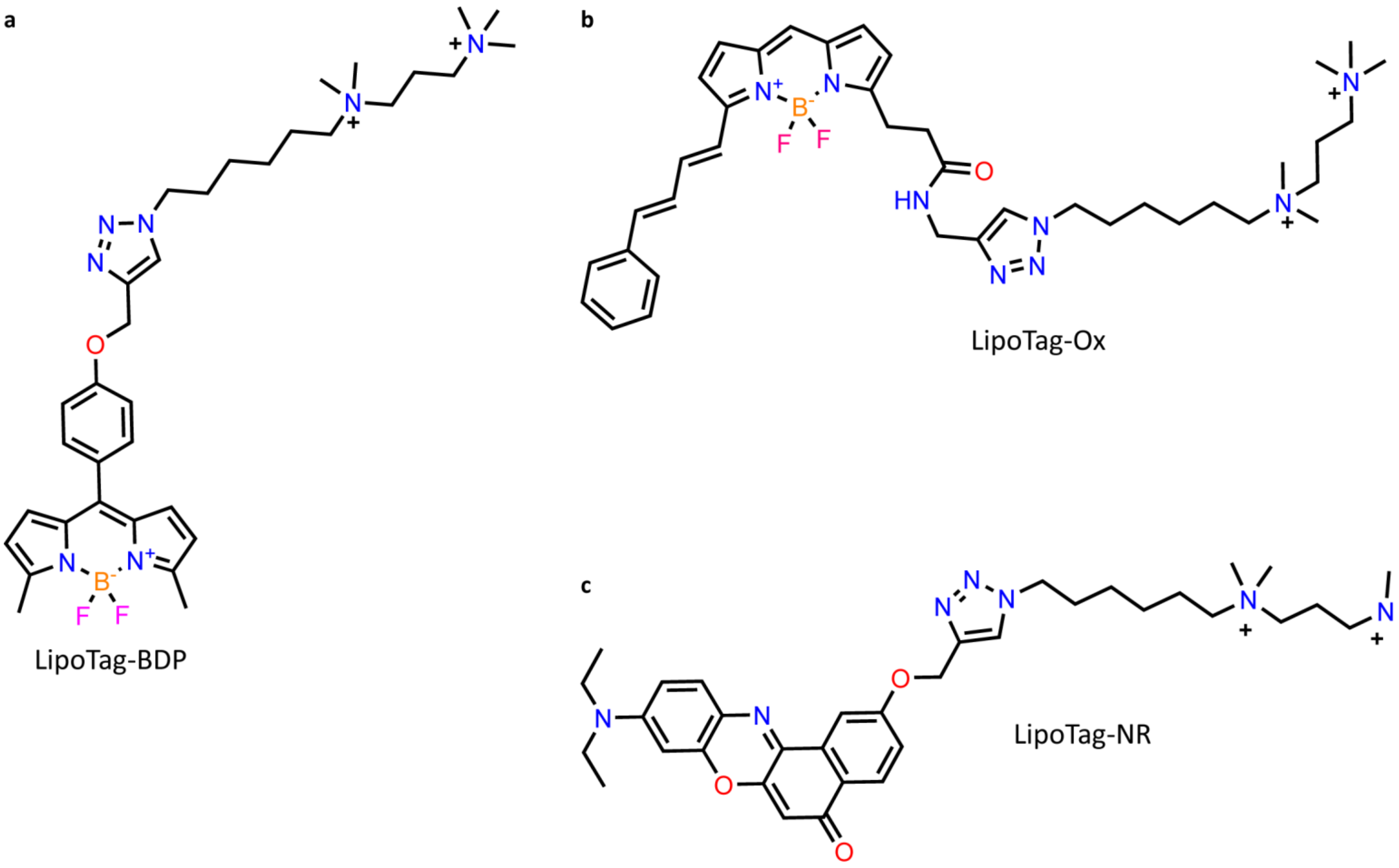


**Fig. S20.** Chemical structures of functional LipoTag probes.

Chemical structure of the membrane tension probe LipoTag-BDP (**a**), the membrane oxidation probe LipoTag-Ox (**b**) and the membrane order probe LipoTag-NR (**c**).

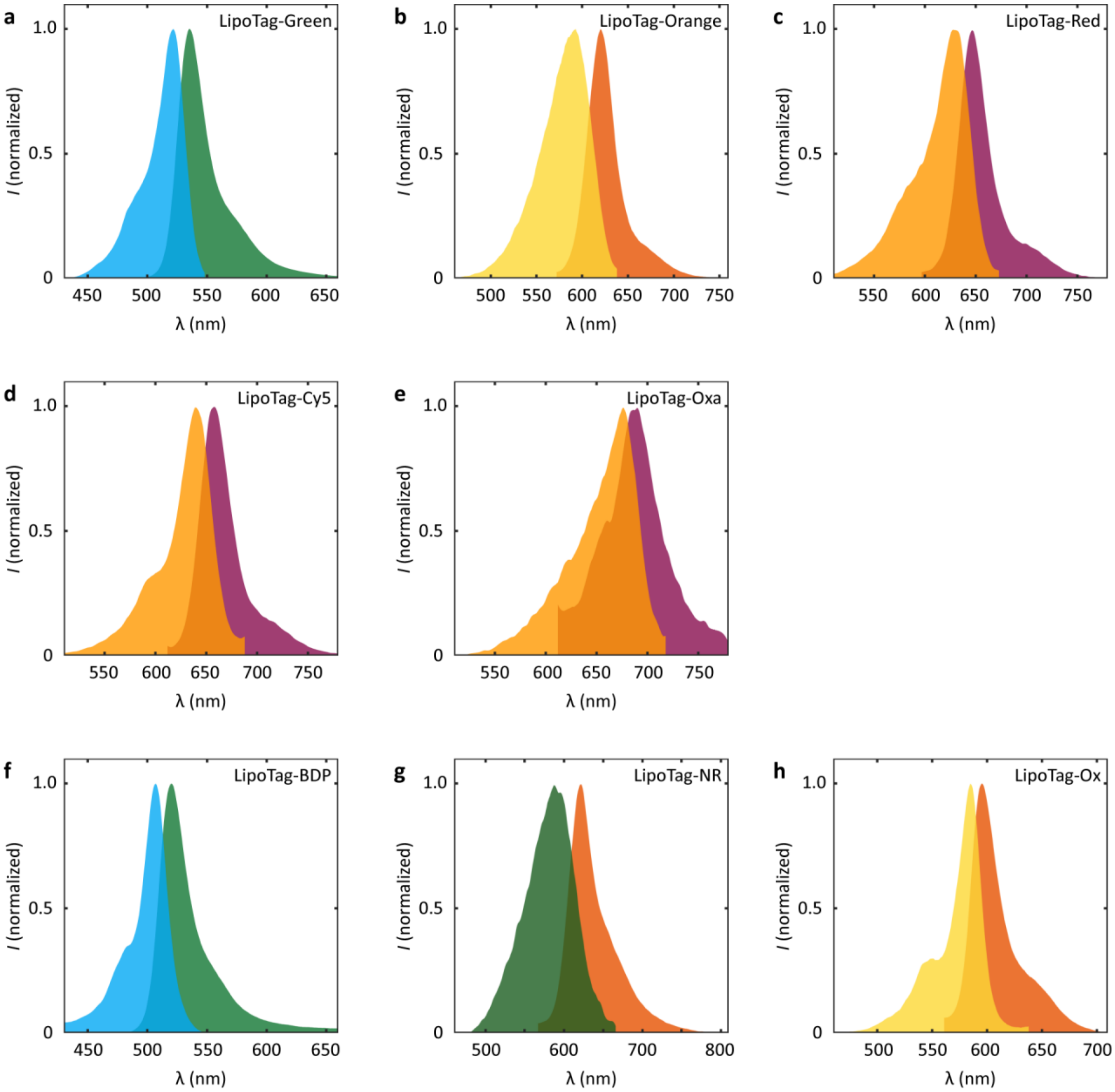


**Fig. S21.** Normalized fluorescence spectra of all LipoTag probes.

Normalized fluorescence excitation and emission spectra for LipoTag-Green (**a**), LipoTag-Orange (**b**), LipoTag-Red (**c**), LipoTag-Cy5 (**d**), LipoTag-Oxa (**e**), LipoTag-BDP (**f**), LipoTag-NR (**g**) and LipoTag-Ox (**h**).

### Number of samples and observations

**Table S1.** Number of samples, multiple cells and regions were often observed per sample

| Experiment | Figures | Sample numbers |
|---|---|---|
| Arabidopsis root imaging (non-functional) | Fig. 1b-d<br>Fig. 2a,b<br>Fig. S1a-t<br>Fig. S10a-f<br>Fig. S18a,b | LipoTag-Green n>20<br>LipoTag-Orange n>20<br>LipoTag-Red n>20<br>LipoTag-Cy5 n=2<br>LipoTag-Oxa n=2<br>FM1-43 n=2<br>DiD n=2<br>DiA n=2<br>BDP-green alkyne n=2<br>BDP-TR n=2<br>BDP630/650 n=2<br>BDP-rotor alkyne n=2 |
| Non-Arabidopsis imaging (non-funtional) | Fig. 2d-o<br>Fig. S11a-f<br>Fig. S12a-k<br>Fig. S13a-g<br>Fig. S14a-f<br>Fig. S15a-f | *C. richardii* n=2<br>*M. polymorpha* n=4<br>*S.rigidula* n=4<br>*Ectocarpus sp* n=5<br>*F. serratus* n=3<br>*S. latissimi* n=3<br>*S. Pombe* n=4<br>*V. dahliae* n=6<br>*E. coli* n>10<br>*M. musculus* n>10 |
| Multicolor imaging | Fig 2p,q | n=6 |
| FRAP | Fig. 1e | LipoTag-Green n=14<br>LipoTag-Orange n=20<br>LipoTag-Red n=19<br>FM4-64 n=16 |
| Toxicity assay | Fig. 1f | n>5 wells with >10 protoplasts per well |
| ABA treatment stomata LipoTag-BDP FLIM | Fig. 3c-e | Open (non-treated) n=19<br>Closed (treated) n=20 |
| Ablation LipoTag-BDP FLIM | Fig. 3f-h | n=9 |
| Sterol depletion MβCD LipoTag-NR ratiometric | Fig. 4c-e | Control n=6<br>Treated n=8 |
| Ablation LipoTag-NR ratiometric | Fig. 4f-h | n=6 |
| Hemin treatment LipoTag-Ox ratiometric | Fig. 5c-e<br>Fig. S19a-d | Treated n=6 per timepoint<br>Mock treatment n=4 |
| Ablation LipoTag-Ox | Fig. 5f-h | Ablation n=9<br>Mock n=3 |
| Tissue penetration | Fig. S3a-t<br>Fig. S5a-j<br>Fig. S6a-j<br>Fig. S7a-j | LipoTag-Green n=3<br>LipoTag-Orange n=3<br>LipoTag-Red n=3<br>FM4-64 n=3 |
| Concentration range | Fig. S2a-p<br>Fig. S4a-r | LipoTag-Green n=3<br>LipoTag-Orange n=3<br>LipoTag-Red n=3<br>FM4-64 n=3 |

| | | |
|---|---|---|
| Plasmolysis | Fig. S9a-i | LipoTag-Green n=4<br>LipoTag-Orange n=6<br>LipoTag-Red n=5<br>FM4-64 n=4 |
| Marchantia gemma plasmodesmata staining | Fig. S16a-c | Aniline blue n = 3<br>Aniline blue + LT-Red n = 3<br>LT-Red = 3 |
| LipoTag-BDP calibration FLIM | Fig. S17 | n=5 per point |

### NMR spectra

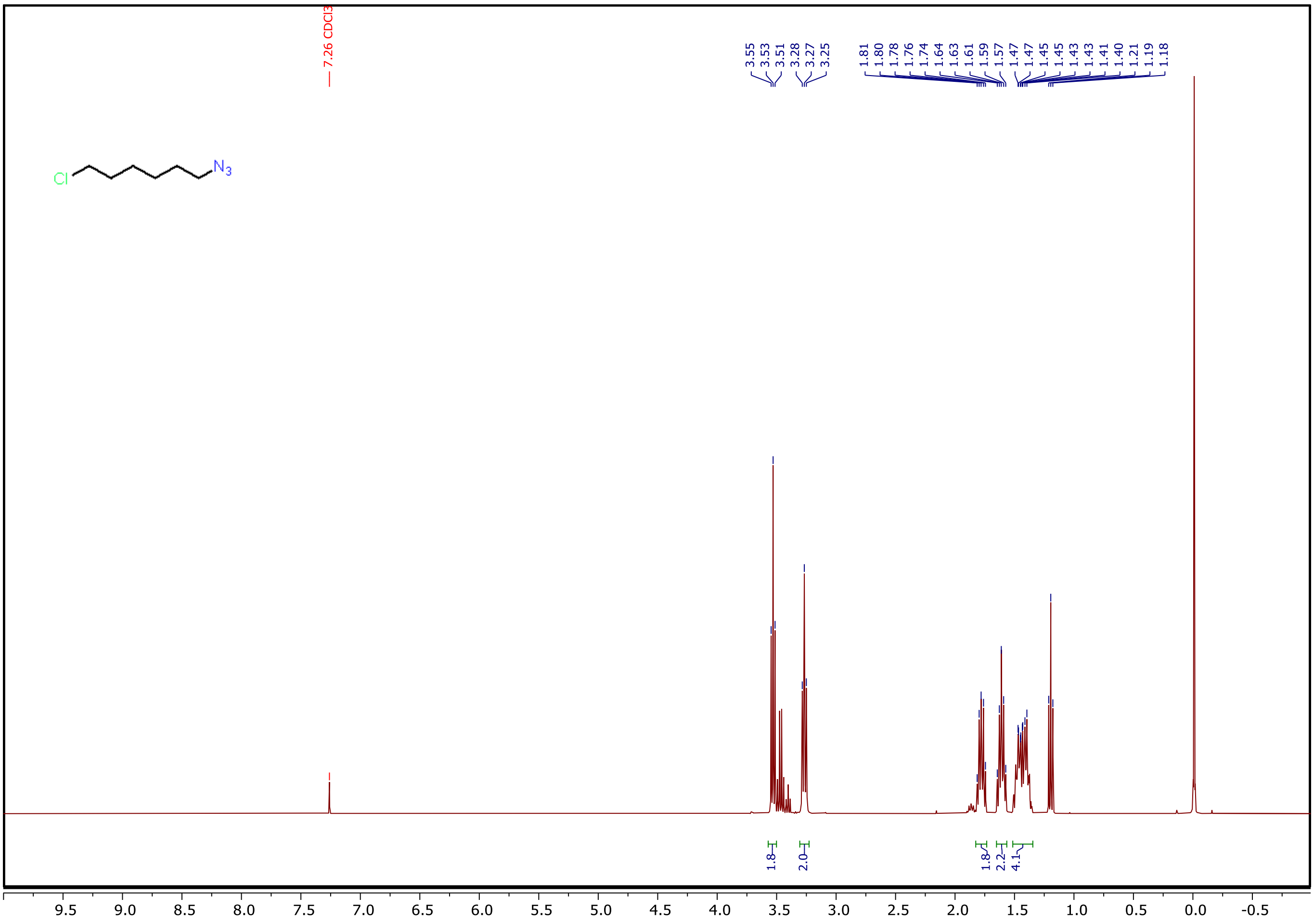

**Fig. S22.** $^{1}H$ spectrum of **1**.

$^{1}H$ NMR (400 MHz, $CDCl_3$) δ 3.53 (t, *J* = 6.6 Hz, 2H), 3.27 (t, *J* = 6.9 Hz, 2H), 1.83 – 1.73 (m, 2H), 1.65 – 1.56 (m, 2H), 1.51 – 1.35 (m, 4H).

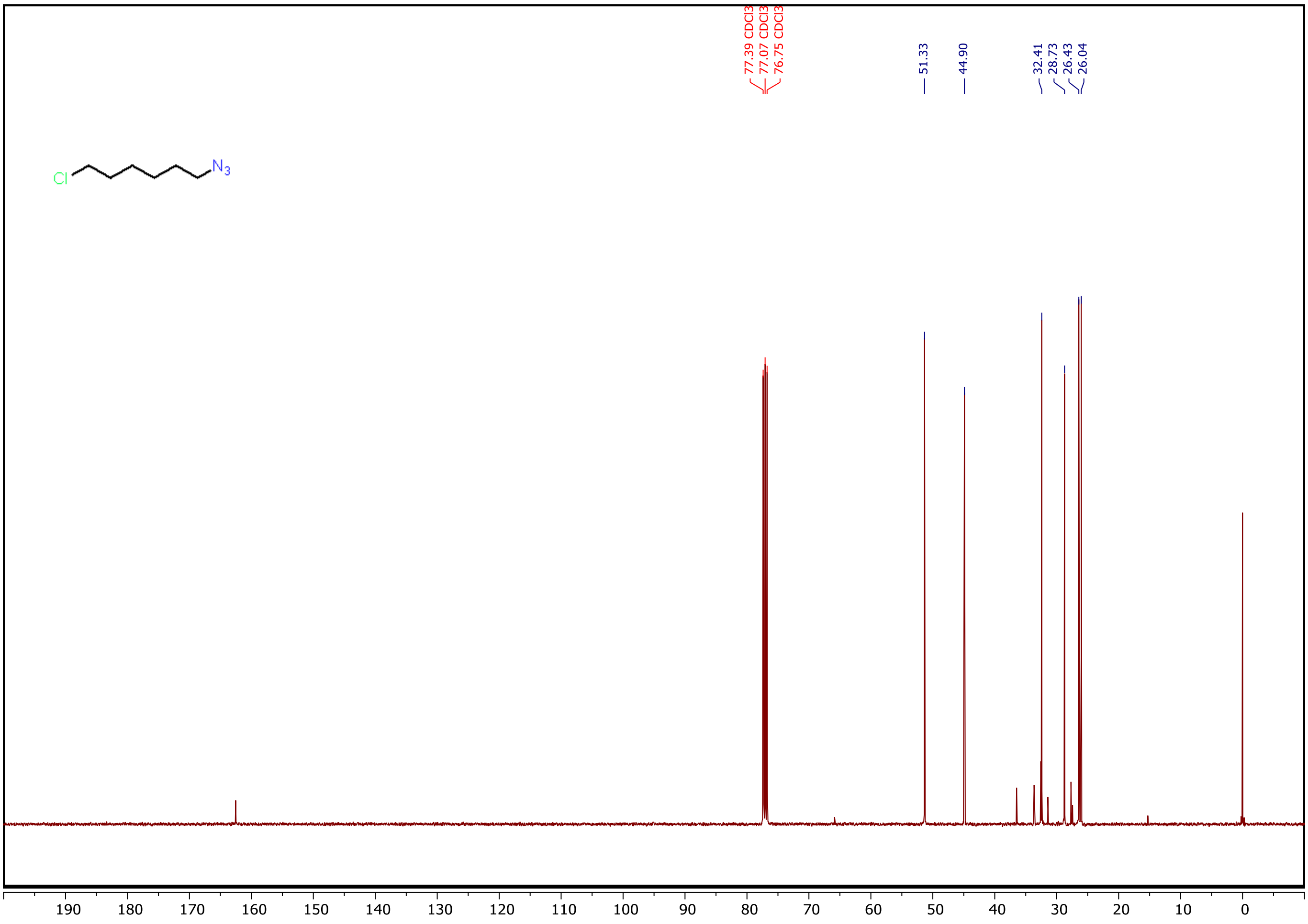


**Fig. S23.** $^{13}$C spectrum of **1**.

$^{13}$C NMR (101 MHz, CDCl3) δ 51.33, 44.90, 32.41, 28.73, 26.43, 26.04.

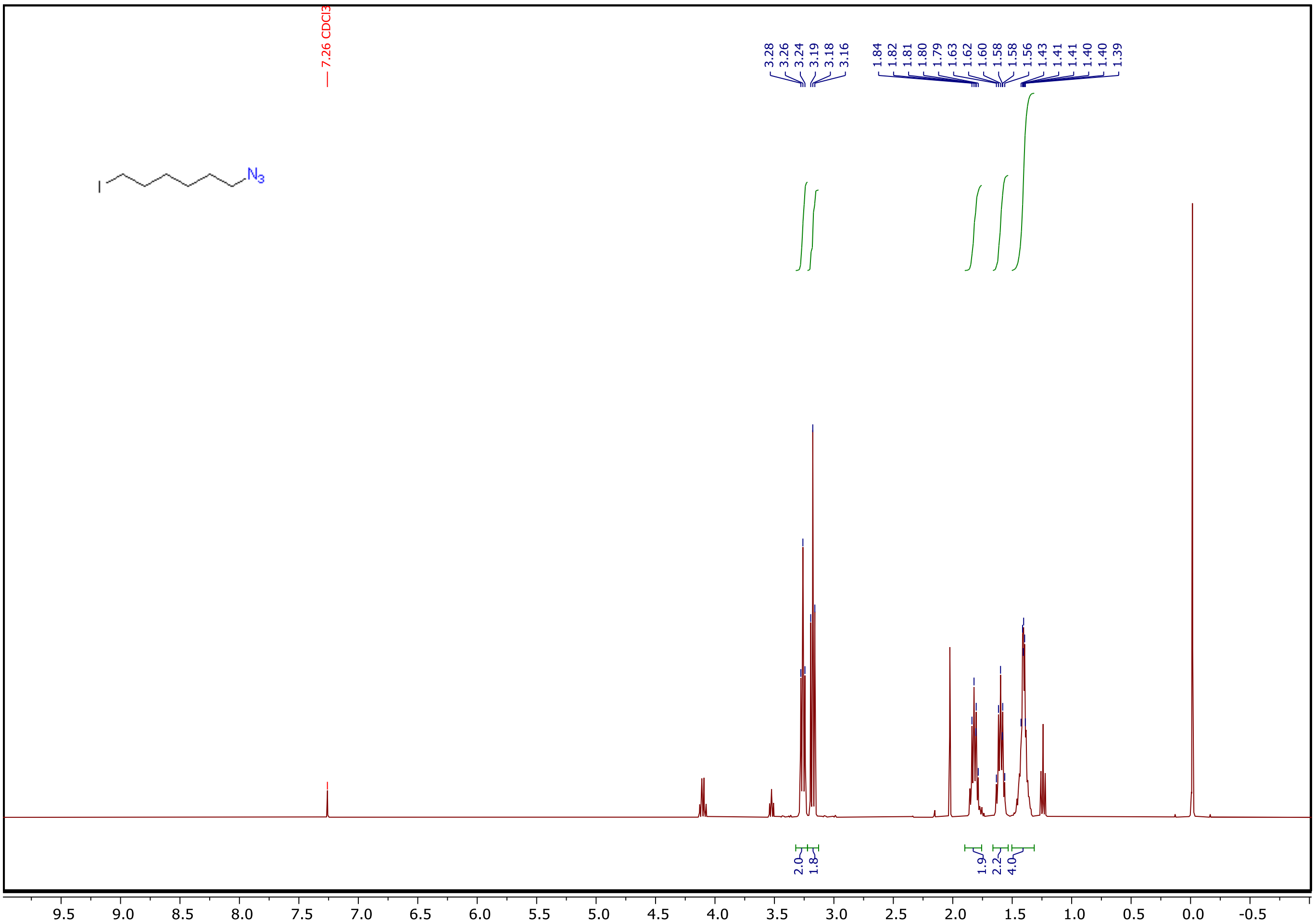

**Fig. S24.** $^1$H spectrum of **2**.

$^1$H NMR (400 MHz, $CDCl_3$) δ 3.26 (t, *J* = 6.9 Hz, 2H), 3.18 (t, *J* = 6.9 Hz, 2H), 1.88 – 1.72 (m, 2H), 1.66 – 1.53 (m, 2H), 1.51 – 1.32 (m, 4H).

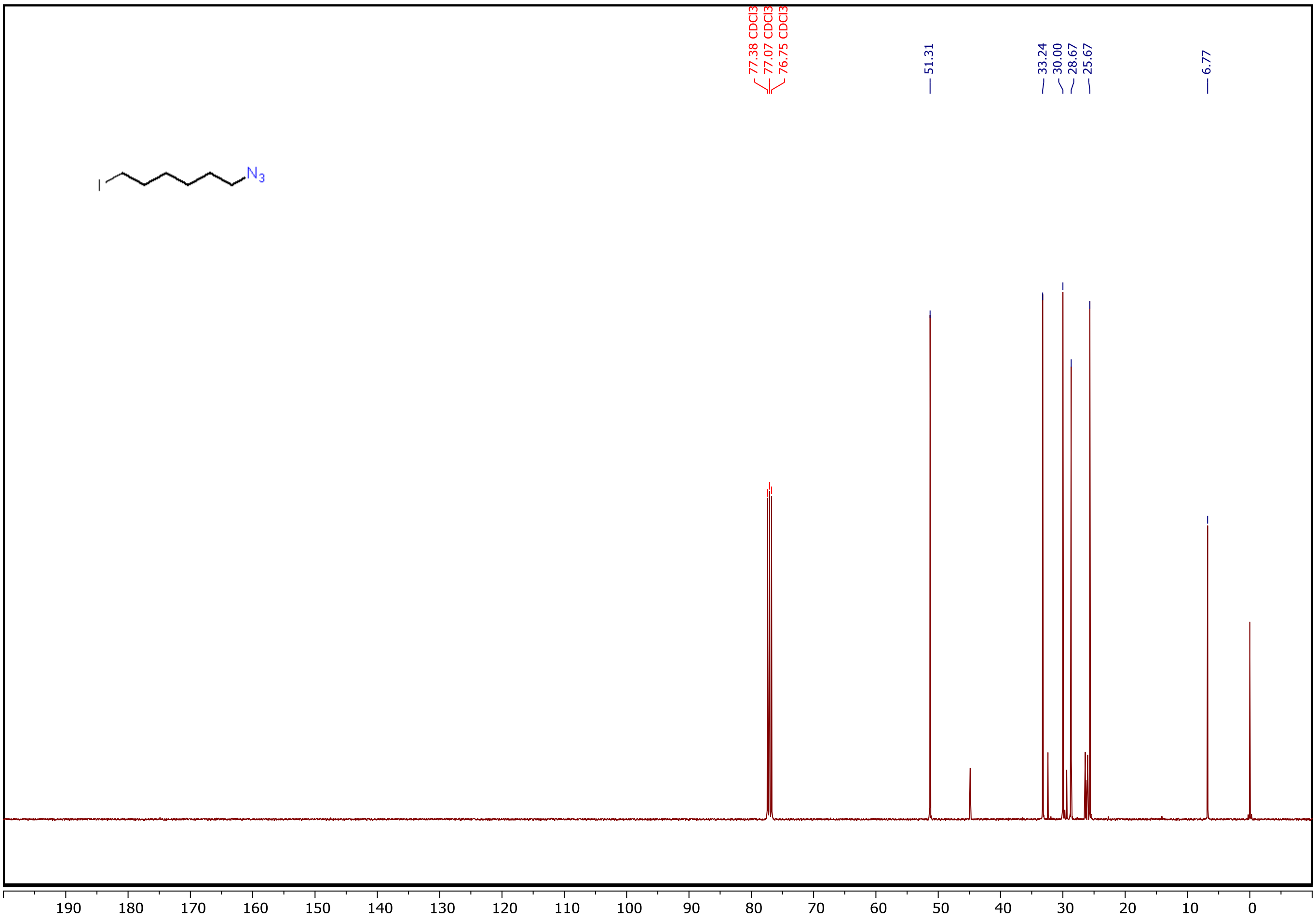


**Fig. S25.** $^{13}C$ spectrum of **2**.

$^{13}C$ NMR (101 MHz, $CDCl_3$) δ 51.31, 33.24, 30.00, 28.67, 25.67, 6.77.

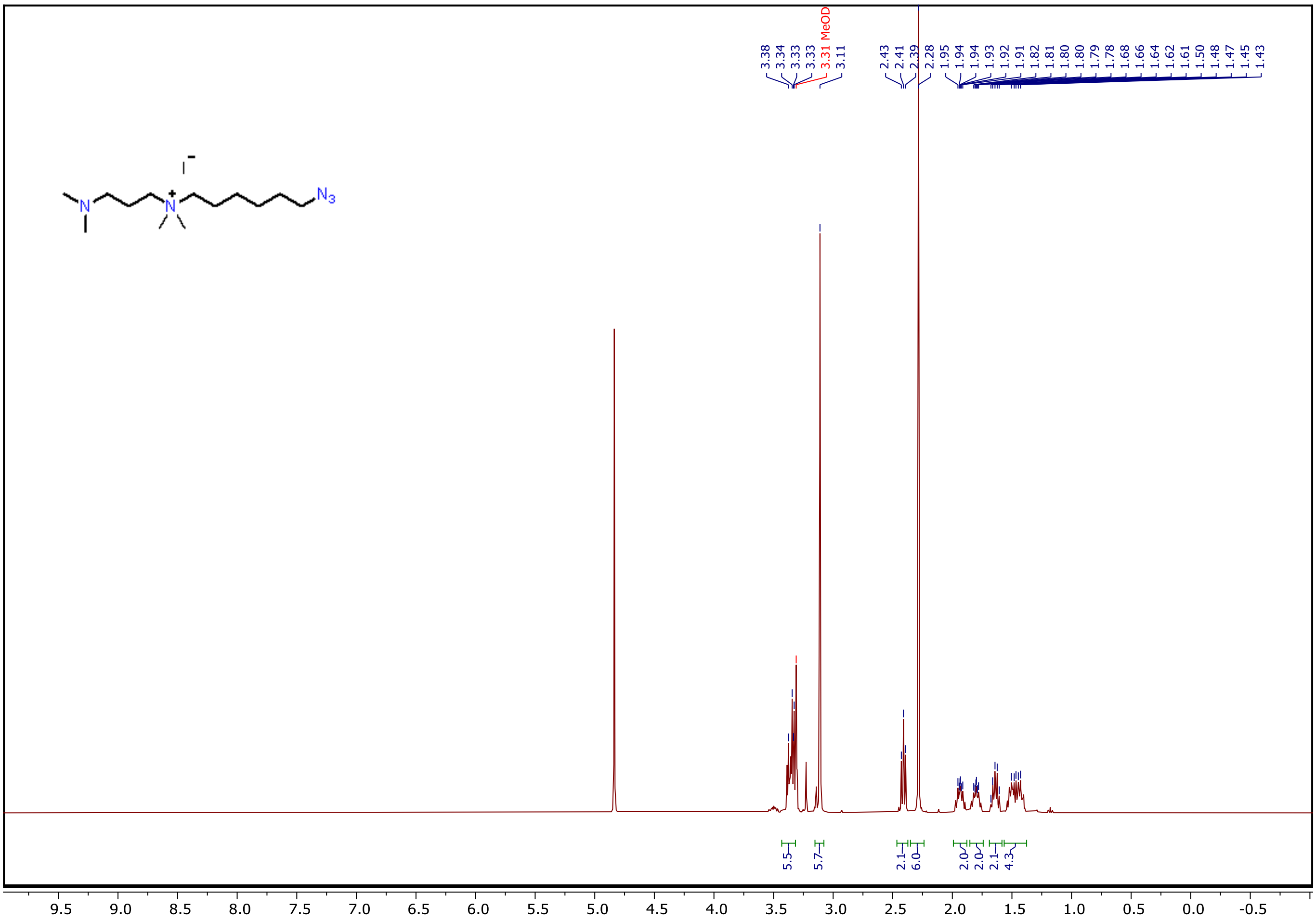

**Fig. S26.** $^{1}$H spectrum of **3**.

$^{1}$H NMR (400 MHz, MeOD) δ 3.43 – 3.32 (m, 6H), 3.11 (s, 6H), 2.41 (t, J = 7.1 Hz, 2H), 2.28 (s, 6H), 1.99 – 1.88 (m, 2H), 1.85 – 1.74 (m, 2H), 1.69 – 1.58 (m, 2H), 1.57 – 1.38 (m, 4H).

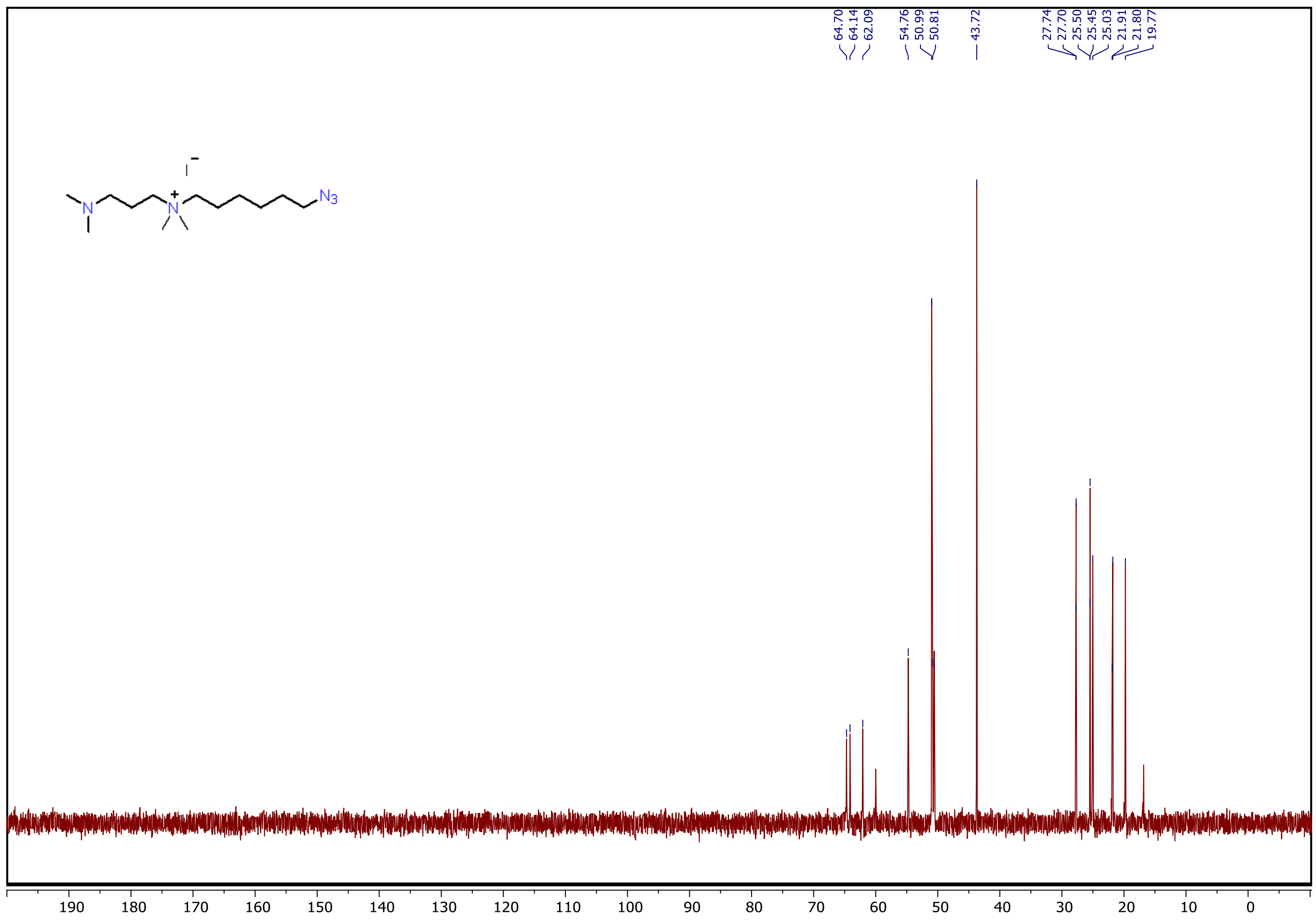


**Fig. S27.** $^{13}C$ spectrum of **3**.

$^{13}C$ NMR (101 MHz, D2O) δ 64.70, 64.14, 62.09, 54.76, 50.99, 50.81, 43.72, 27.74, 27.70, 25.50, 25.45, 25.03, 21.91, 21.80, 19.77.

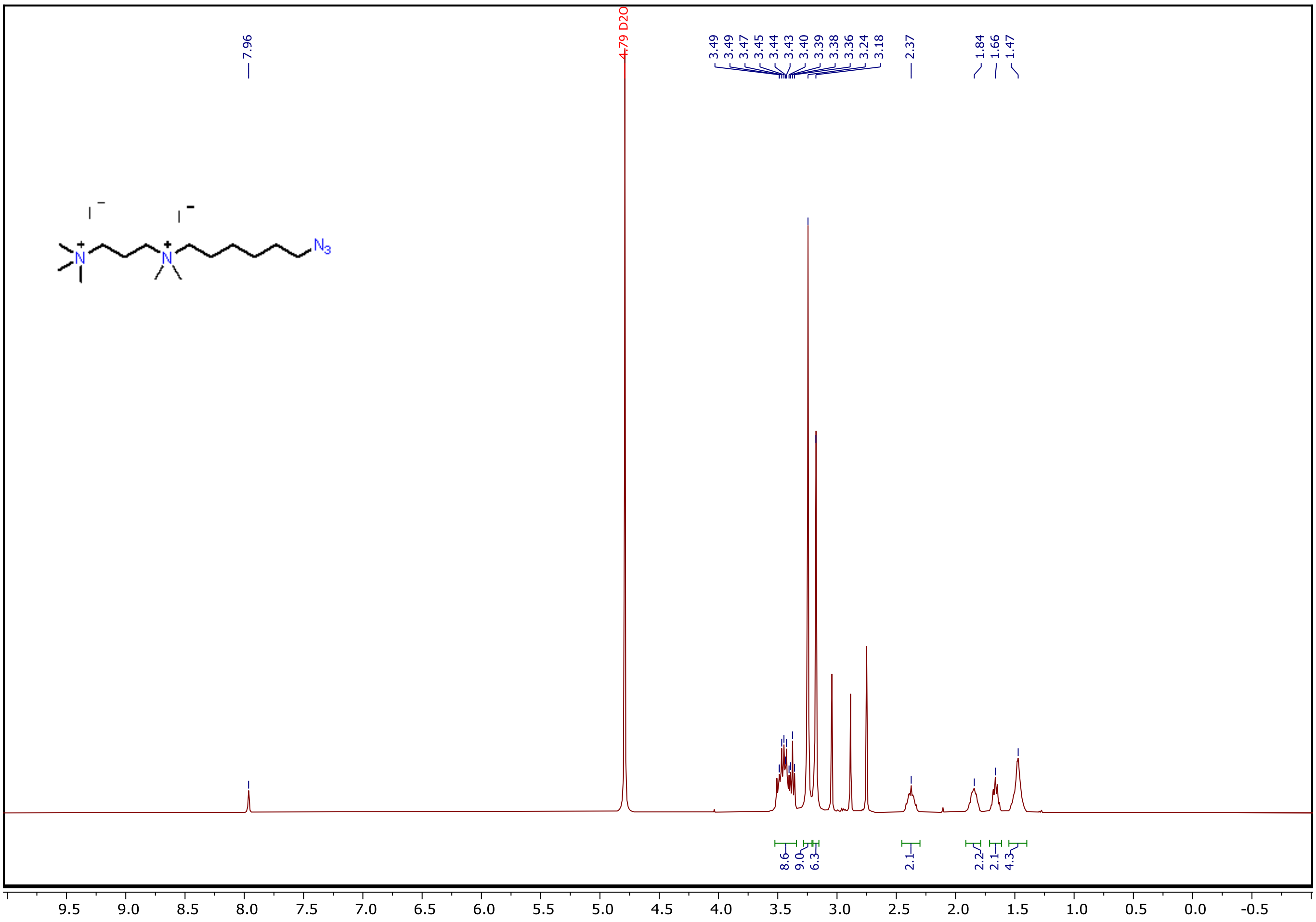


**Fig. S28.** $^1$H spectrum of **4**.

$^1$H NMR (400 MHz, $D_2O$) δ 3.52 – 3.34 (m, 9H), 3.24 (s, 9H), 3.18 (s, 6H), 2.37 (s, 2H), 1.84 (s, 2H), 1.66 (s, 2H), 1.47 (s, 4H).

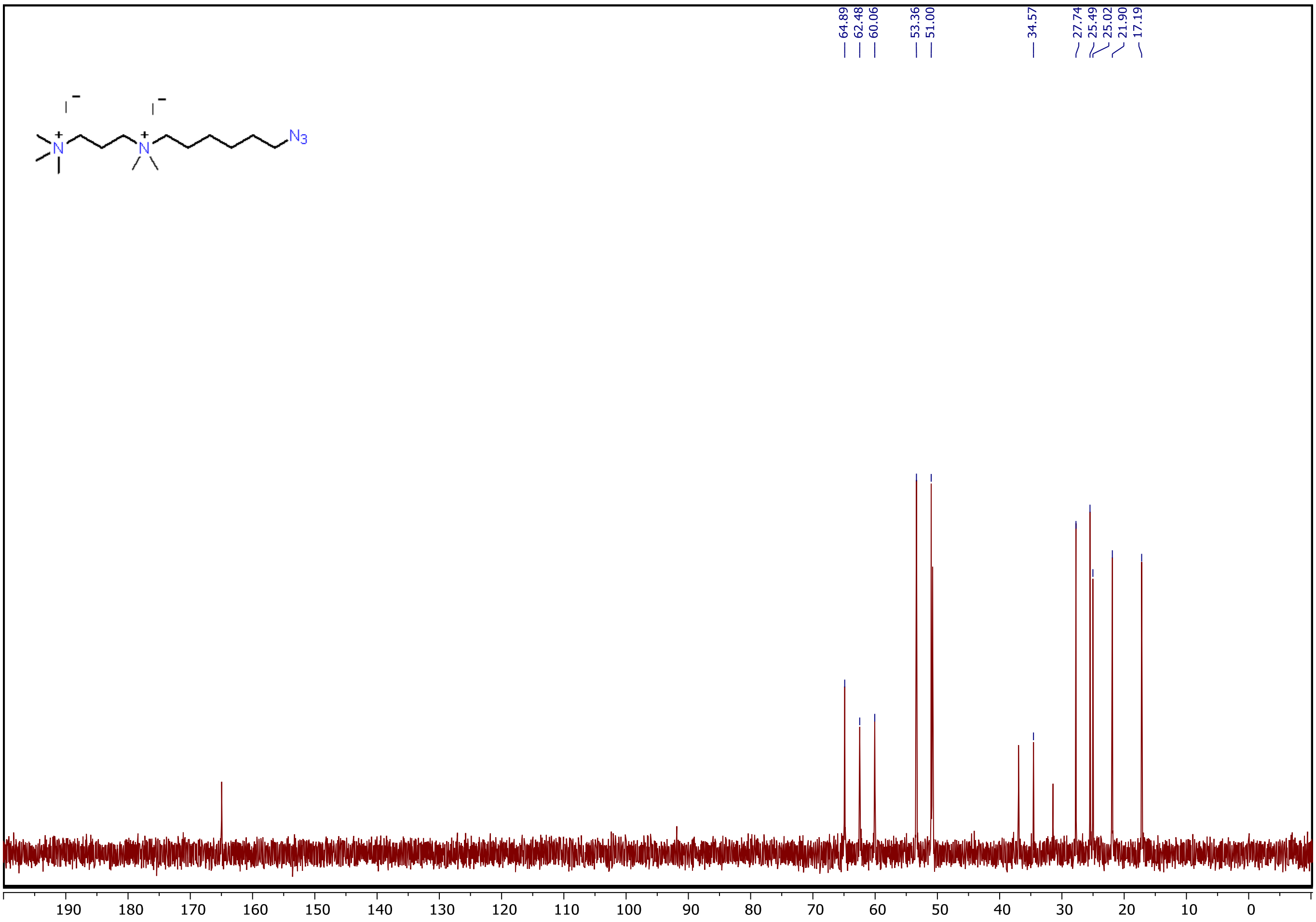

**Fig. S29.** $^{13}C$ spectrum of **4**.

$^{13}C$ NMR (101 MHz, $D_2O$) δ 64.89, 62.48, 60.06, 53.36, 51.00, 34.57, 27.74, 25.49, 25.02, 21.90, 17.19.

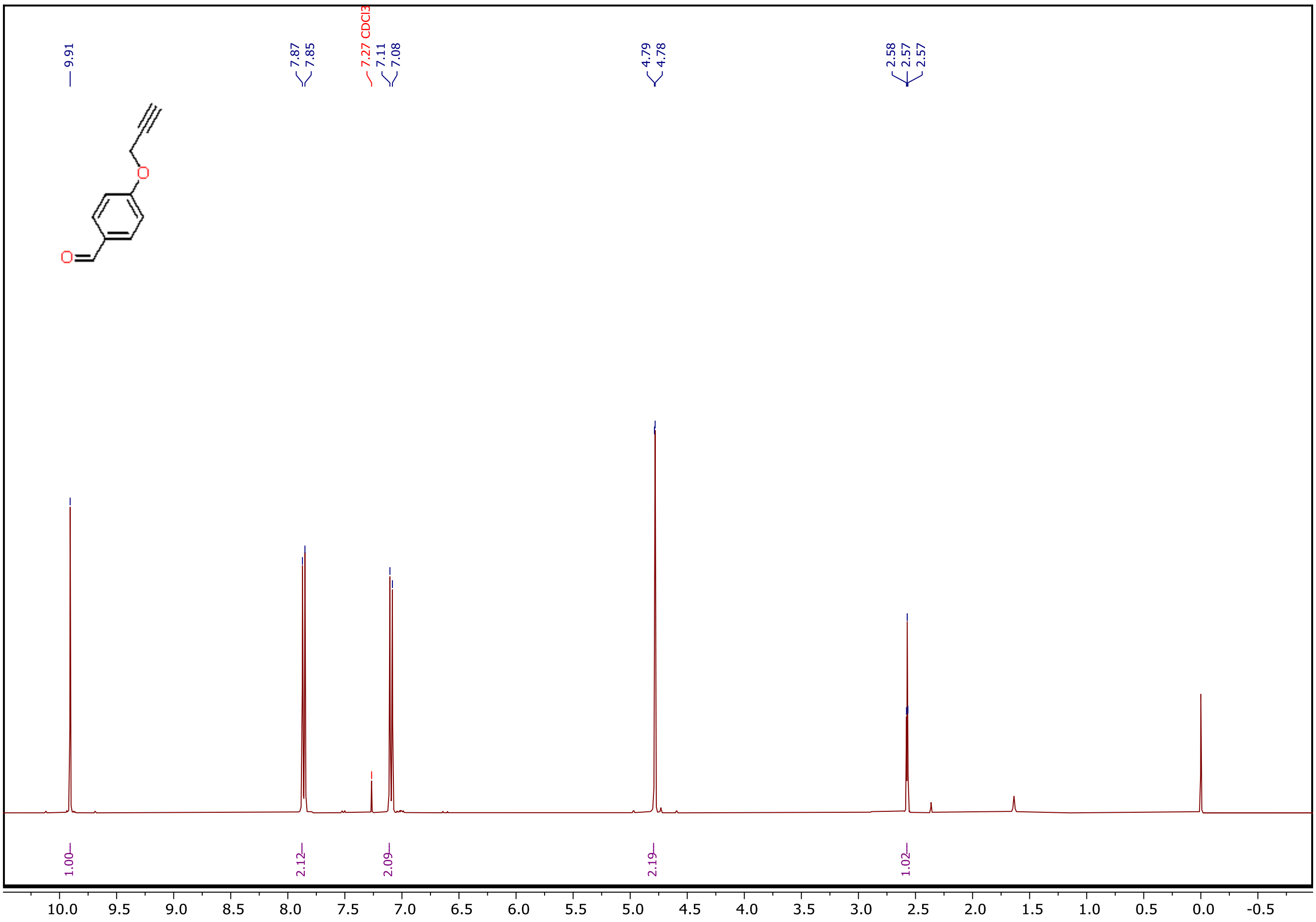


**Fig. S30.** $^{1}H$ spectrum of **5**.

$^{1}H$ NMR (400 MHz, CDCl3) δ 9.91 (s, 1H), 7.86 (d, J = 8.8 Hz, 2H), 7.09 (d, J = 8.8 Hz, 2H), 4.78 (d, J = 2.4 Hz, 2H), 2.57 (t, J = 2.4 Hz, 1H).

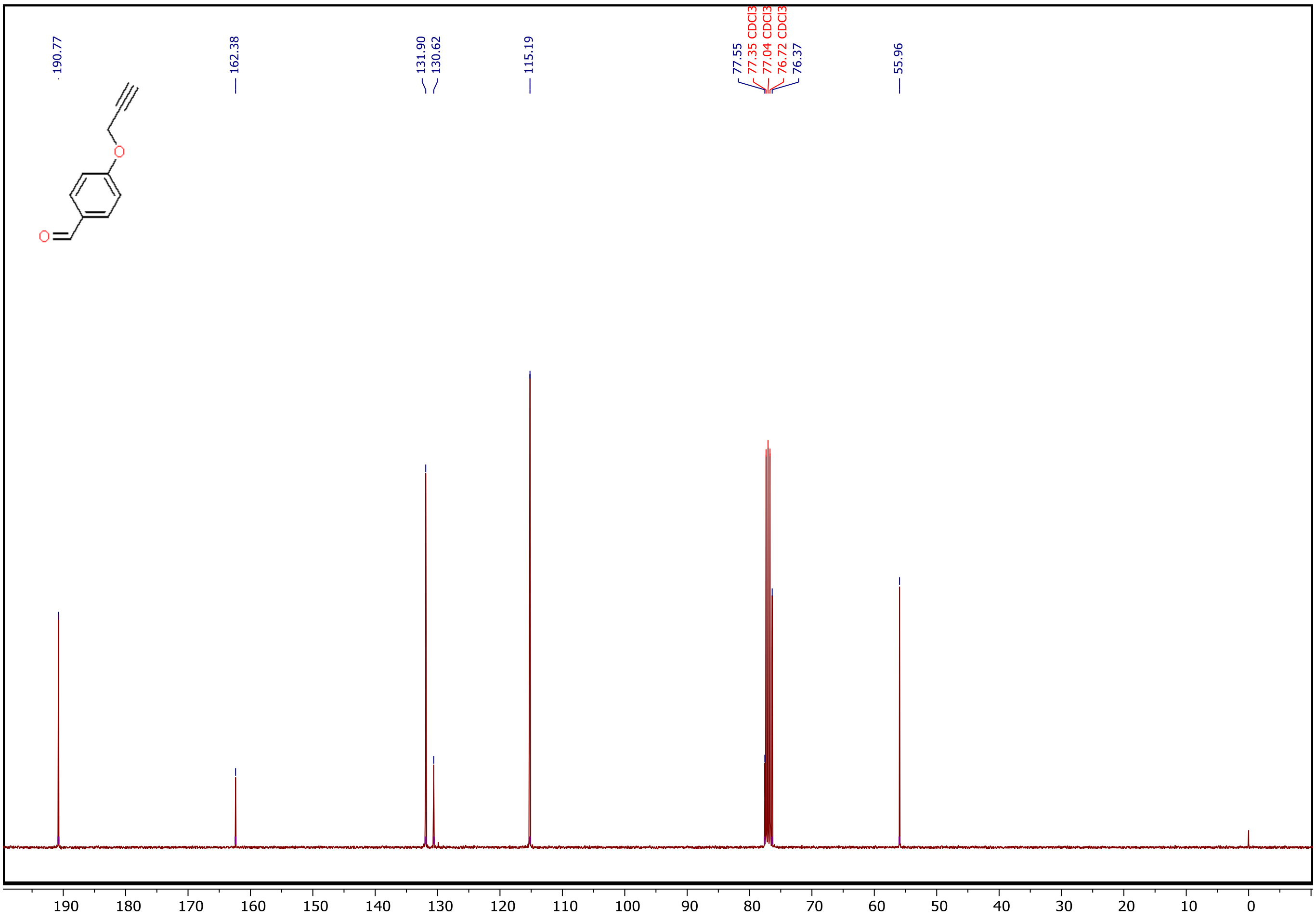


**Fig. S31.** $^{13}C$ spectrum of **5**.

$^{13}C$ NMR (101 MHz, $CDCl_3$) δ 190.77, 162.38, 131.90, 130.62, 115.19, 77.55, 76.37, 55.96.

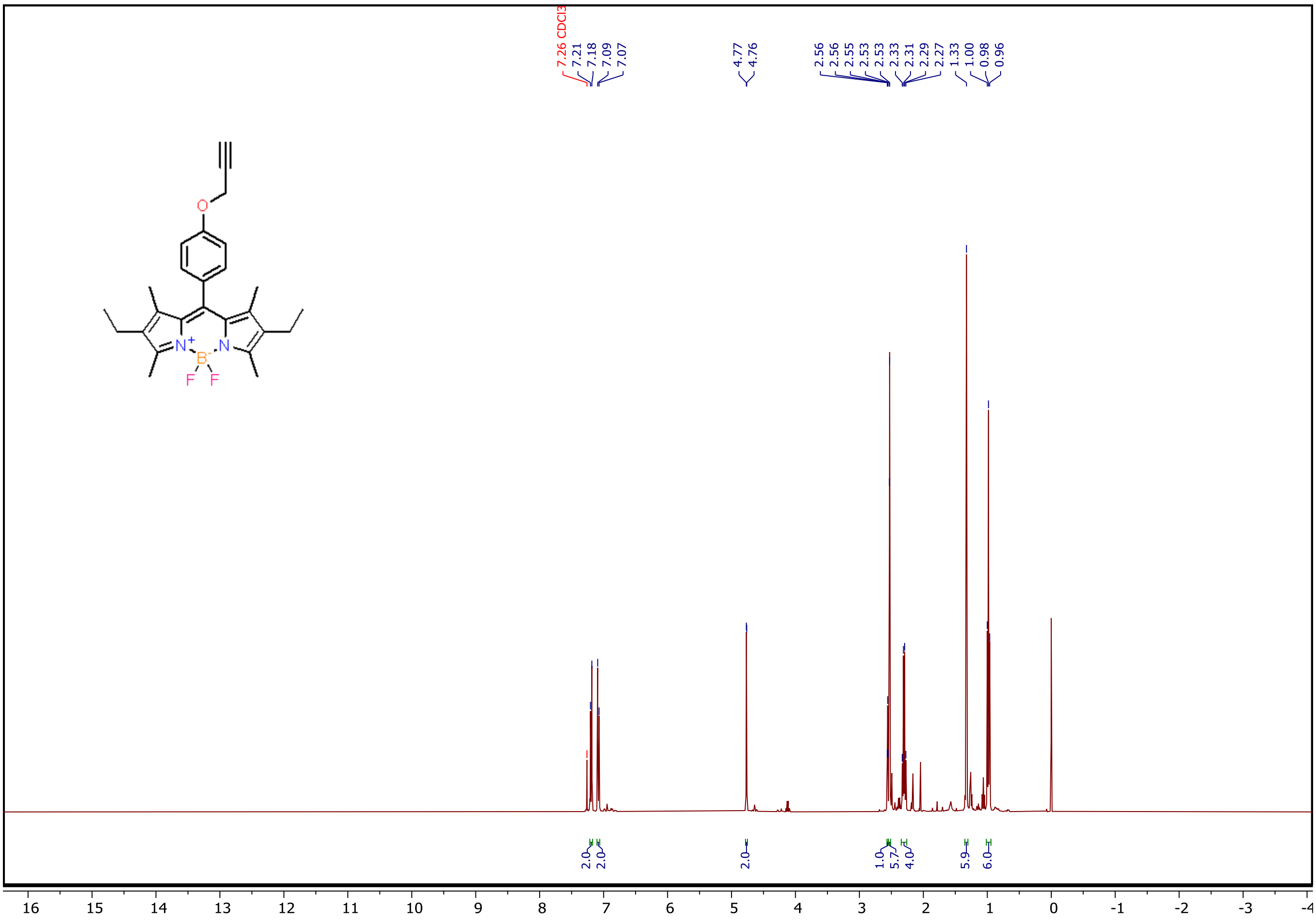

**Fig. S32.** $^{1}H$ spectrum of **6**.

$^{1}H$ NMR (400 MHz, CDCl3) δ 7.19 (d, J = 8.6 Hz, 2H), 7.08 (d, J = 8.7 Hz, 2H), 4.77 (d, J = 2.4 Hz, 2H), 2.56 (t, J = 2.4 Hz, 1H), 2.53 (d, J = 1.3 Hz, 6H), 2.30 (q, J = 7.5 Hz, 4H), 1.33 (s, 6H), 0.98 (t, J = 7.5 Hz, 6H).

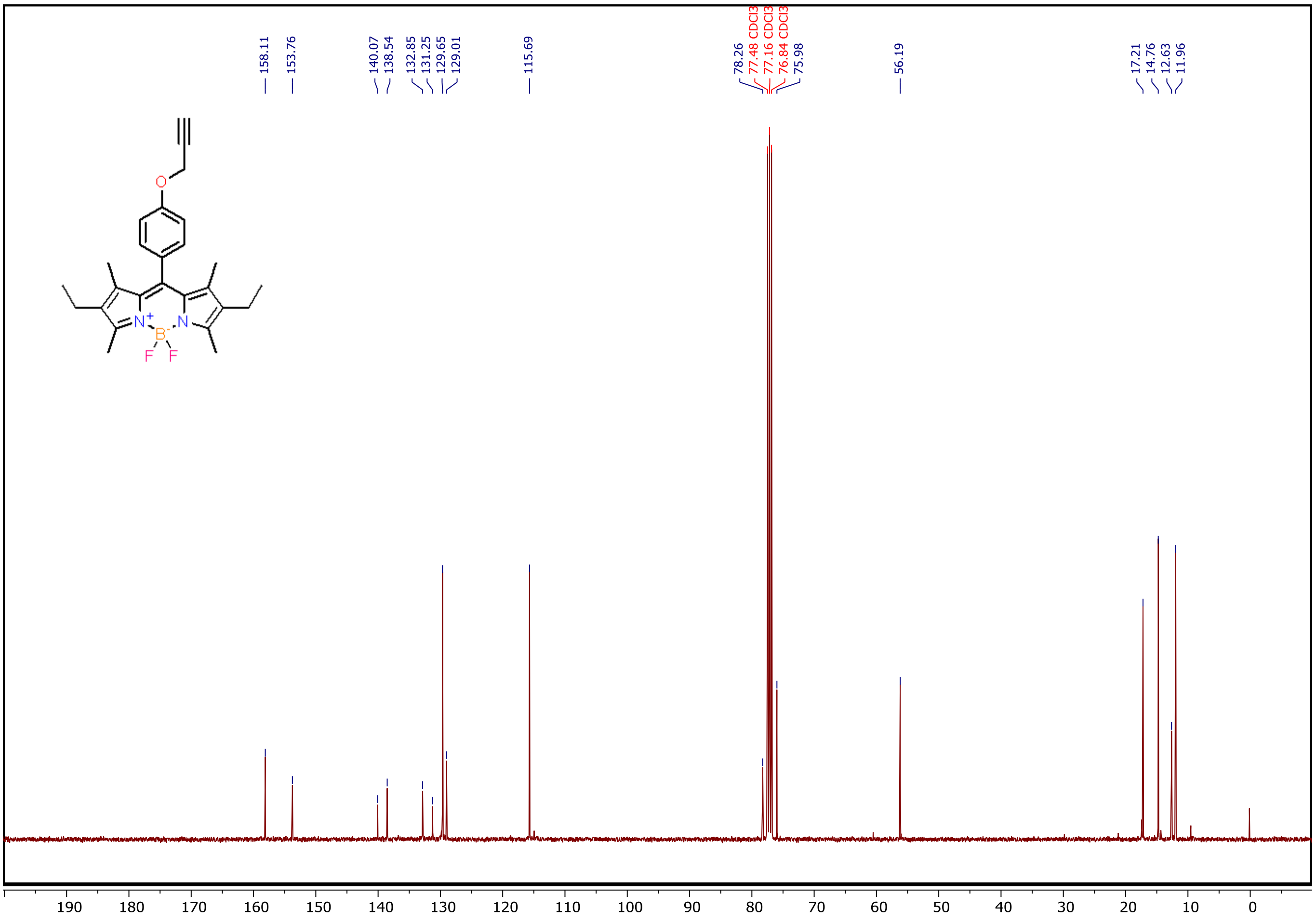


**Fig. S33.** $^{13}$C spectrum of **6**.

$^{13}$C NMR (101 MHz, CDCl3) δ 158.11, 153.76, 140.07, 138.54, 132.85, 131.25, 129.65, 129.01, 115.69, 78.26, 75.98, 56.19, 17.21, 14.76, 12.63, 11.96.

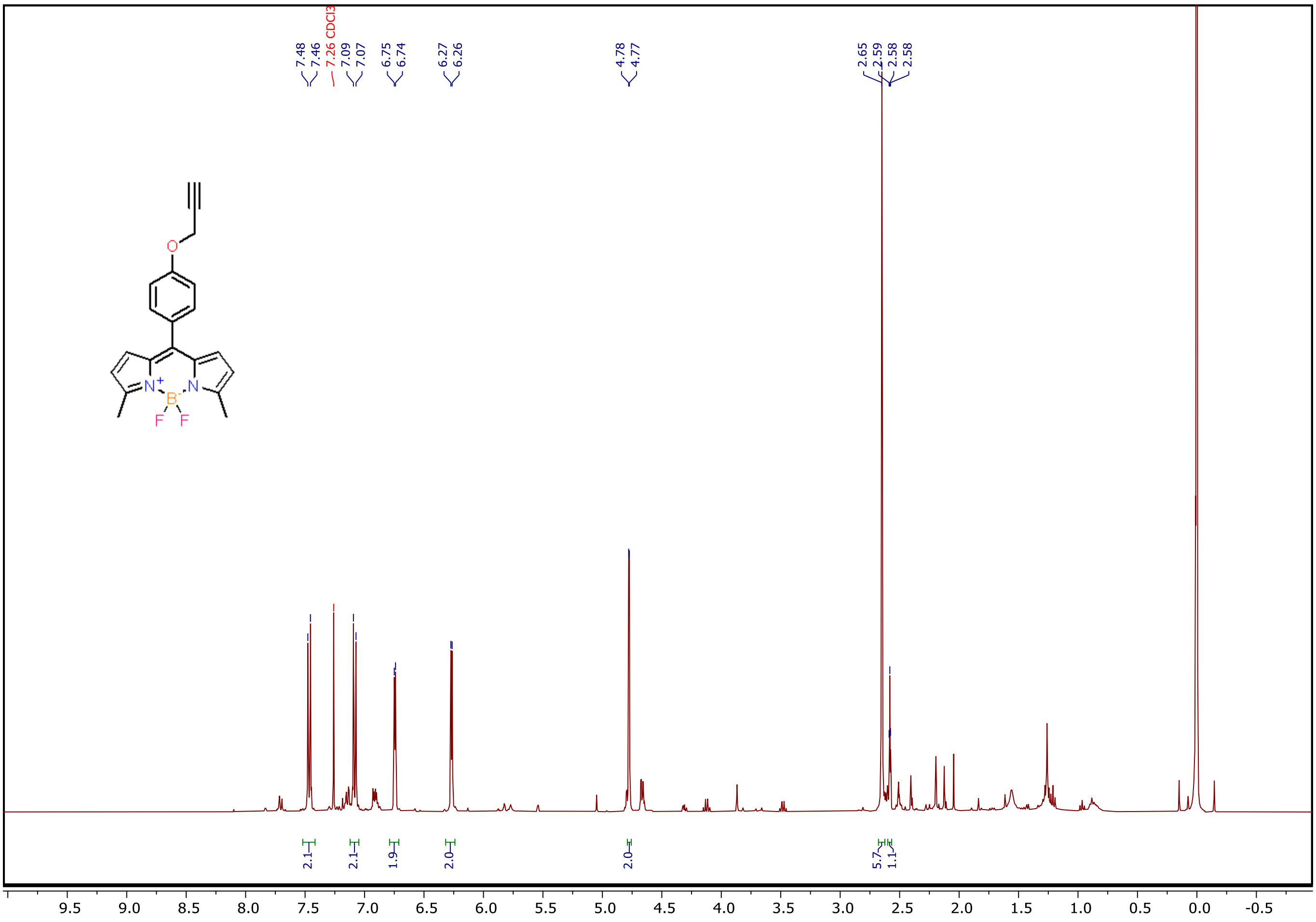

**Fig. S34.** $^{1}H$ spectrum of **7**.

$^{1}H$ NMR (400 MHz, $CDCl_3$) δ 7.47 (d, *J* = 8.7 Hz, 2H), 7.08 (d, *J* = 8.7 Hz, 2H), 6.74 (d, *J* = 4.1 Hz, 2H), 6.27 (d, *J* = 4.2 Hz, 2H), 4.78 (d, *J* = 2.4 Hz, 2H), 2.65 (s, 6H), 2.58 (t, *J* = 2.4 Hz, 1H).

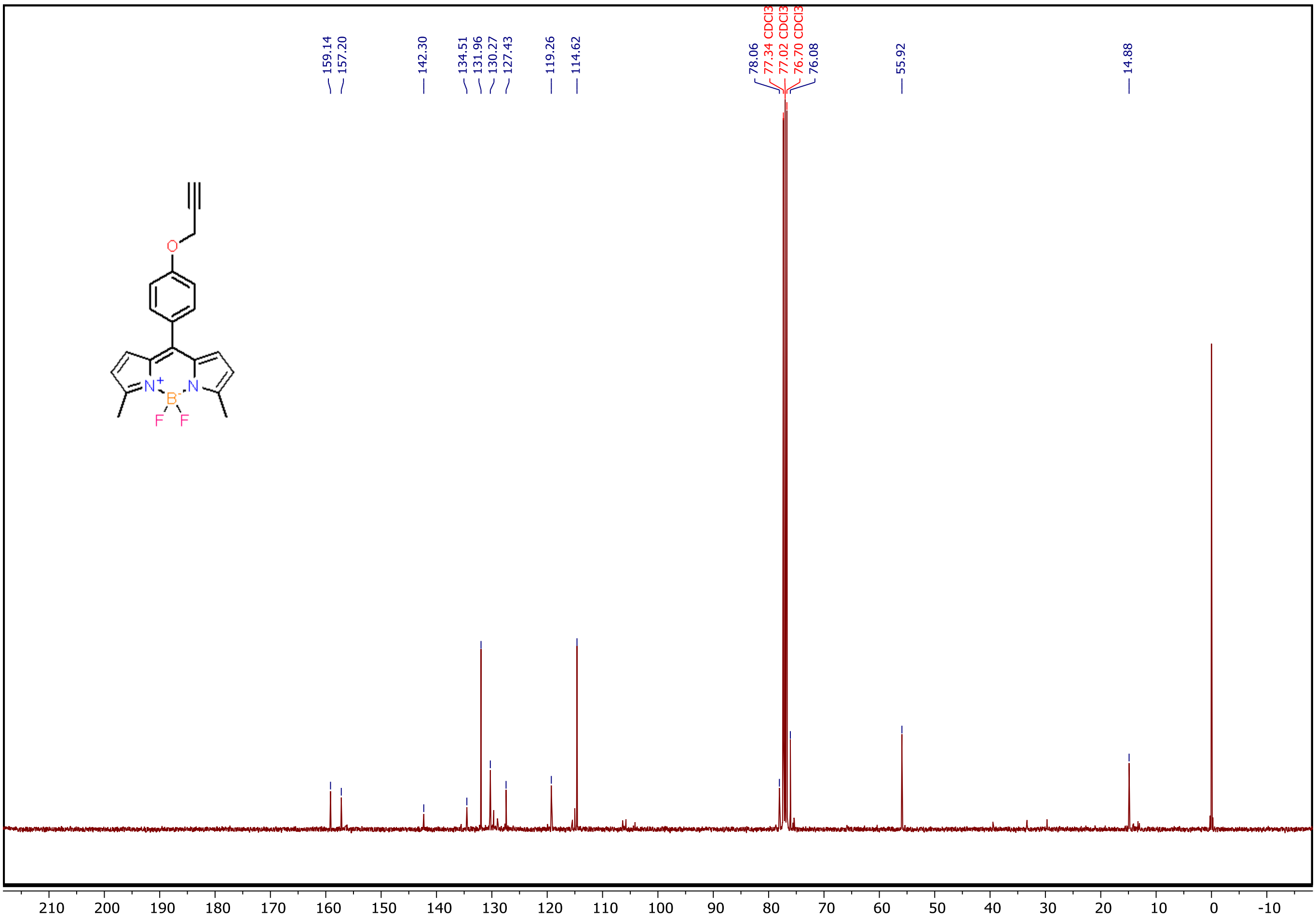


**Fig. S35.** $^{13}C$ spectrum of **7**.

$^{13}C$ NMR (101 MHz, $CDCl_3$) δ 159.14, 157.20, 142.30, 134.51, 131.96, 130.27, 127.43, 119.26, 114.62, 78.06, 76.08, 55.92, 14.88.

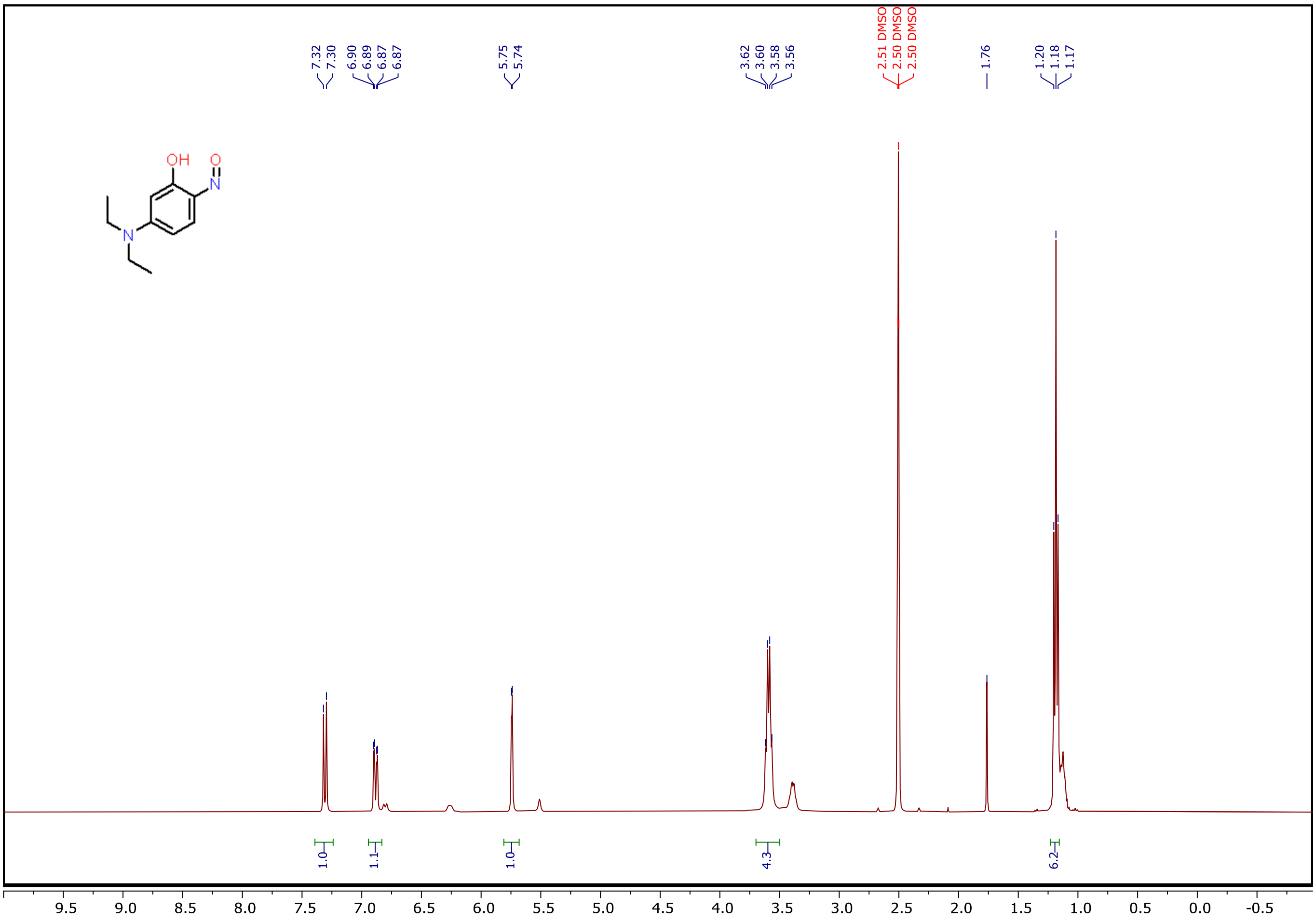


**Fig. S36.** $^{1}$H spectrum of **8**.

$^{1}$H NMR (400 MHz, DMSO) δ 7.31 (d, J = 9.9 Hz, 1H), 6.88 (dd, J = 10.0, 2.6 Hz, 1H), 5.74 (d, J = 2.6 Hz, 1H), 3.59 (q, J = 7.1 Hz, 4H), 1.18 (t, J = 7.1 Hz, 6H).

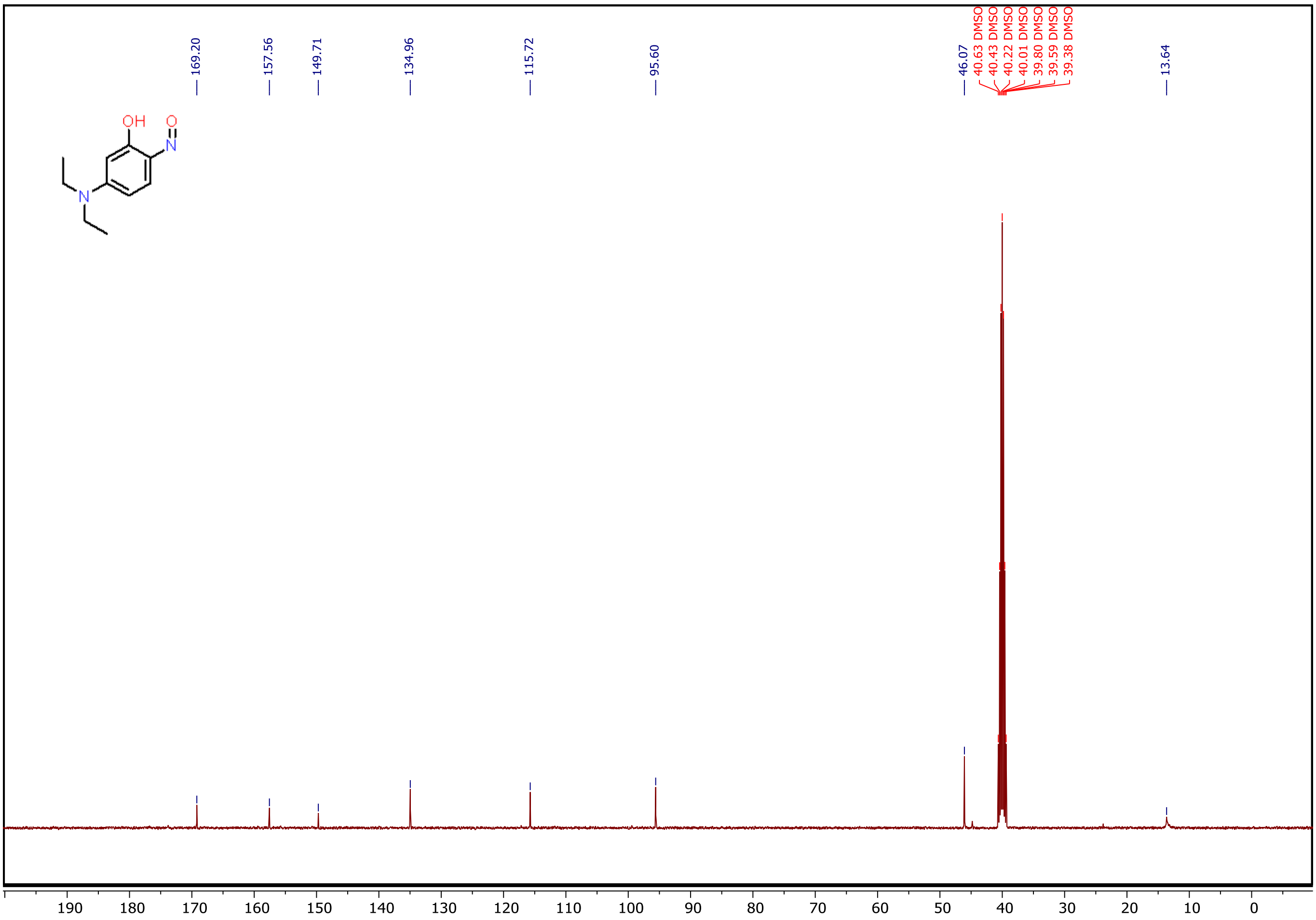


**Fig. S37.** $^{13}C$ spectrum of **8**.

$^{13}C$ NMR (101 MHz, DMSO) δ 169.20, 157.56, 149.71, 134.96, 115.72, 95.60, 46.07, 13.64.

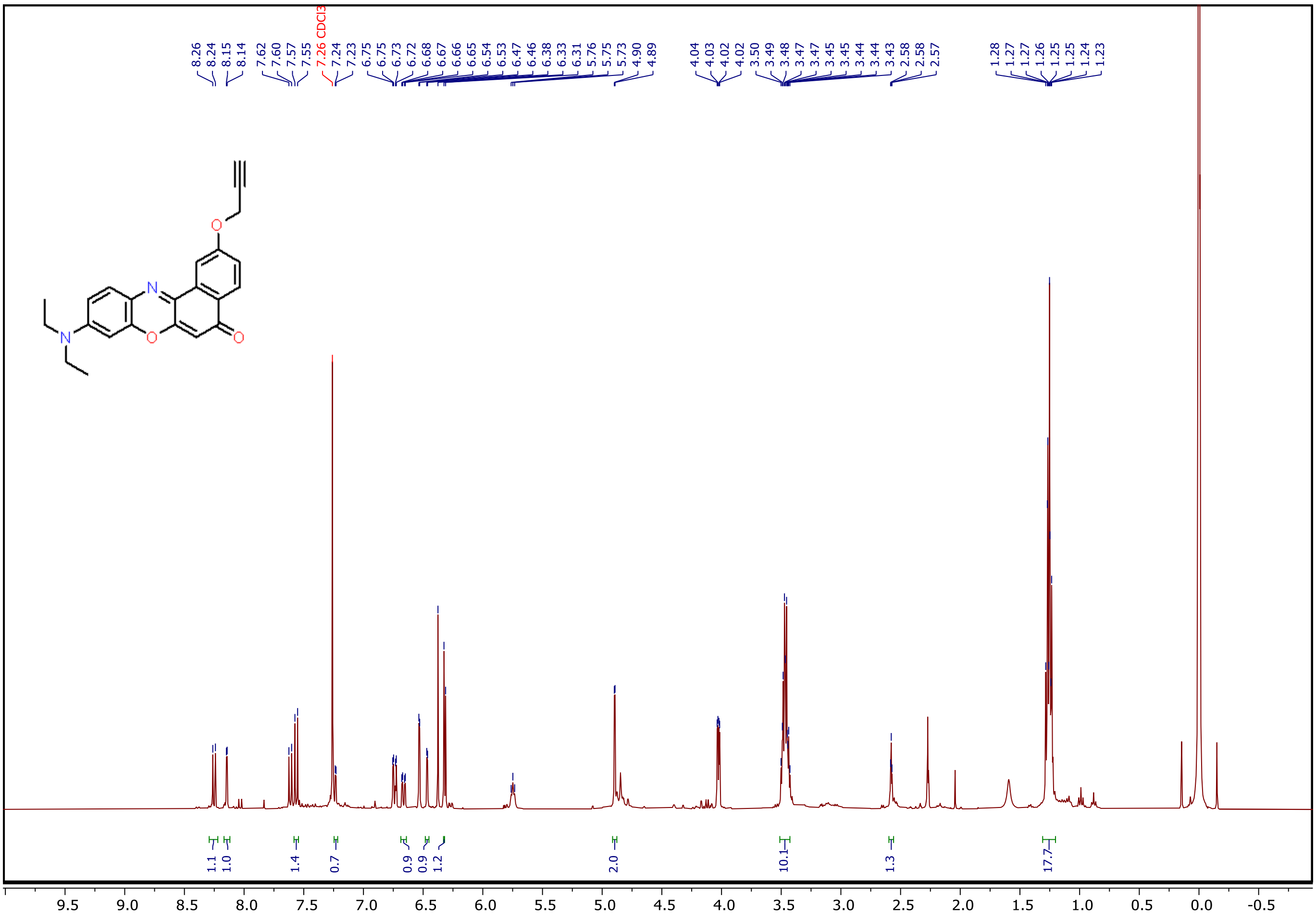

**Fig. S38.** $^{1}$H spectrum of **9**.

$^{1}$H NMR (400 MHz, $CDCl_3$) δ 8.25 (d, *J* = 8.8 Hz, 1H), 8.14 (d, *J* = 2.6 Hz, 1H), 7.56 (d, *J* = 9.1 Hz, 1H), 7.23 (d, *J* = 2.6 Hz, 1H), 6.66 (dd, *J* = 9.0, 2.7 Hz, 1H), 6.47 (d, *J* = 2.7 Hz, 1H), 6.33 (s, 1H), 4.90 (d, *J* = 2.4 Hz, 2H), 3.47 (qd, *J* = 7.1, 4.3 Hz, 10H), 2.58 (t, *J* = 2.4 Hz, 1H), 1.26 (td, *J* = 7.1, 5.7 Hz, 18H)

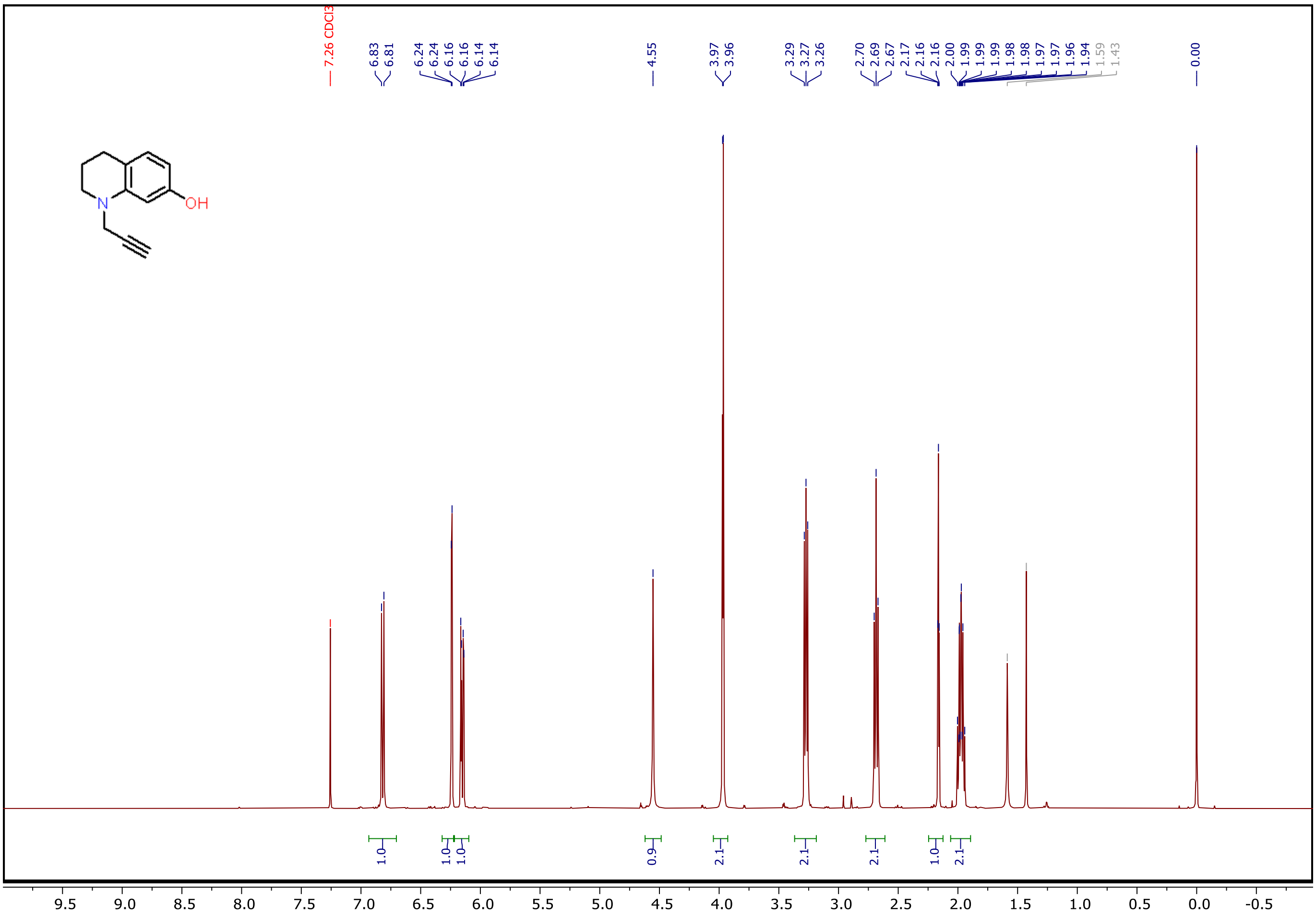

**Fig. S41.** $^{1}H$ spectrum of **10**.

$^{1}H$ NMR (400 MHz, CDCl3) δ 6.82 (d, J = 8.0 Hz, 1H), 6.24 (d, J = 2.4 Hz, 1H), 6.15 (dd, J = 8.0, 2.4 Hz, 1H), 4.55 (s, 1H), 3.97 (d, J = 2.4 Hz, 2H), 3.34 – 3.18 (m, 2H), 2.69 (t, J = 6.5 Hz, 2H), 2.16 (t, J = 2.4 Hz, 1H), 2.02 – 1.93 (m, 2H).

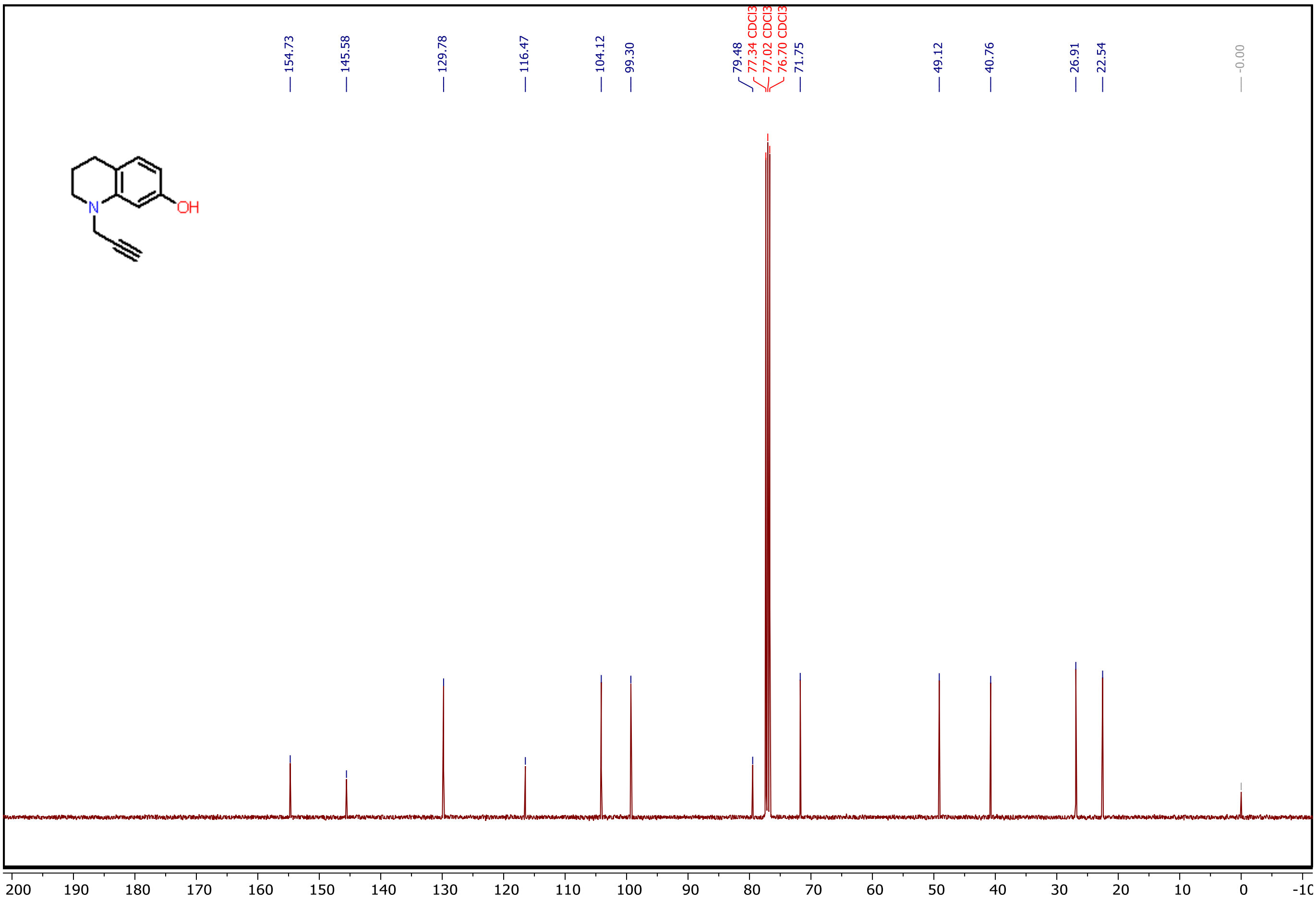


**Fig. S42.** $^{13}C$ spectrum of **10**.

$^{13}C$ NMR (101 MHz, CDCl3) δ 154.73, 145.58, 129.78, 116.47, 104.12, 99.30, 79.48, 71.75, 49.12, 40.76, 26.91, 22.54.

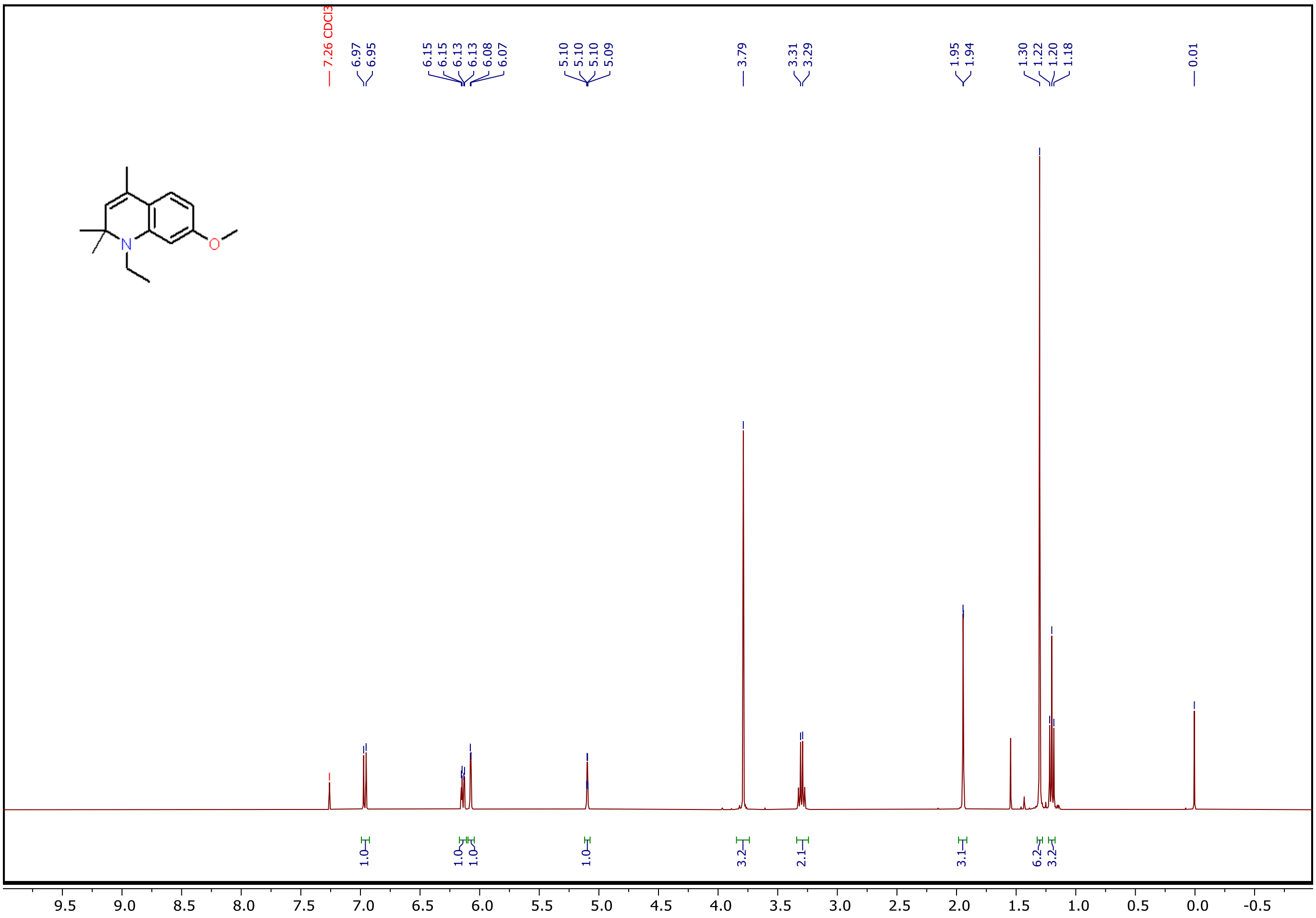


**Fig. S39.** $^1$H spectrum of **11**.

$^1$H NMR (400 MHz, $CDCl_3$) δ 6.96 (d, *J* = 8.3 Hz, 1H), 6.14 (dd, *J* = 8.3, 2.4 Hz, 1H), 6.08 (d, *J* = 2.4 Hz, 1H), 5.10 (q, *J* = 1.4 Hz, 1H), 3.79 (s, 3H), 3.30 (d, *J* = 7.1 Hz, 2H), 1.94 (d, *J* = 1.4 Hz, 3H), 1.30 (s, 6H), 1.20 (t, *J* = 7.0 Hz, 3H).

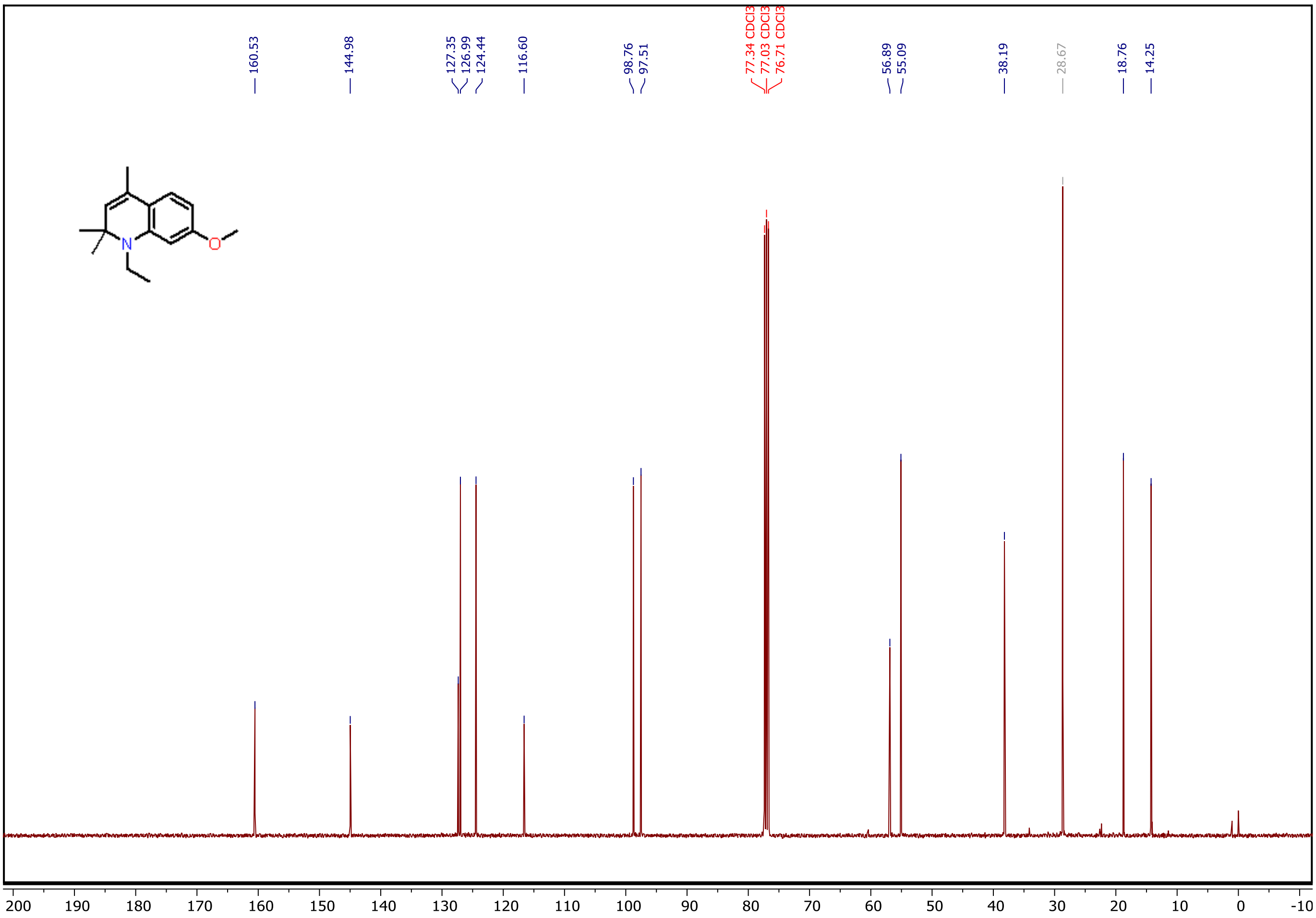


**Fig. S40.** $^{13}$C spectrum of **11**.

$^{13}$C NMR (101 MHz, CDCl3) δ 160.53, 144.98, 127.35, 126.99, 124.44, 116.60, 98.76, 97.51, 56.89, 55.09, 38.19, 18.76, 14.25.

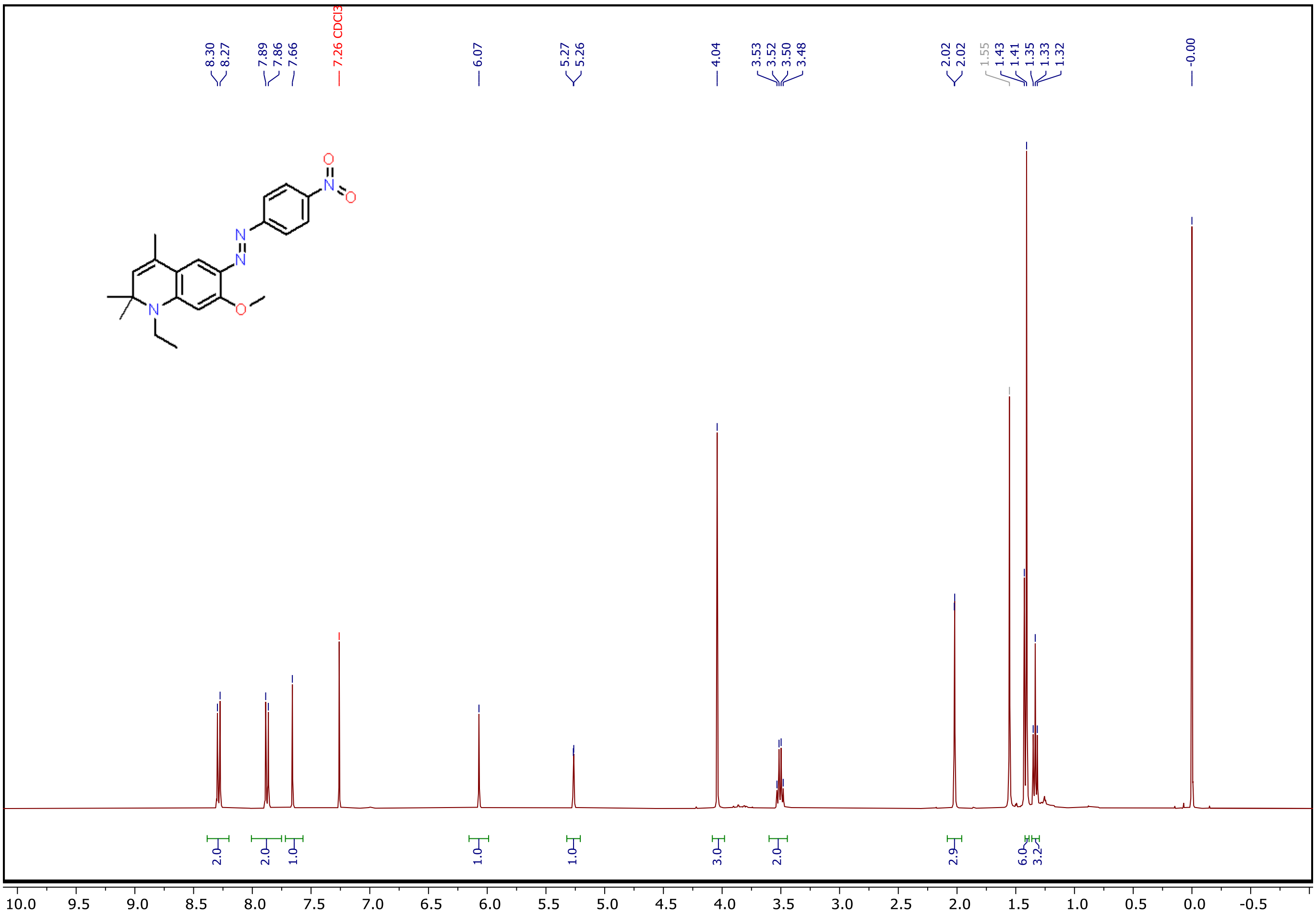

**Fig. S43.** $^{1}$H spectrum of **12**.

$^{1}$H NMR (400 MHz, CDCl3) δ 8.29 (d, J = 9.0 Hz, 2H), 7.87 (d, J = 9.0 Hz, 2H), 7.66 (s, 1H), 6.07 (s, 1H), 5.27 (d, J = 1.5 Hz, 1H), 4.04 (s, 3H), 3.51 (q, J = 7.1 Hz, 2H), 2.02 (d, J = 1.4 Hz, 3H), 1.41 (s, 6H), 1.33 (t, J = 7.0 Hz, 3H).

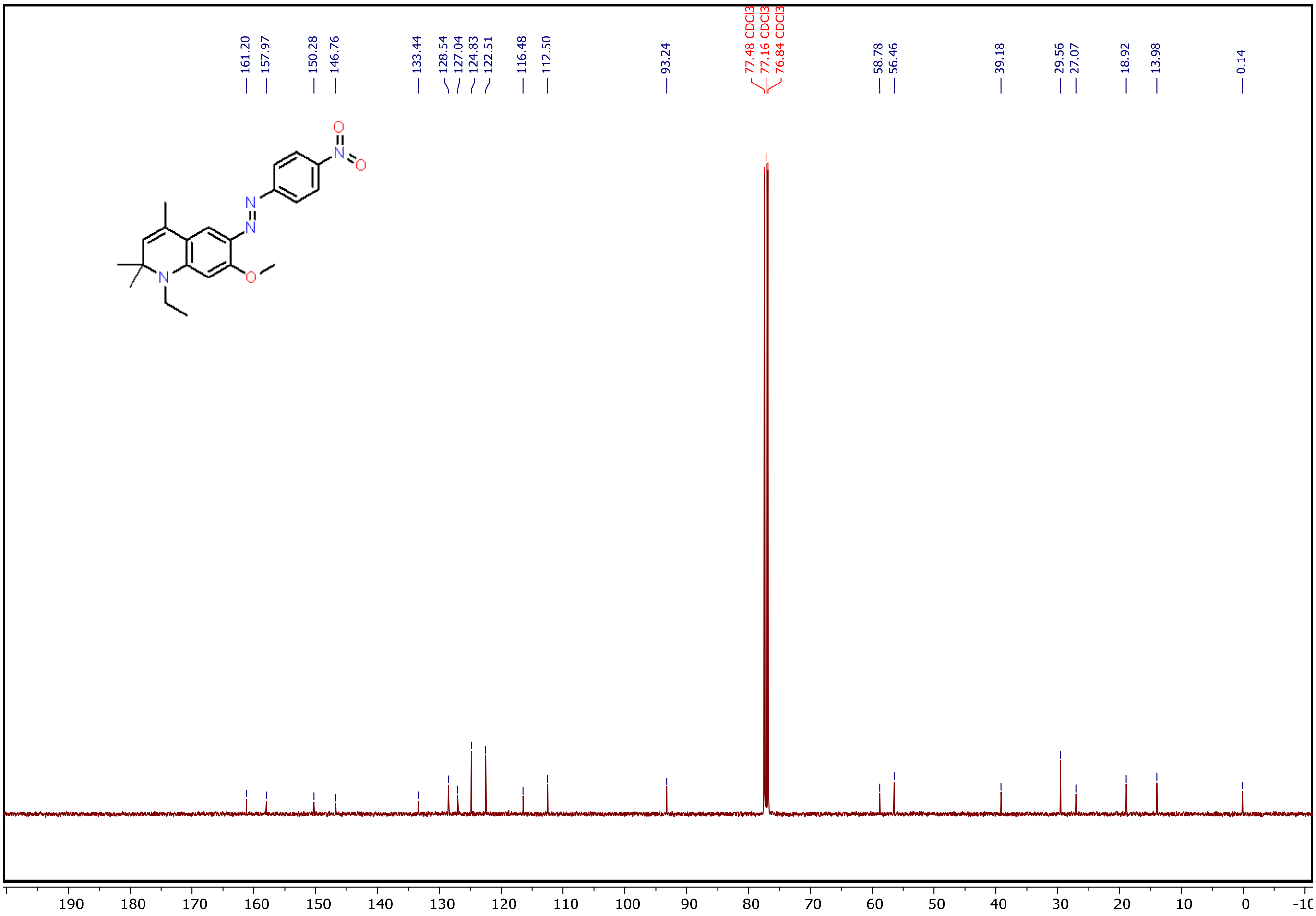

**Fig. S44.** $^{13}C$ spectrum of **13**.

$^{13}C$ NMR (101 MHz, $CDCl_3$) δ 161.20, 157.97, 150.28, 146.76, 133.44, 128.54, 127.04, 124.83, 122.51, 116.48, 112.50, 93.24, 77.48, 77.16, 76.84, 58.78, 56.46, 39.18, 29.56, 27.07, 18.92, 13.98, 0.14.

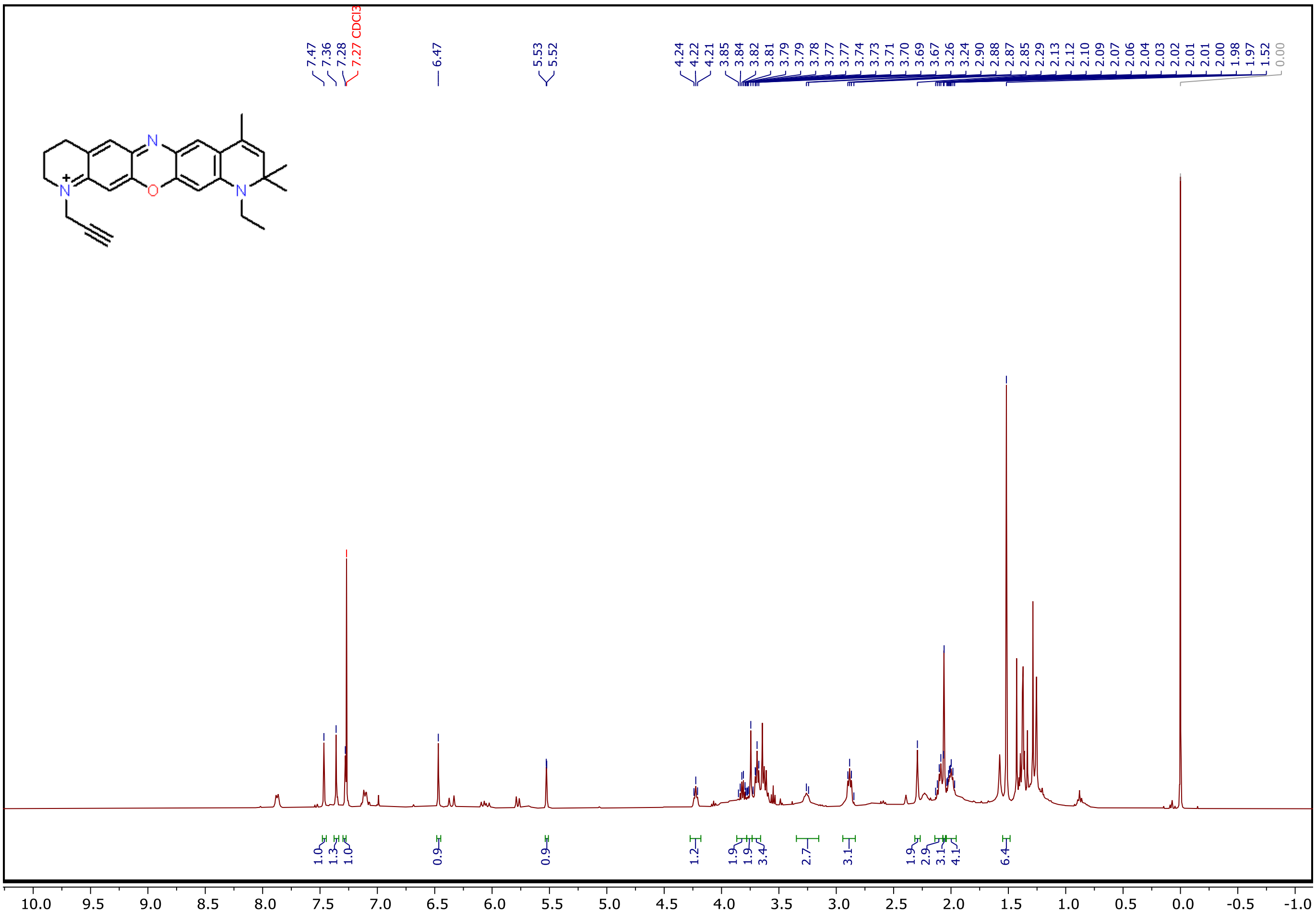

**Fig. S45.** $^1$H spectrum of **14**.

$^1$H NMR (400 MHz, CDCl3) δ 7.47 (s, 1H), 7.36 (s, 1H), 7.28 (s, 1H), 6.47 (s, 1H), 5.53 (d, J = 1.5 Hz, 1H), 4.27 – 4.18 (m, 1H), 3.82 (q, J = 5.7 Hz, 2H), 3.74 (s, 2H), 3.73 – 3.66 (m, 3H), 3.25 (d, J = 7.5 Hz, 3H), 2.88 (t, J = 6.3 Hz, 3H), 2.29 (s, 2H), 2.14 – 2.07 (m, 3H), 2.06 (s, 3H), 2.04 – 1.95 (m, 4H), 1.52 (s, 6H).
